# Community evenness and sample size affect estimates of predation intensity and prey selection: A model-based validation

**DOI:** 10.1101/2022.07.18.500550

**Authors:** Madhura Bhattacherjee, Devapriya Chattopadhyay

## Abstract

Predation estimates inferred from the preserved records of predation traces are essential in evaluating the evolutionary effect of ecological interactions. It is, however, crucial to establish how sampling intensity and community composition of an assemblage influence the reliability of these measures.

Using a resampling technique, we evaluated the effect of sampling intensity and a community’s evenness on the inferred predation estimates. We theoretically simulated model communities representing different levels of evenness, predation intensity, and predatory behavior (selective, non-selective). We calculated the total predation intensity and the number of prey species for each community. We then resampled each community without replacement and noted variations in the inferred measure from the accurate estimate as the sampling intensity increased. Our results demonstrate that the evenness of a community does not influence the inferred predation intensity for non-selective predation. However, communities with highly selective predation are sensitive to evenness and sampling intensity; inferred predation intensity of these assemblages can substantially differ from the actual value. The inferred number of prey species is also influenced by the community’s original evenness, predation selectivity, and predation intensity. When predation is selective, sampling intensity heavily influences communities with low evenness and low predation intensity; inferred predation intensity is underrepresented in smaller sample sizes. For communities of low evenness and predation intensity where rare species are attacked preferentially, the inferred prey richness differs significantly at a small sample size.

We proposed a post-facto standardization method for comparing predation estimates of discrete communities that differ in the sample size. We validated its utility using the published predation data of the Plio-Pleistocene molluscan fossil assemblage. The present approach attempts to provide critical insight into the reliability of predation estimates and may help in comparing predation patterns across time and space. Several factors, including preservation bias, might impact the final predation signature of an assemblage. It warrants a future research direction to develop a comprehensive framework of post-hoc standardization of assemblages with differing predation styles and preservation history.

## Introduction

The role of predation in shaping the marine ecosystems through time has been a common theme of study (Vermeij 1977; Vermeij et al. 1981; Signor and Brett 1984; Langerhans 2007; Stanley 2008; Barnes et al. 2010; Gorzelak et al. 2012; Kotta et al. 2018; Petsios et al. 2021). The relationship between the prey and predator is complex in theoretical terms posing a challenge in predicting the evolutionary outcome of predation (DeAngelis et al.1975; Berryman 1992; Haque 2012; Abrams 2015). For evaluating the evolutionary effects of predation, researchers rely on the deep time record of predation (Kitchell and Kitchell 1980; Vermeij et al. 1981; Kelley and Hansen 1993; Vermeij 1993; McNamara 1994; Kowalewski et al. 2005; Huntley and Kowalewski 2007; Baumiller et al. 2010; Klompmaker et al. 2017; Bicknell and Paterson 2018). Therefore, the accurate estimation of predation measures is of primary importance to studies of predator-prey systems.

For establishing predation events and inferring predation intensities, ecological studies use direct observations or indirect measures such as compositional characterization of digested food and fecal matter (Nilsen et al. 2012; Pringle et al. 2019). Although it is possible to recover direct observational evidence of predation events in past ecosystems by studying “caught-in-the-act” occurrences (Ehret et al. 2009; Ebert et al. 2015), paleoecological studies primarily rely on preserved predation traces, such as drill holes and repair scars (DeAngelis et al. 1985; Kelley and Hansen 1993; Dietl and Alexander 2000; Dietl et al. 2004; Alexander and Dietl 2005; Klompmaker and Kelley 2015). Based on the neontological experiments and field observations, complete drill holes and repair scars are interpreted as a successful attack by carnivorous gastropod (Carriker 1951; Kitchell et al. 1981; Kowalewski 2004; Hutchings and Herbert 2013; Chattopadhyay et al. 2014*a*; Mondal et al. 2014) and an unsuccessful predation attempt by durophagous predator respectively (Carriker 1951; Blundon and Kennedy 1982; Dietl and Alexander 2009). These traces recording the predation attempts on the prey’s hard shells, are some of the best quantifiable proxies for inferring predation from the fossil record (for review see (Alexander and Dietl 2003; Kelley and Hansen 2003; Klompmaker et al. 2019)). The frequency of repair scar (RF) and complete drill holes (DF) are used for evaluating various aspects of predation in deep time, including predation intensity and prey selection (Kitchell et al. 1981; Kelley and Hansen 1993; Kowalewski et al. 1998; Dietl 2003; Kase and Ishikawa 2003; Chattopadhyay and Baumiller 2010; Chattopadhyay and Dutta 2013; Tyler et al. 2013).

Inferences about interactions from predation traces have their limitations. The implicit assumption for such interpretation is that other processes do not alter the quantitative data provided by predation traces. It is recognized, however, that biases introduced through taphonomy may influence the biological reliability of these measures affecting the overall frequency of traces, site stereotypy, prey selection, and size selection (Roy et al. 1994; Nebelsick 1999; Zuschin et al. 2003; Kosloski 2011; Gorzelak et al. 2013; Chattopadhyay et al. 2014*b*; Chojnacki and Leighton 2014; Sime and Kelley 2016; Dyer et al. 2018; Pruden et al. 2018; Smith et al. 2019; Salamon et al. 2020). Apart from taphonomy, methods of collection and subsequent analyses may also influence the interpretation of predation patterns. In contrast to the bulk collection, targeted sampling of specific size class or taxon impacts inferred predation intensities (Kowalewski and Hoffmeister 2003; Kosloski et al. 2008; Ottens et al. 2012; Hattori et al. 2014; Chattopadhyay et al. 2016; Hausmann et al. 2018). Theoretical investigations also demonstrated the effect of sample size on inferred predation intensity (Smith et al. 2018, 2022). Analytical techniques to evaluate and compare predation measures across groups often impact the inferences (Kowalewski 2002; Leighton 2002; Grey et al. 2006; Stafford and Leighton 2011; Dietl and Kosloski 2013; Smith et al. 2018; Budd and Mann 2019).

Aspects of a specific community, such as evenness, selectivity of predation, and sampling intensity, may influence predation inferences drawn at the community level, such as predation intensity, prey selection. Such influences are crucial for studies that combine predation data from discrete samples and reconstruct temporal/spatial changes in predation patterns. Using theoretical simulation based on a resampling technique, we develop a methodological framework to understand the effect of community evenness, sampling intensity, and the nature of predation selectivity on inferred predation estimates. We attempt to estimate these effects on the inferred predation intensity and the number of prey species. The inferred number of prey species provides an insight into the choice of prey by the predator. We also propose a method of post-facto standardization and validate our approach using predation data from four Plio-Pleistocene fossil assemblages of Florida.

## Materials and method

We created several hypothetical live assemblages of molluscs that a specific group of predators attacks with differing probabilities. We use a resampling method to compare the predation patterns inferred from these assemblages. We assumed that all individuals are finally represented in the death assemblage, each predation attempt leaves a distinct mark on the prey, and all predators demonstrate the same prey-selection behavior in specific situations. We acknowledge that some of the particular values of species richness, evenness, predation intensity, and predation selectivity might be rare in nature. Our attempt, however, is to design and test a general framework applicable to a large spectrum of community structures and predation patterns, even if some end-member scenarios do not have a natural representation. It is also true that predation patterns observed in fossil assemblages may differ from that of the death assemblage due to taphonomic factors which have not been considered in the present study.

### Model assemblages

We created 30 hypothetical model assemblages, each with 30 species and 3000 individuals with varying evenness, predation intensity, and prey preferences (Table 1). Each model assemblage had a unique combination of evenness, predation intensity, and prey preference. To evaluate evenness, we used Pielou’s evenness index, one of the commonly used measures of evenness. We calculated the evenness of an assemblage (E_T_) as

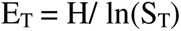

Where,

H = Shannon’s diversity index
S_T_ = Total number of species in the assemblage

**Table 1.** A summary of the model assemblages used for this study with varying evenness, predation intensity and predator preference.

| Evenness | Structure | PI<sub>T</sub> for Case 1<br>(Preys of all species are attacked with a probability of 0.2, 0.5 and 0.8 for low, medium and high PI<sub>prey</sub>, respectively) |  |  | PI<sub>T</sub> for Case 2<br>(Preys of only common species are attacked with a probability of 0.2, 0.5 and 0.8 for low, medium and high PI<sub>prey</sub> respectively) |  |  | PI<sub>T</sub> for Case 3<br>(Preys of only rare species are attacked with a probability of 0.2, 0.5 and 0.8 for low, medium and high PI<sub>prey</sub> respectively) |  |  |
| --- | --- | --- | --- | --- | --- | --- | --- | --- | --- | --- |
|  |  | Low | Medium | High | Low | Medium | High | Low | Medium | High |
| E<sub>T</sub> = 0.2 | N<sub>(S=1:29)</sub>=10, N<sub>(S=30)</sub>=2710<br>[1*2710 + 29*10] = 3000 | 0.2 | 0.5 | 0.8 | 0.18 | 0.45 | 0.72 | 0.02 | 0.05 | 0.08 |
| E<sub>T</sub> = 0.5 | N<sub>(S=1:3)</sub>=910, N<sub>(S=4:30)</sub>=10<br>[3*910 + 27*10] = 3000 | 0.2 | 0.5 | 0.8 | 0.18 | 0.46 | 0.73 | 0.02 | 0.05 | 0.07 |
| E<sub>T</sub> = 0.7 | N<sub>(S=1:5)</sub>=500, N<sub>(S=6:30)</sub>=20<br>[5*500 + 25*20] = 3000 | 0.2 | 0.5 | 0.8 | 0.17 | 0.42 | 0.67 | 0.03 | 0.08 | 0.13 |
| E<sub>T</sub> = 1 | N<sub>(S=1:30)</sub>=100<br>[30*100] = 3000 | 0.2 | 0.5 | 0.8 | NA | NA | NA | NA | NA | NA |

The evenness in these models ranged from a theoretical minimum of 0.1 to a theoretical maximum of 1. Model assemblages with maximum evenness of one had 100 individuals for 30 species. Assemblages with intermediate evenness of 0.7 had five common species with 500 individuals each and 25 rare species with 20 individuals each (Table 1). Assemblages with low evenness of 0.5 had 910 individuals in each of the three common species and ten individuals in each of the 27 rare species. For assemblages with a very low evenness of 0.2, there are only one common species with 2710 individuals, and the remaining 29 rare species consist of 10 individuals each.

We calculated the predation intensity at the assemblage level (PI_T_) and for prey species (PI_prey_). The total number of prey species is S_prey_. PI_T_ is calculated as

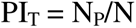

Where,

N_P_ = Number of individuals with predation mark
N = Total number of individuals in the assemblage

PI_prey_ denotes predation intensity in the species that have been attacked. The predation intensity of the total assemblage (PI_T_) was categorized into three levels: low (0.2), medium (0.5), and high (0.8) (Table 1). A certain number of individuals from specific species would be considered prey with predation marks as dictated by the (PI_T_). The prey-preference of the predator can either be non-selective or selective. In the case of non-selective predation (Case 1), all species have an equal probability of being attacked irrespective of their abundance (Fig 1). Selective predation represents assemblages where prey species have an unequal chance of being attacked relative to their abundance. In model assemblages with selective predation, we constructed two cases; the predator can either attack the common species (Case 2) (Fig 1) or the rare species (Case 3) (Fig 1).

**Figure 1.**
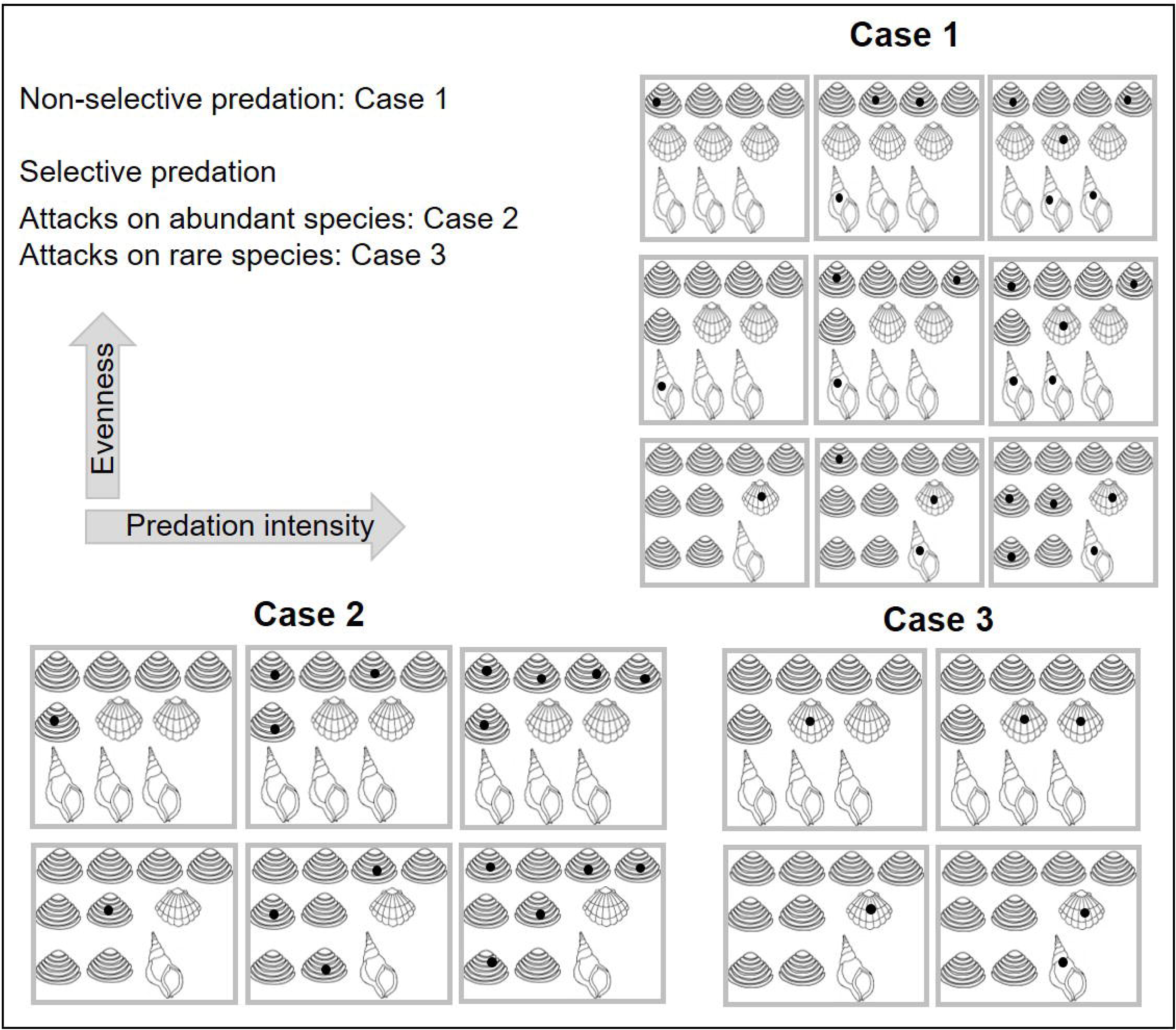
An illustrative diagram of model assemblages with varying degrees of evenness, predation intensity, and predation style (selective and non-selective). Mollusc drawings are from publicdomainpictures.net with subsequent modifications. [Figure 1. Two-column; Grayscale]

For all models, the probability of an attack is determined by the PI_T_ which can be 0.2, 0.5 or 0.8. For scenarios of selective predation, only certain species are available as prey and we assign the probability of attack as 0 to the rest of the species. In the case of selective predation on abundant species with low predation intensity, for instance, the probability of an attack is assigned as 0.2 for all the individuals of common species and 0.0 for all the individuals of rare species. Selective predation has not been considered for assemblages with maximum evenness because all the species are equally abundant and share an equal probability of attack.

### Simulation design

We performed a simulation to evaluate the effect of sample size on inferred predation intensity (PI_T.inf_) and the number of prey species (S_prey.inf_) for all the model assemblages. In the simulation, 100 individuals were drawn randomly from a model assemblage. The number of attacked individuals (N_P_) and the number of prey species (S_prey_) represented by the attacked individuals were counted in those 100 individuals. Inferred predation intensity (PI_T.inf_) for the drawn sample is calculated as a ratio of the number of attacked individuals and the total number of individuals (i.e., 100 in the first draw). We kept the step size as 100 to gain an accurate representation of predation intensity and to avoid the issues related to insufficient sample size (Kosloski et al. 2008; Dietl and Kosloski 2013; Smith et al. 2022). The exact process is repeated 30 times without replacement until all the individuals from the assemblage are sampled. Following the principles of rarefaction analysis that are helpful when attempting to standardize sampling effort, we chose to use subsampling without replacement (Kowalewski and Novack-Gottshall 2010). This entire process was iterated 1000 times. The mean and standard deviation are calculated for inferred predation intensity (PI_T.inf_) and prey species richness (S_prey.inf_) over 1000 iterations for a specific model assemblage. The difference in predation intensity (Diff_PI_) is calculated as the difference between PI_T_ and PI_T.inf_ for an assemblage. Similarly, the difference between S_prey_ and S_prey.inf_ is taken as the difference of prey species richness (Diff_S_). The same technique is applied to all the model assemblages.

### Predation dataset

We used data on predation records of molluscs from four Pleistocene localities in Florida (Chattopadhyay and Baumiller 2010) to validate the proposed technique. The goal is to quantitatively evaluate if we can compare the predation estimates of discrete communities characterized by different evenness, predation style and sample size. The dataset consists of abundance, drilling frequency, and repair scar frequency of 14 molluscan species. We drew samples without replacement from each locality with increasing sample size. The sample size for each draw was a hundred until the last draw; in the last draw, where the remaining sample size is less than 200, all are drawn. For Punta Gorda (total=2418 individuals), 100 individuals were drawn 23 times, and 118 individuals were drawn for the last (24^th^) draw. A similar procedure is followed for Miami Canal (total = 4794 individuals), Mc Queens pit (total=659 individuals), and Chiquita (total=894 individuals).

We used a sampling standardization protocol to compare these assemblages and assess the sensitivity of the inferred predation intensity (PI_T.inf_) and inferred prey-species richness (S_prey.inf_) on sampling intensity. The sample size of Mc Queens pit (659) is considered a reference as it has the smallest sample size among all four locations. The distribution of inferred predation intensity (PI_T_) is compared for all assemblages at a sample size of 500 by a pairwise comparison using Kolmogorov-Smirnov (K-S) test. If the pairwise K-S test shows significant differences between all pairs of assemblages, then the variation between assemblages is not caused by sampling and community evenness. If some assemblages show non-significant differences in the pairwise K-S test, then small sample size might influence inferred predation intensity and prey richness. Larger sample size is considered a reference, and the pairwise comparison using the K-S test is repeated again for those pairs of assemblage. The process is repeated until the maximum number of assemblage pairs shows significant differences. Following a similar protocol, the distribution of inferred prey-species richness (S_prey.inf_) is also compared.

All simulations and statistical analyses were performed in R (version 4.2.0) (R Core Development Team, 2012).

## Results

### Inferred predation intensity

The inferred predation intensity (PI_T.inf_) may vary substantially from the actual value of overall predation intensity (PI_T_) and predation intensity of prey groups (PI_prey_), especially at smaller sample sizes (Fig 2). For non-selective predation (Case 1), Diff_PI_ is affected by the sample size, although not by evenness. At a smaller sample size, the difference is higher (Diff_PI=_0.04) implying a lower PI_T.inf_ than the actual PI_T_ value. PI_T.inf_ converges to PI_T_ with increasing sample size (Fig 3, Table 5).

**Figure 2.**
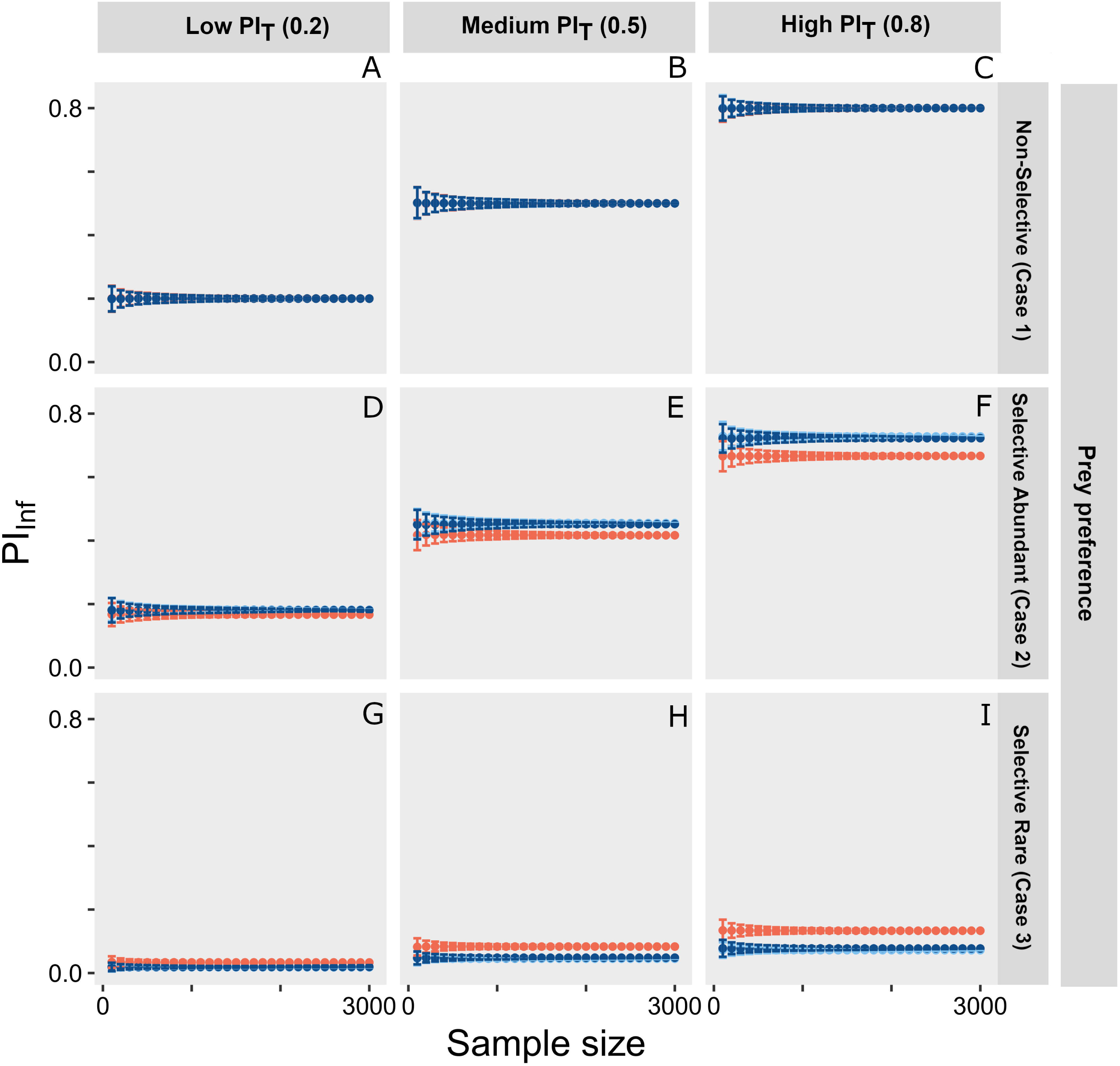
Plot showing variation in inferred predation intensity (PI_T.inf_) with varying sample sizes for different model assemblages. The rows indicate the different degrees of the selectiveness of predation, and the columns indicate predation intensity in the original assemblage (PI_T_). The warmer colors represent assemblages with higher evenness. [Figure 2. Two columns; Color]

**Figure 3.**
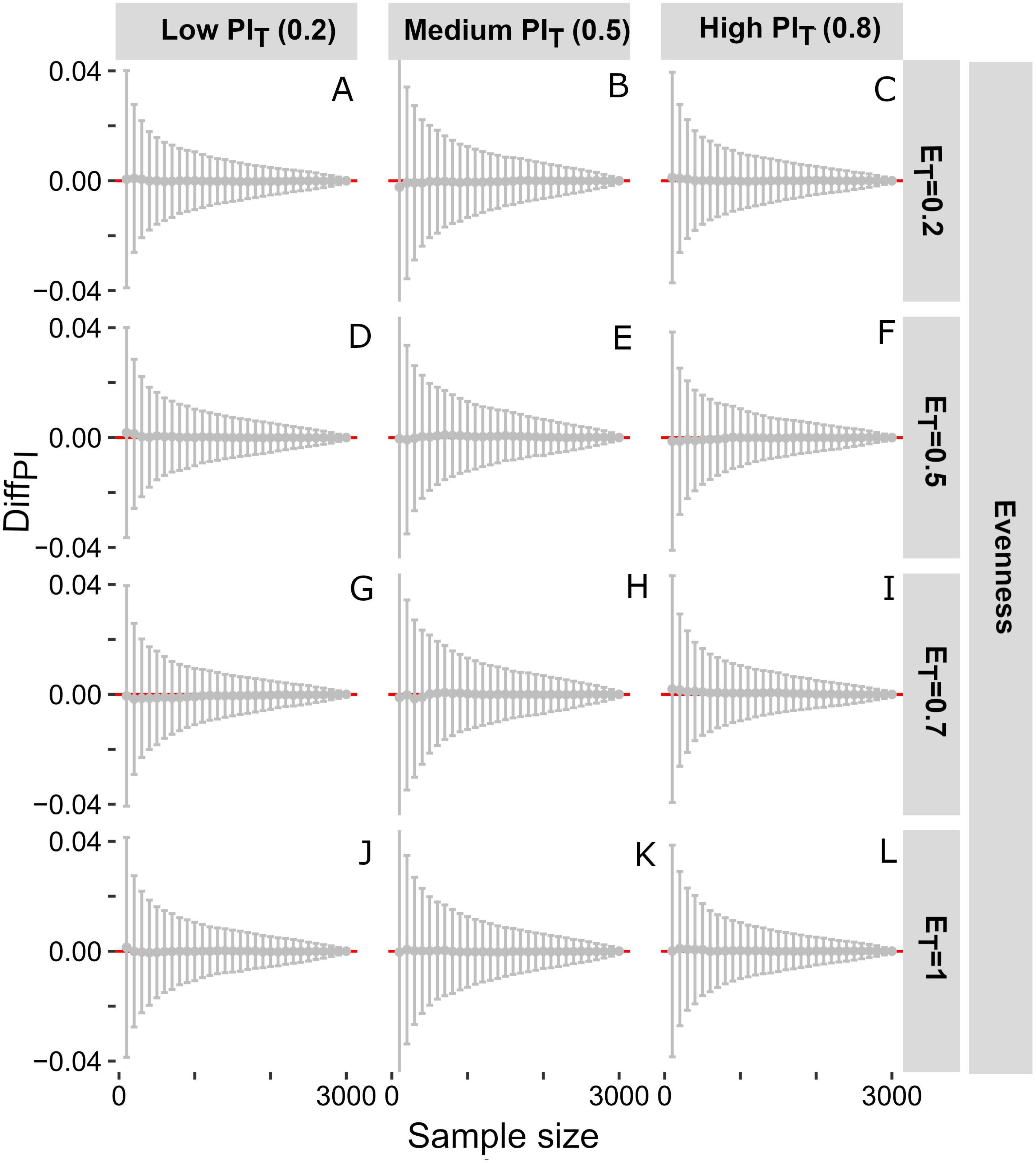
Plot showing the difference between the original (PI_T_) and inferred predation intensity (PI_T.inf_) at varying sample size for non-selective predation (Case 1). The rows indicate evenness and the columns represent original predation intensity. The red line represents the zero line where overall and inferred predation intensities are the same (PI_T.inf_ = PI_T_). The grey dots and bars represent the mean and standard deviation of the simulated differences for specific model assemblages. [Figure 3. Two columns; Color]

**Table 2.**
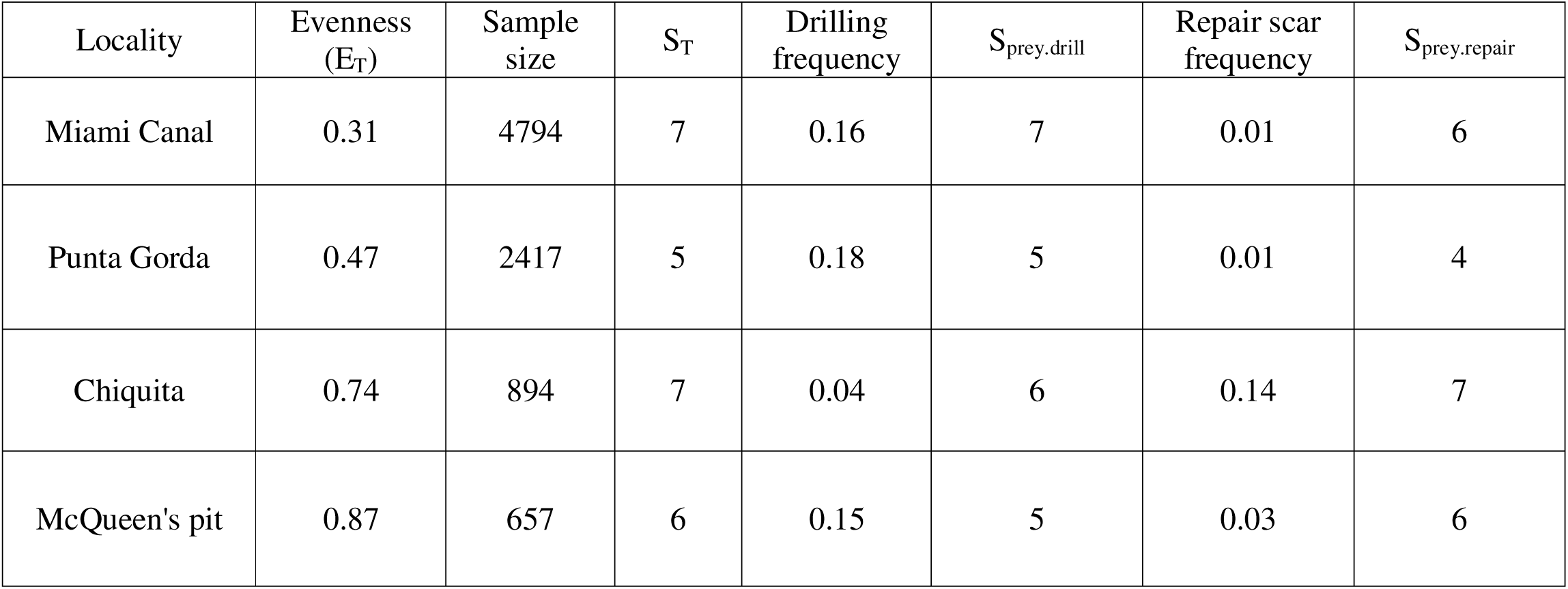
A summary of the predation data from four Plio-Pleistocene fossil assemblages of Florida.

| Locality | Evenness<br>($E_T$) | Sample<br>size | $S_T$ | Drilling<br>frequency | $S_{\text{prey.drill}}$ | Repair scar<br>frequency | $S_{\text{prey.repair}}$ |
| --- | --- | --- | --- | --- | --- | --- | --- |
| Miami Canal | 0.31 | 4794 | 7 | 0.16 | 7 | 0.01 | 6 |
| Punta Gorda | 0.47 | 2417 | 5 | 0.18 | 5 | 0.01 | 4 |
| Chiquita | 0.74 | 894 | 7 | 0.04 | 6 | 0.14 | 7 |
| McQueen's pit | 0.87 | 657 | 6 | 0.15 | 5 | 0.03 | 6 |

**Table 3.** The result of Spearman rank order correlation test for proportional abundance and PI_prey_ for the predation estimates across four Plio-Pleistocene fossil assemblages of Florida (Chattopadhyay and Baumiller, 2010). The statistically significant (p<0.05) results are marked in bold.

| Predation | Location | rho | p | Inferred scenario |
| --- | --- | --- | --- | --- |
| Drilling | Punta Gorda | 0.87 | 0.05 | Case 2 |
|  | McQueen's pit | 0.83 | 0.06 | Case 1 |
|  | Chiquita | 0.68 | 0.08 | Case 1 |
|  | Miami canal | **0.99** | **<0.001** | - |
| Durophagy | Punta Gorda | 0.21 | 0.74 | Case 1 |
|  | McQueen's pit | 0.46 | 0.35 | Case 1 |
|  | Chiquita | 0.24 | 0.61 | Case 1 |
|  | Miami canal | **0.79** | **0.03** | Case 2 |

**Table 4.** The test-statistic (D) of Kolmogorov–Smirnov test comparing the predation estimates across four Plio-Pleistocene fossil assemblages of Florida using sample-standardization protocol. All the results are statistically significant (p<0.05).

| Estimate | Predation | Location | McQueen's pit | Chiquita | Miami Canal |
| --- | --- | --- | --- | --- | --- |
| Predation intensity | Drilling | Punta Gorda | **0.8** | **1** | **0.53** |
|  |  | McQueen's pit |  | **0.24** | **0.31** |
|  |  | Chiquita |  |  | **1** |
|  | Durophagy | Punta Gorda | **0.9** | **1** | **0.19** |
|  |  | McQueen's pit |  | **0.96** | **0.85** |
|  |  | Chiquita |  |  | **1** |
| Prey species richness | Drilling | Punta Gorda | **0.38** | **0.31** | **0.13** |
|  |  | McQueen's pit |  | **0.24** | **0.36** |
|  |  | Chiquita |  |  | **0.29** |
|  | Durophagy | Punta Gorda | **0.84** | **1** | **0.41** |
|  |  | McQueen's pit |  | **1** | **0.95** |
|  |  | Chiquita |  |  | **1** |

**Table 5.** A summary of the difference in inferred predation intensity from the original value for the model assemblages. Each cell contains information about the mean value and standard deviation of Diff_PI_; the first two represents the sign and magnitude of the mean value. A positive mean value of Diff_PI_ indicates a larger value of original than inferred predation intensity (PI_T_ > PI_T.inf_).

| Evenness | $\text{Diff}_{\text{PI}}$ for Case1 | | | $\text{Diff}_{\text{PI}}$ for Case2 | | | $\text{Diff}_{\text{PI}}$ for Case3 | | |
| --- | --- | --- | --- | --- | --- | --- | --- | --- | --- |
|  | Low | Medium | High | Low | Medium | High | Low | Medium | High |
| $E_{\text{T}} = 0.2$ | +ve,<br><0.001,<br>0.013 | +ve,<br><0.001,<br>0.016 | +ve,<br><0.001,<br>0.013 | -ve,<br><0.001,<br>0.012 | -ve,<br>0.001,<br>0.015 | -ve,<br>0.003,<br>0.014 | +ve,<br><0.001,<br>0.004 | +ve,<br>0.002,<br>0.006 | +ve,<br>0.002,<br>0.008 |
| $E_{\text{T}} = 0.5$ | +ve,<br><0.001,<br>0.013 | +ve,<br><0.001,<br>0.016 | +ve,<br><0.001,<br>0.013 | -ve,<br>0.002,<br>0.012 | +ve,<br>0.005,<br>0.015 | +ve,<br>0.002,<br>0.014 | +ve,<br>0.002,<br>0.004 | +ve,<br>0.005,<br>0.006 | -ve,<br>0.002,<br>0.008 |
| $E_{\text{T}} = 0.7$ | +ve,<br><0.001,<br>0.012 | -ve,<br><0.001,<br>0.015 | -ve,<br><0.001,<br>0.012 | +ve,<br>0.003,<br>0.012 | +ve,<br>0.004,<br>0.016 | +ve,<br>0.003,<br>0.016 | -ve,<br>0.003,<br>0.006 | -ve,<br>0.003,<br>0.009 | -ve,<br>0.003,<br>0.011 |
| $E_{\text{T}} = 1$ | -ve,<br><0.001,<br>0.013 | +ve,<br><0.001,<br>0.015 | +ve,<br><0.001,<br>0.013 | NA | NA | NA | NA | NA | NA |

Evenness influences inferred predation intensity (PI_T.inf_) when the predation is non-selective (Case 2 and 3) (Fig 2). When the common species are preferentially attacked (Case 2), Diff_PI_ is low (mean = -0.0003, standard deviation = 0.0124) for communities with lower evenness (E_T_=0.2) and low original predation intensity (PI_T_ =0.2) implying good correspondence between PI_T.inf_ and PI_T_ (Fig 4, Table 5). Communities with higher evenness (E_T_>0.2) showed high Diff_PI_ (Table 5) even at a higher sample size implying that PI_T.inf_ will be different from PI_T_ (Fig 4). Except for one specific model assemblage (E_T_=0.5, PI_T_ =0.2), all assemblages show a lower PI_T.inf_ than the original PI_T_ (Table 5).

**Figure 4.**
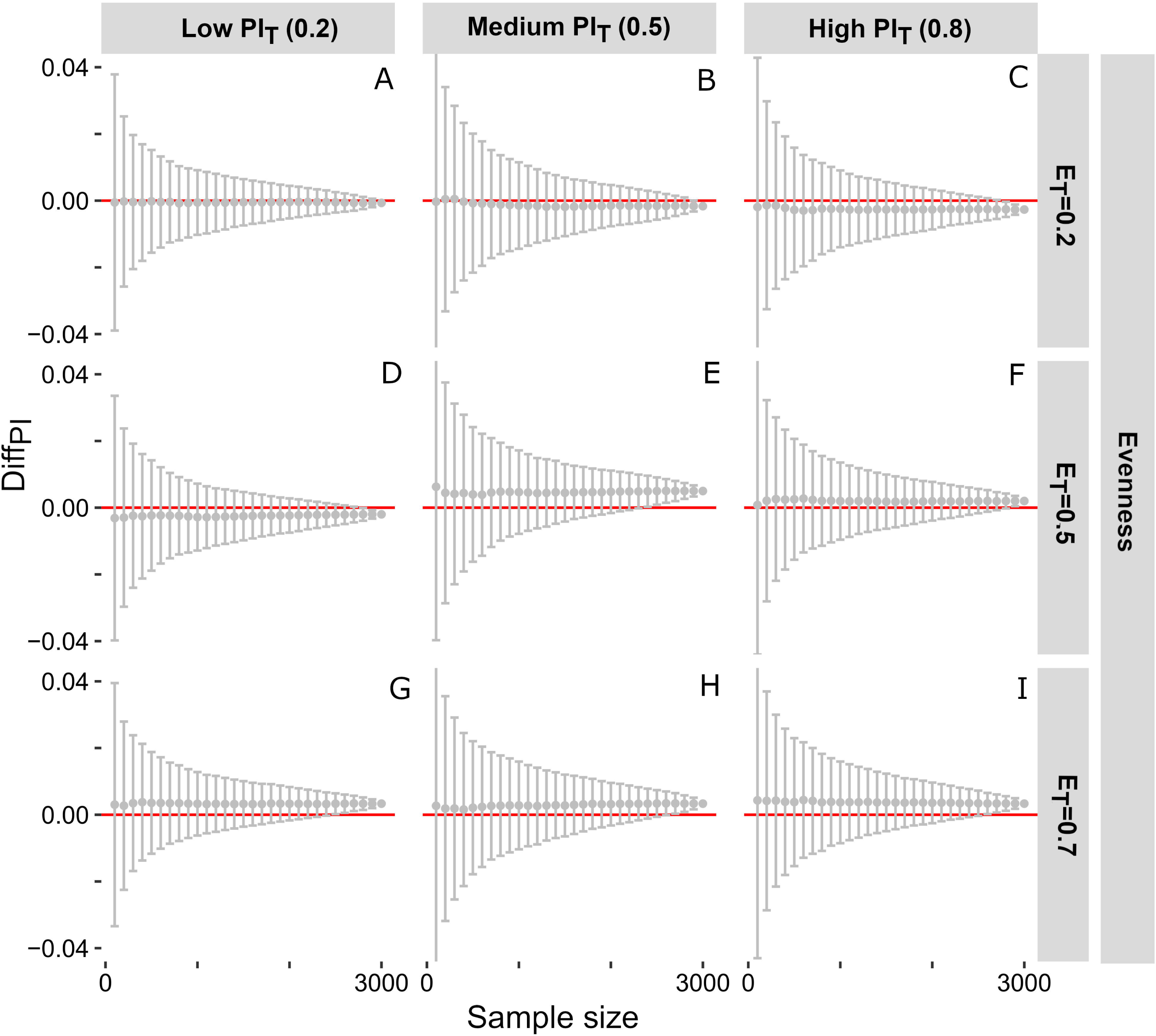
Plot showing the difference between the original (PI_T_) and inferred predation intensity (PI_T.inf_) at varying sample size for selective predation of the common species (Case 2). The rows indicate evenness and the columns represent original predation intensity. The red line represents the zero line where overall and inferred predation intensities are the same (PI_T.inf_ = PI_T_). The grey dots and bars represent the mean and standard deviation of the simulated differences for specific model assemblages. [Figure 4. Two columns; Color]

When rare species are attacked (Case 3), the Diff_PI_ varies depending on the combination of evenness and predation intensity. The Diff_PI_ is positive for all communities with low evenness (E_T_=0.2) irrespective of the predation intensity (Fig 5A-C, Table 5), implying, a lower value of PI_T.inf_ compared to PI_T_. Communities with high evenness (E_T_=0.7) showed negative Diff_PI_, indicating a higher PI_T.inf_ than PI_T_ (Fig 5G-I). The Diff_PI_ value in communities with medium evenness (E_T_=0.5) depends on predation intensity; in those communities, PI_T.inf_ is lower compared to PI_T_ for lower predation intensity (PI_T_ < 0.8) (Fig 5D-E) and higher in for higher predation intensity (Fig 5F, Table 5). However, the variation in Diff_PI_ is lower for Case 3 compared to similar communities in Case 2 (Table 5).

**Figure 5.**
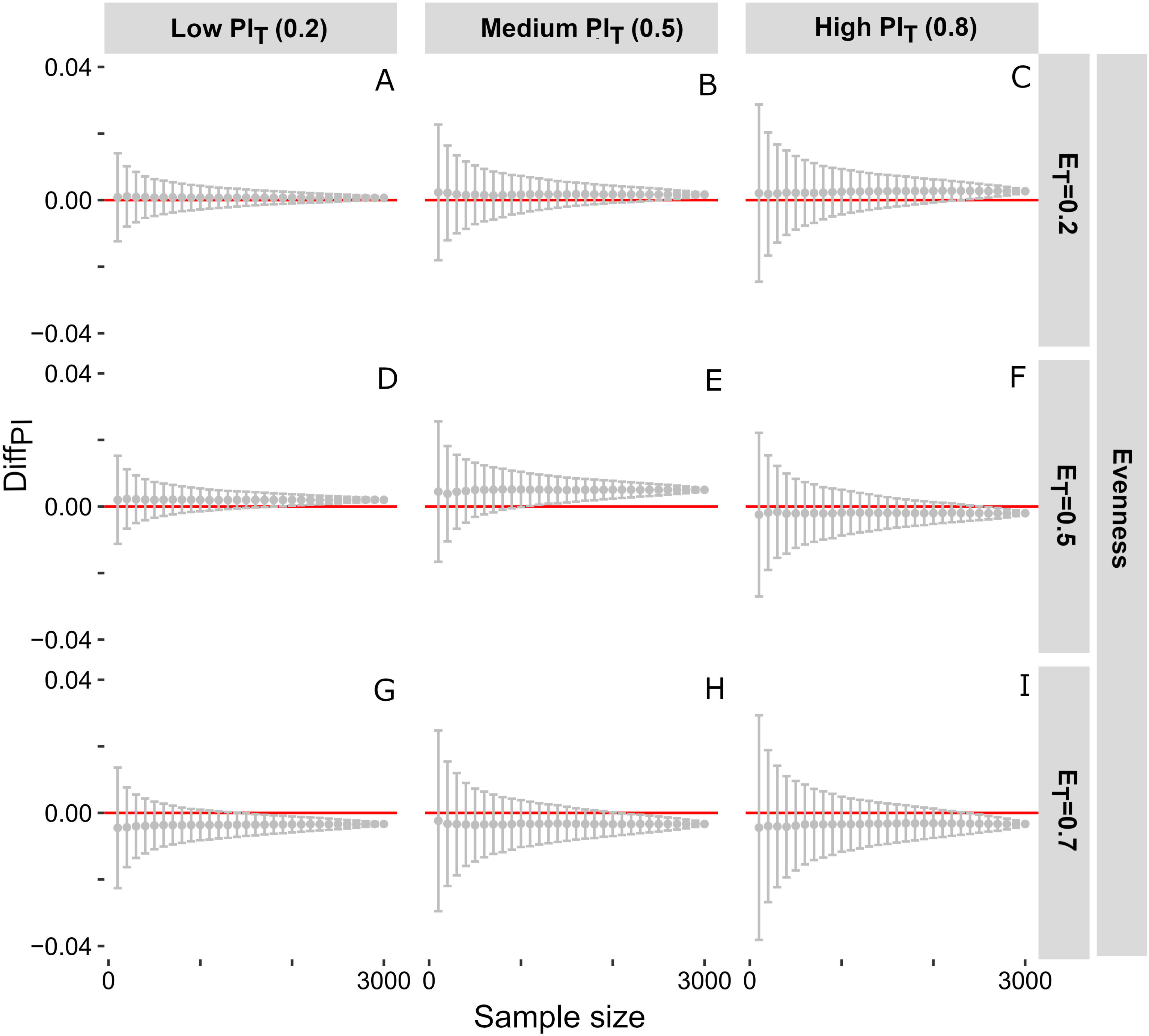
Plot showing the difference between the original (PI_T_) and inferred predation intensity (PI_T.inf_) at varying sample size for selective predation of the rare species (Case 3). The rows indicate evenness and the columns represent original predation intensity. The red line represents the zero line where overall and inferred predation intensities are the same (PI_T.inf_ = PI_T_). The grey dots and bars represent the mean and standard deviation of the simulated differences for specific model assemblages. [Figure 5. Two columns; Color]

### Inferred number of prey species

The inferred number of prey species (S_prey.inf_) follows a rarefaction curve where S_prey.inf_ increases with increasing sample size before plateauing and converging to the actual value of S_prey_ (Fig 6). The required sample size for convergence depends on evenness and selectivity of predation. In the case of non-selective predation (Case 1), the Diff_S_ decreases exponentially with increasing sample size and the converges to zero at a range of sample sizes depending on the evenness and predation intensity (Fig 7, Table 6). At a given predation intensity, convergence occurs at a smaller sample size with increasing evenness. For example, at low predation intensity (PI_T_=0.2), the required sample size for Diff_S_ to converge to 0 is 3000 when evenness is low (E_T_=0.2) (Fig 7A) and 1200 when evenness is high (E_T_=1) (Fig 7J, Table 6).

**Figure 6.**
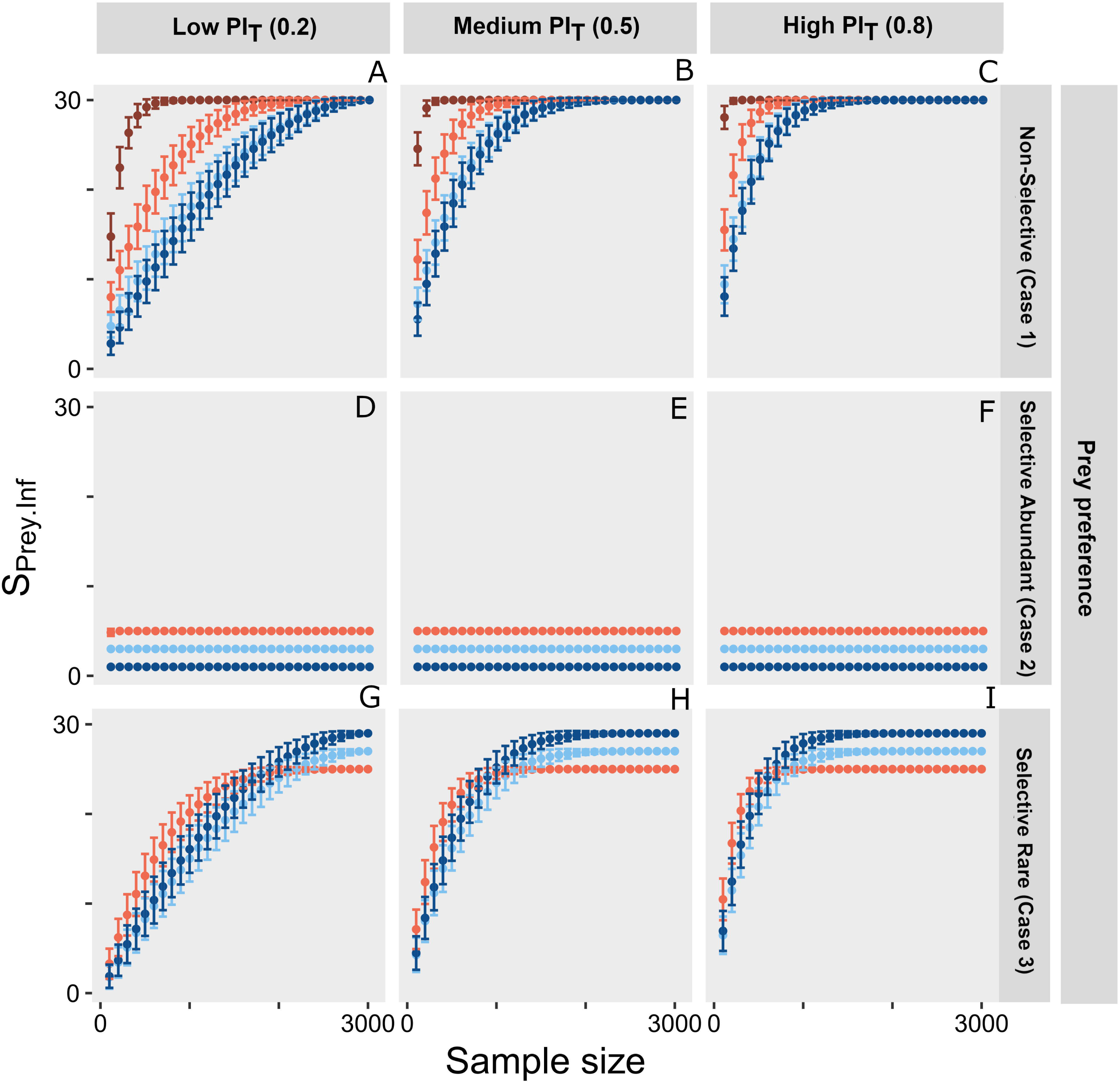
Plot showing variation in inferred prey species richness (S_prey.inf_) with varying sample sizes for different model assemblages. The rows indicate the different degrees of the selectiveness of predation, and the columns indicate prey species richness in the original assemblage (S_prey_). The warmer colors represent assemblages with higher evenness. [Figure 6. Two columns; Color]

**Figure 7.**
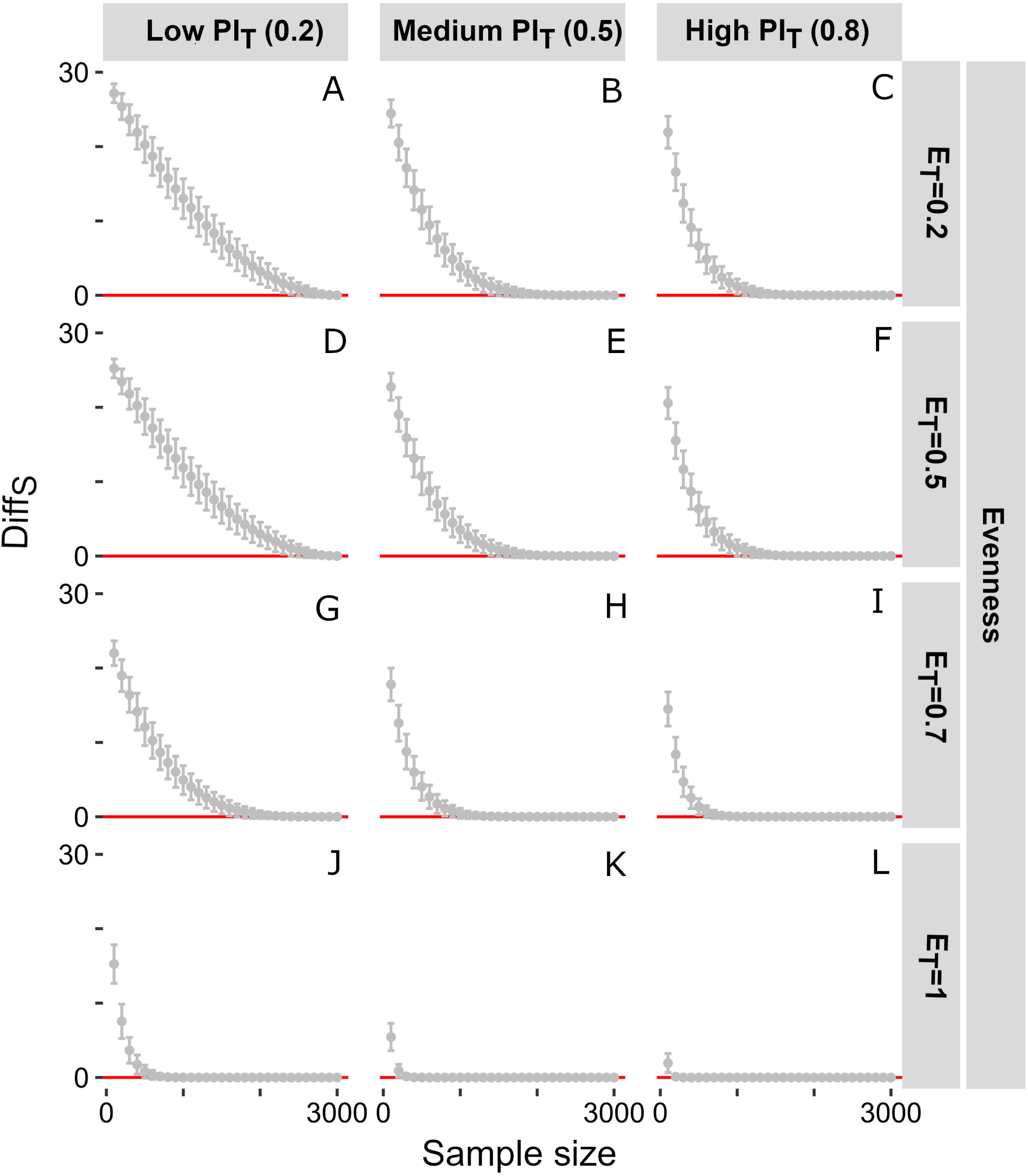
Plot showing the difference between the original (S_prey_) and prey species richness (S_prey.inf_) at varying sample size for non-selective predation (Case 1). The rows indicate evenness and the columns represent original predation intensity. The red line represents the zero line where overall and inferred prey species richness are the same (S_prey.inf_ = S_prey_). The grey dots and bars represent the mean and standard deviation of the simulated differences for specific model assemblages [Figure 7. Two columns; Color]

**Table 6.** A summary of the difference in inferred prey species richness from the original value for the model assemblages. Each cell contains the minimum sample size required for Diff_S_ to converge to zero for each model assemblages. A smaller number indicates that the inferred prey species richness converges to the original value (S_prey.inf_ = S_prey_) at smaller sample size.

| Evenness | Required sample size for convergence of $\text{Diff}_S$ for Case 1 | | | Required sample size for convergence of $\text{Diff}_S$ for Case 2 | | | Required sample size for convergence of $\text{Diff}_S$ for Case 3 | | |
| --- | --- | --- | --- | --- | --- | --- | --- | --- | --- |
|  | Low | Medium | High | Low | Medium | High | Low | Medium | High |
| $E_T = 0.2$ | 3000 | 2800 | 2100 | 100 | 100 | 100 | 3000 | 2700 | 2200 |
| $E_T = 0.5$ | 3000 | 2700 | 2100 | 100 | 100 | 100 | 3000 | 2700 | 2200 |
| $E_T = 0.7$ | 2900 | 1900 | 1400 | 300 | 100 | 100 | 2800 | 1900 | 1400 |
| $E_T = 1$ | 1200 | 700 | 400 | NA | NA | NA | NA | NA | NA |

In selective predation when common species are preyed upon (Case 2), Diff_S_ does not reflect any sensitivity to the sample size (Fig 8). This is due to the low value of S_prey_ that converges to its actual value within the first few draws (Fig 8, Table 6). However, when the rare species are attacked (Case 3), S_prey.inf_ is highly sensitive to the sample size because the required sample size for convergence of Diff_S_ depends on evenness and predation intensity (Fig 9). In general, communities with higher evenness require small sample size for convergence of Diff_S_ implying less sensitivity of S_prey.inf_ on sample size at a given predation intensity (Fig 9). For example, at low predation intensity (PI_T_=0.2), the required sample size for Diff_S_ to converge to 0 is 3000 when evenness is low (E_T_=0.2) (Fig 9A) and 2800 when evenness is high (E_T_=0.7) (Fig 9G, Table 6).

**Figure 8.**
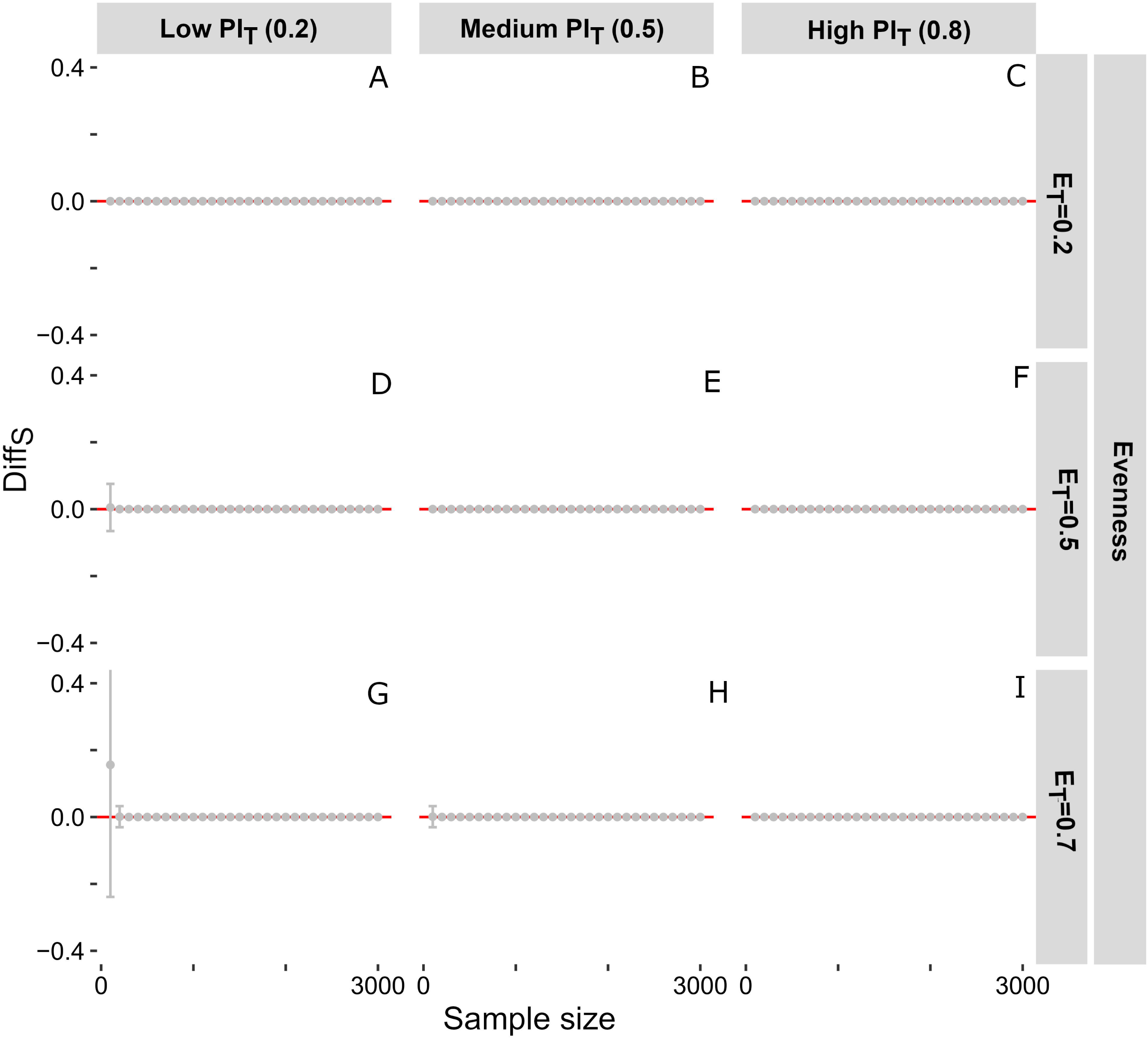
Plot showing the difference between the original (S_prey_) and prey species richness (S_prey.inf_) at varying sample size for selective predation of common species (Case 2). The rows indicate evenness and the columns represent original predation intensity. The red line represents the zero line where overall and inferred prey species richness are the same (S_prey.inf_ = S_prey_). The grey dots and bars represent the mean and standard deviation of the simulated differences for specific model assemblages [Figure 8. Two columns; Color]

**Figure 9.**
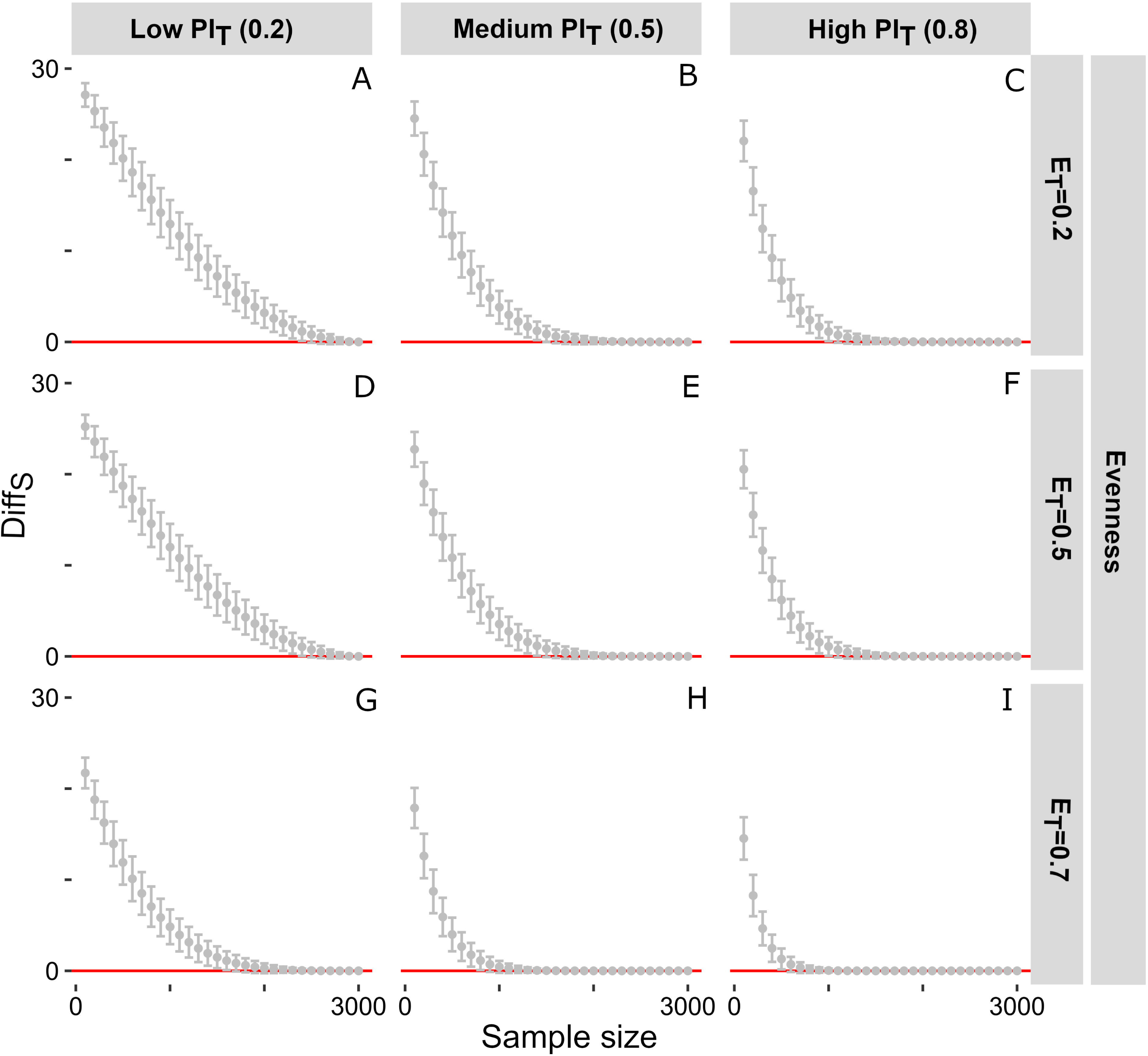
Plot showing the difference between the original (S_prey_) and prey species richness (S_prey.inf_) at varying sample size for selective predation of rare species (Case 3). The rows indicate evenness and the columns represent original predation intensity. The red line represents the zero line where overall and inferred prey species richness are the same (S_prey.inf_ = S_prey_). The grey dots and bars represent the mean and standard deviation of the simulated differences for specific model assemblages [Figure 9. Two columns; Color]

### Inferred predation estimates from Florida

The assemblages from the four localities of Florida are different in terms of their evenness and sample size (Table 2). Except for Miami Canal, all the localities show a case of non-selective predation represented by the lack of correlation between the relative abundance of the prey and prey-specific predation intensity (PI_prey_) for drilling and durophagous predation (Table 3). The significant positive correlation in Miami Canal implies that this is not non-selective predation (Case 1). Because the drilling predators attack all the species, it does not represent Case 2 or Case 3 for drilling predation. For durophagous predation where the most abundant species are attacked, it approximates Case 2.

For inferred predation intensity (PI_T.inf_), there is substantial overlap between three localities (Punta Gorda, Miami Canal, and Mc Queens pit) for both drilling and durophagy (Fig 10). For inferred prey species richness (S_prey.inf_), the assemblages show slightly different patterns between drilling and durophagous predation. For drilling predation, all the assemblages show a substantial overlap (Fig 10). The durophagous predation record, however, shows a separation between communities with low evenness (Punta Gorda) and high-evenness (Mc Queens pit, Chiquita) (Fig 10).

**Figure 10.**
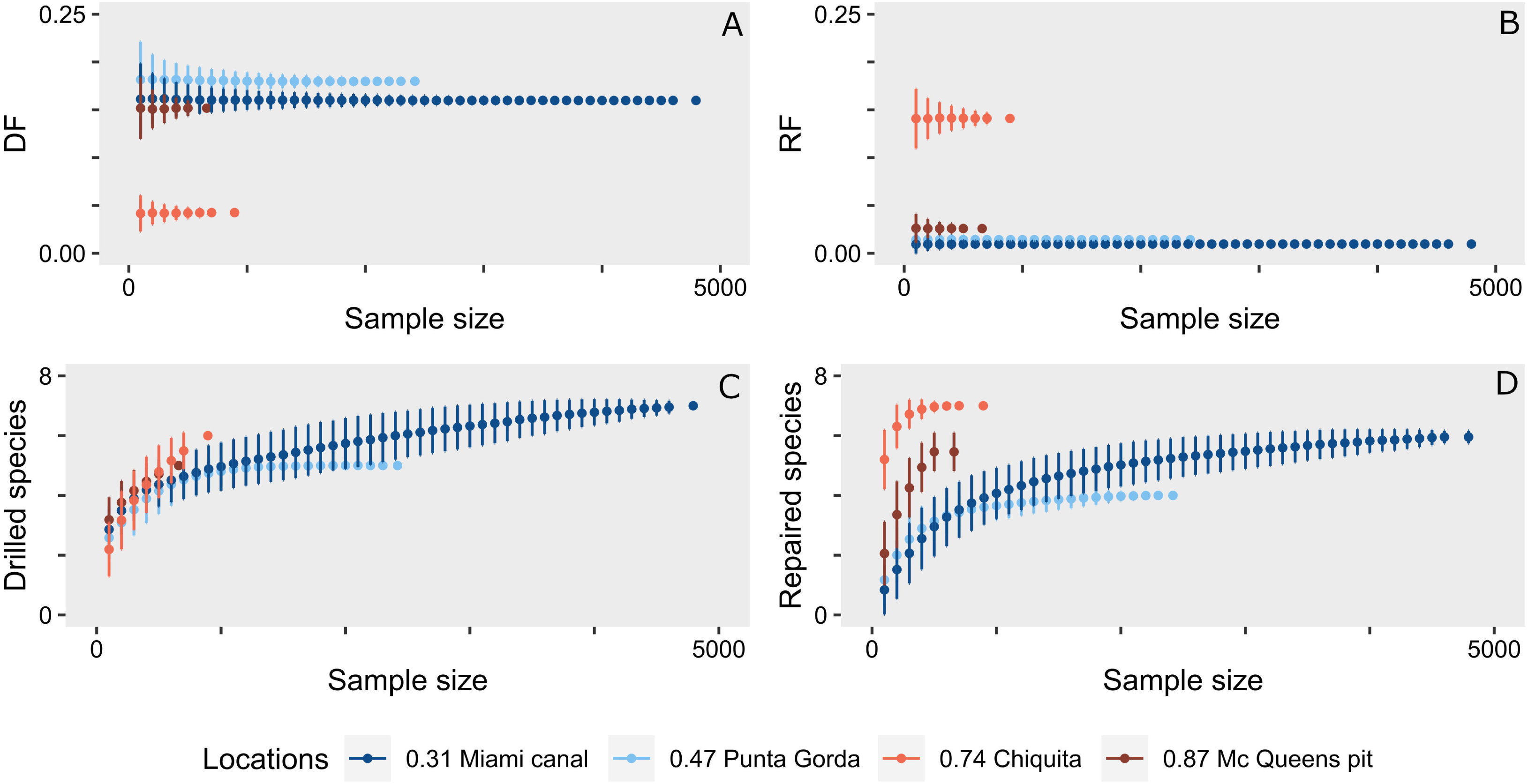
Plot showing variation in inferred estimates of drilling and durophagous predation with varying degrees of sampling for four Pleistocene molluscan assemblages of Florida with different evenness (E_T_). The top row represents the sample size variation in inferred predation intensity (PI_inf_). The bottom row shows the inferred number of prey species (S_prey.inf_) with varying sample sizes. The warmer colours represent assemblages of higher evenness. [Figure 10. Two columns; Color]

For inferred predation intensity (PI_T.inf_) and species richness (S_prey.inf_), the sample size-standardized resampling protocol (described before) shows a significant difference (*p* <0.005) in all pairwise comparison (Table 4). This implies that the difference across assemblages cannot be explained by the sampling intensity or the evenness of the assemblage.

## Discussion

Paleontological research on predation has expanded rapidly in scope, methods, and goals. In the recent years, many studies focused on documenting the evidence of predation from times, geographic areas, and taxa that are poorly known for their predation record (Rojas et al. 2014; Randle and Sansom 2019; Bicknell and Holland 2020; Gordillo and Malvé 2021; Klompmaker and Landman 2021; Gordillo et al. 2022) and using predation records for testing evolutionary hypotheses (Klompmaker et al. 2017; Gehling and Droser 2018; Harper et al. 2018; Lerosey-Aubril and Peel 2018; Petsios et al. 2021). In contrast, a relatively small number of studies focused on the analytical methods to evaluate the reliability of predation measures in recent years (Smith et al. 2018, 2019, 2022; Budd and Mann 2019). Our model demonstrates how the inferred predation intensities may vary with community structures, predation selectivity, and sampling intensities. It highlighted the importance of these factors in influencing the predation estimates of live and death/fossil assemblages; it also underscores why it is necessary to develop a methodological framework of sample standardization before comparing predation estimates of assemblages separated by time and space.

### Effect on the inferred intensity

Our simulation results show that communities’ evenness does not significantly change the inferred predation intensity when random encounters between predator and prey guide predation. However, it is uncommon to find predation events completely random in the natural world. Predators select prey species to maximize net energy gain within the constraints of many factors, including reproductive demands, predator interference, predation risk, avoidance of prey, deterrents, and predator behavior (Seitz et al. 2001; Stephens and Krebs 2019). The inferred predation intensity may differ significantly from the original predation intensity in such selective predation. Following the considerations of optimal foraging theory (Hughes 1980; Pyke 1984; Burrows and Hughes 1991; Stephens and Krebs 2019), two aspects make predation selective. The first is the relative ease with which a predator encounters prey. Encounter in the marine ecosystem is determined by several things, including the abundance of the prey, accessibility of the prey, landscape heterogeneity, predator abundance, abundance of secondary predators, and habitat type (Ryer and Olla 1995; Seitz et al. 2001; Sims et al. 2006; Casey and Chattopadhyay 2008; Martinelli et al. 2015). Keeping the other factors constant, the probability of encounter increases with the relative abundance of a prey species (Vermeij 1983; Leighton 2002; Leonard-Pingel and Jackson 2013); this decreases the foraging time and increases the net energy gain of the predator. The second aspect is the traits (morphological, ecological, behavioral) of the prey that dictate the net energy gain of the predator. The final selection by the predator is often a combination of these factors. A higher attack rate may be found in an abundant prey species due to its higher encounter rate than a rarer species. This would lead to scenarios similar to Case 2, where the inferred predation intensity of low-evenness communities would be higher than the actual predation intensity. This inflated measure results from the over-representation of common species in smaller samples that are primarily attacked.

Most often than not, the encounter frequency does not dictate the attack frequency, and the selection of prey is guided by the prey traits such as size, morphology, behavior (Kitchell et al. 1981; Palmqvist et al. 1996; Leighton 2001; Zlotnik and Ceranka 2005; Chattopadhyay and Dutta 2013; Chattopadhyay et al. 2014*a*, 2015, 2020; Martinelli et al. 2015; Chandroth and Chattopadhyay 2022). These would be similar to Case 3, where the most dominant groups are not preyed upon. The inferred predation intensity of low-evenness communities would be lower than the actual predation intensity. This apparent drop in predation intensity results from the lack of representation of rare species in smaller samples that are never attacked. It is especially problematic because this difference is substantial for all evenness. This observation is consistent with the findings by Smith et al. (2021) where they demonstrated the effects of overdispersion and zero inflation using count data of predation traces. They concluded that the crucial element underlying these effects was sample size. Their results support our findings that predation measures lack reliability in a small sample size.

### Effect on inferred selectivity

Predation is known to impact the structure of a community, including the overall richness, distribution and evenness (Schemske et al. 2009; Freestone et al. 2011, 2020). It is therefore important to evaluate the inherent dependence of the predation inferences on the community structure before considering the evolutionary impact of predation on shaping the community structure in deep time. Our models demonstrate that the inferred number of prey species may depend on the evenness of the live community. Communities with low evenness differ significantly from the original prey species and yield fewer inferred prey species even when the predation is non-selective (Case 1). This may lead to the development of an artificial selectivity, primarily driven by the preferential counting of the dominant species and not by the biological preference demonstrated by the predators. Therefore, any community with low evenness suffers from the high likelihood of underrepresenting the number of prey species. The deviation from the true prey-species richness is higher for smaller sample size and lower predation intensity. Communities with higher predation intensity will provide the true prey-species richness at a smaller sample size than communities with lower predation intensity. Selective predation (as indicated by Case 2 and 3) also creates similar deviations.

The sensitivity of inferred prey species richness on sample size, evenness, and original predation selectivity makes the comparison of prey species richness in spatially or temporally distinct assemblages somewhat unreliable unless they are normalized. This is especially important when comparing predation estimates from assemblages representing different time-bins or environments likely to show varying diversity/evenness.

## Paleontological case study

The assemblages from the four Florida localities have been used to interpret the relationship between durophagy and drilling predation (Chattopadhyay and Baumiller, 2010). However, the study’s conclusions did not consider sample size or community structure. The assemblages at these localities differ in their evenness and sample size (Table 2). Only in Miami Canal, predatory attacks (durophagous and drilling) are guided by the relative abundance of prey species and hence deviates from non-selective predation. The sample size-standardized resampling protocol revealed a significant difference in pairwise comparison for all inferred predation intensity and prey species richness estimates. This implies that the differences in predation measure across assemblages are largely independent of sampling intensity or the evenness of the assemblage.

It is essential to recognize that several factors played a role in this particular case, making these assemblages less susceptible to community evenness and sampling intensity. Because three localities (Punta Gorda, Mc Queens pit, Chiquita) are showing non-selective predation, they are less likely to be affected by sample size. Moreover, they have medium to high evenness, making them less sensitive to sample size. Miami Canal, however, is characterized by low evenness (0.31), shows evidence of selective predation and low predation intensity (PI_T_<0.2). Assemblages with these characteristics are more prone to differ significantly from actual predation measures at small sample sizes (Fig 4A). Because Miami Canal has the largest sample size among the localities, makes it less likely to be affected by these factors. Hence, the observed S_prey.drill_ and S_prey.repair_ are least likely to be affected by these factors.

## Proposed protocol of post-facto standardization of predation data

The following protocol may be followed to compare predation intensity and selectivity of spatially/temporally distinct assemblages to avoid misinterpretation. Using the protocol in the simulation model, inferred predation intensity (PI_T.inf_) and inferred prey-species richness (S_prey.inf_) need to be calculated at a specific step size of 100 for all assemblages. The step size of all assemblages should be equal till the last step when the remaining number of individuals in that assemblage are drawn. The step size can be lowered till 30 if the total assemblage size is small. Reducing the step size any further may create erroneous results due to a smaller sample size (Kosloski et al. 2008; Dietl and Kosloski 2013; Smith et al. 2022). To understand the sensitivity of inferred predation intensity (PI_T_) on sample size, the assemblage with the smallest sample size should be considered as the reference. The distribution of inferred predation intensity (PI_T_) for all assemblages should be compared at that sample size by a pairwise comparison using Kolmogorov-Smirnov (K-S) test. Suppose the pair-wise comparison yields a significant difference between two assemblages. In that case the differences in inferred predation intensity (PI_T_) cannot be explained by sample size alone and hence, likely to represent the actual variation. These pairs would be considered comparable at that sample size. Suppose assemblages show non-significant difference in the pairwise K-S test. In that case, we cannot reject the possibility of small sample size influencing the inferred predation intensity and hence, should not be considered for further comparative analysis at that pre-selected sample size (Fig 11). The same process can be repeated to understand the sensitivity of the inferred prey-species richness (S_prey.inf_) on sample size.

**Figure 11.**
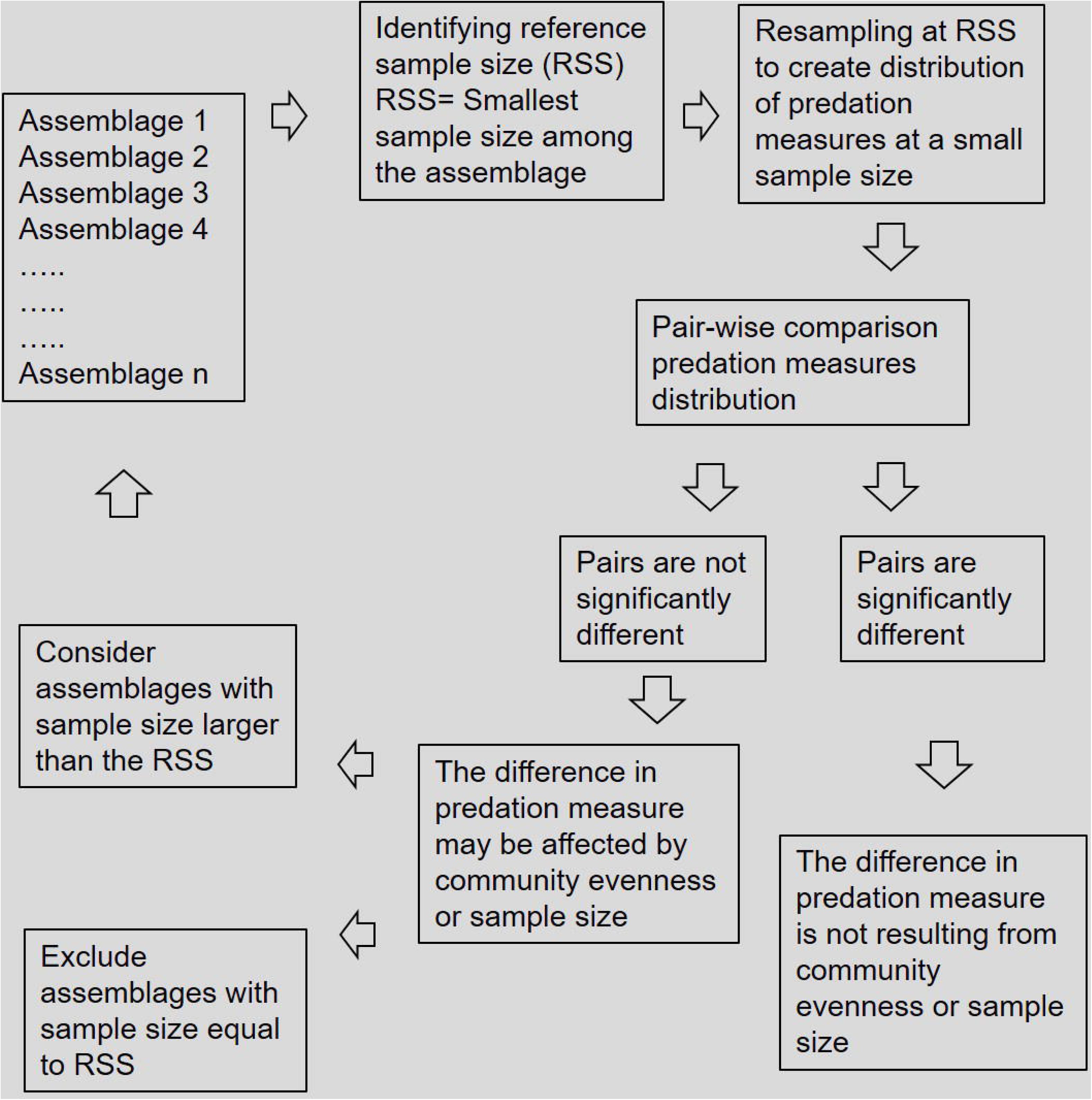
Flowchart of the general framework of proposed method of the post-hoc standardization. [Figure 11. Two columns; Grayscale]

The next step is to compare the remaining pairs with non-significant differences at different sample sizes. If any pair contains the assemblage with previously set reference sample size (smallest sample size), then we cannot include the pair for further analysis. For other pairs where the smallest sample size is larger than the previously set reference, the entire process described above should be repeated. The number of pairs that yield statistical significance can be considered comparable at the new reference sample size. This iteration should be performed with increasing sample sizes till the maximum number of assemblage pairs show significant differences in distribution of inferred predation intensity (PI_T_) and prey species richness (S_prey.inf_). Finally, the pairs that show non-significant differences even at the largest sample size, we cannot reject the influence of sampling intensity and inherent community structure in shaping the predation measures. They should be excluded from comparative analyses of predation signals. Estimating PI_prey_ is difficult, especially for cases where rare species are attacked; excluding species without any predation trace while calculating PI_T_ may give us some insight.

## Caveats and implications

The fossil record of predation has shaped our understanding of how the nature of biotic interaction changed over time and its role as an evolutionary mechanism. Preserved traces, such as drill holes and repair scars, are some of the best quantifiable predation proxies and are often used to assess the evolutionary impact of predation in deep time (Vermeij et al. 1981; Alexander and Dietl 2003; Kelley and Hansen 2003). However, studies aiming to evaluate the predation trend through time are often forced to use predation data from discrete assemblages that differ in sample size, inherent community evenness, and the type of predation selectivity (Harper 2016). Our study demonstrates the effect of such factors on the inferred predation intensity and the recognized prey richness. Comparison between temporally separated collections, such as Paleozoic and Cenozoic predation records that are known to be different in the sample size (and probably predatory behavior), are susceptible to such factors.

Our proposed method of post-facto standardization will be essential for such comparisons and to establish the true nature of biotic interaction through time. It is important to recognize that the proposed protocol is a preliminary attempt toward standardization without considering several complexities. The simulations are primarily developed for communities that are preserving the community structure of the live communities. A number of actualistic studies revealed, however, that the death/fossil assemblage can substantially differ from the live assemblage because they are typically time-averaged, representing a mix of multiple generations (Kidwell et al. 1991; Kidwell and Flessa 1995; Kidwell 2007; Olszewski and Kidwell 2007; Tomašových and Kidwell 2009; Kidwell and Tomasovych 2013; Bhattacherjee et al. 2021). If a specific section of the live community is preferentially lost due to preservation and if the predation signature of those specimens differs from the remaining assemblage, the proposed standardization method will fail to detect that. For example, some predation attempts are size-selective, and larger size classes often show higher predation resistance and lower predation intensity. Because the preservation potential of smaller size classes is lower than larger ones (Cooper et al. 2006), the elective absence of small size class in the fossils would result in a low inferred predation intensity compared to the original value. Multiple interactions during the lifetime or after the death of the prey may change the frequency of the overall assemblage (Kosloski 2011; Gordillo and Archuby 2014). A molluscan community affected by drilling predation may also be subjected to crushing predation; because the durophagous predators only go after the live prey (non-drilled), the relative proportion of drilled shells increase if the predators successfully destroy the shells as part of the predation process (Smith et al. 2019). Predation style and resulting predation trace also differ among predators. Two of the most common types of predations studied in the fossil, drilling and durophagy, are quite different in several aspects. It is possible to identify successful and unsuccessful predation by observing the completeness of the drill holes, successful attacks by durophagous predators often result in unrecognizable fragmentation (Kosloski 2011; Leighton et al. 2016; Dyer et al. 2018). Repair scars represent a failed durophagous attack. Comparing drill holes and repair scars, therefore, are not without limitations. Although our study attempts to recognize the possible source of analytical bias and to recognize them in the observed database, it clearly glosses over the full complexities of predation style, post-mortem alteration and time-averaging. Following the direction of reconstructing fossil assemblages from live data using the modeling approach (Olszewski 2004, 2012), we plan to develop more inclusive frameworks in future to address such complexities.

## Conclusions

The effect of community structure and sampling intensity on the inferred predation estimates is rarely explored. Using a resampling technique, our study demonstrates the impact of these aspects on the estimates of predation intensity and the number of prey species. Our results show that the communities with highly selective predation are the most sensitive to sampling intensity, and the inferred predation intensity of these assemblages can substantially deviate from the actual value. In contrast, predation intensity for non-selective predation tends to be unaffected by sampling intensity. Inferred prey-species richness is also influenced by the nature of community evenness, predation selectivity, and actual predation intensity. For non-selective predation, communities with low evenness and low predation intensity are highly sensitive to sample size. The inferred prey-species richness can be underrepresented significantly at smaller sample size. For selective predation, the sensitivity depends on the nature of selection. The inferred prey-species richness deviates significantly when rare species are attacked preferentially. Our study also provides a framework of post-facto standardization of the predation data to remove the effect of sample size/evenness during comparison. The proposed method, although simple, will provide a fundamental framework for comparison of discrete assemblages as they are often characterized by a difference in sample size, community structure and predation selectivity.

## Supporting information

Supplementary R script (S3)

## Acknowledgments

This project was supported by the SERB core research grant (CRG/2018/002604). The IISER Pune Doctoral fellowship supported MB. We would like to thank Satyaki Mazumdar for his help in designing the R code. We are grateful to the associate editor, Erin Saupe and reviewer, Jansen Smith for their detailed comments that substantially improved the quality of the manuscript.

## Supplementary materials

Figure S1. The plot showing variation in inferred predation intensity (PI_T.inf_) and inferred the number of prey species (S_prey.inf_) with specific sample sizes for different model assemblages. The rows represent different degrees of selectivity of predation and the columns indicate predation intensity in the original assemblage (PI_T_). The warmer colors represent higher evenness. [Figure S1. Two columns; Color]

S2. This is a compiled datasheet used for analyzing the predation estimates across four Plio-Pleistocene fossil assemblages of Florida (Chattopadhyay and Baumiller, 2010)

S3. This is the R-script used for the present study. The required data is available at S2.

