## Supplementary figures and images for "Community evenness and sample size affect estimates of predation intensity and prey selection: A model-based validation"

### Supplementary Figure (S1)

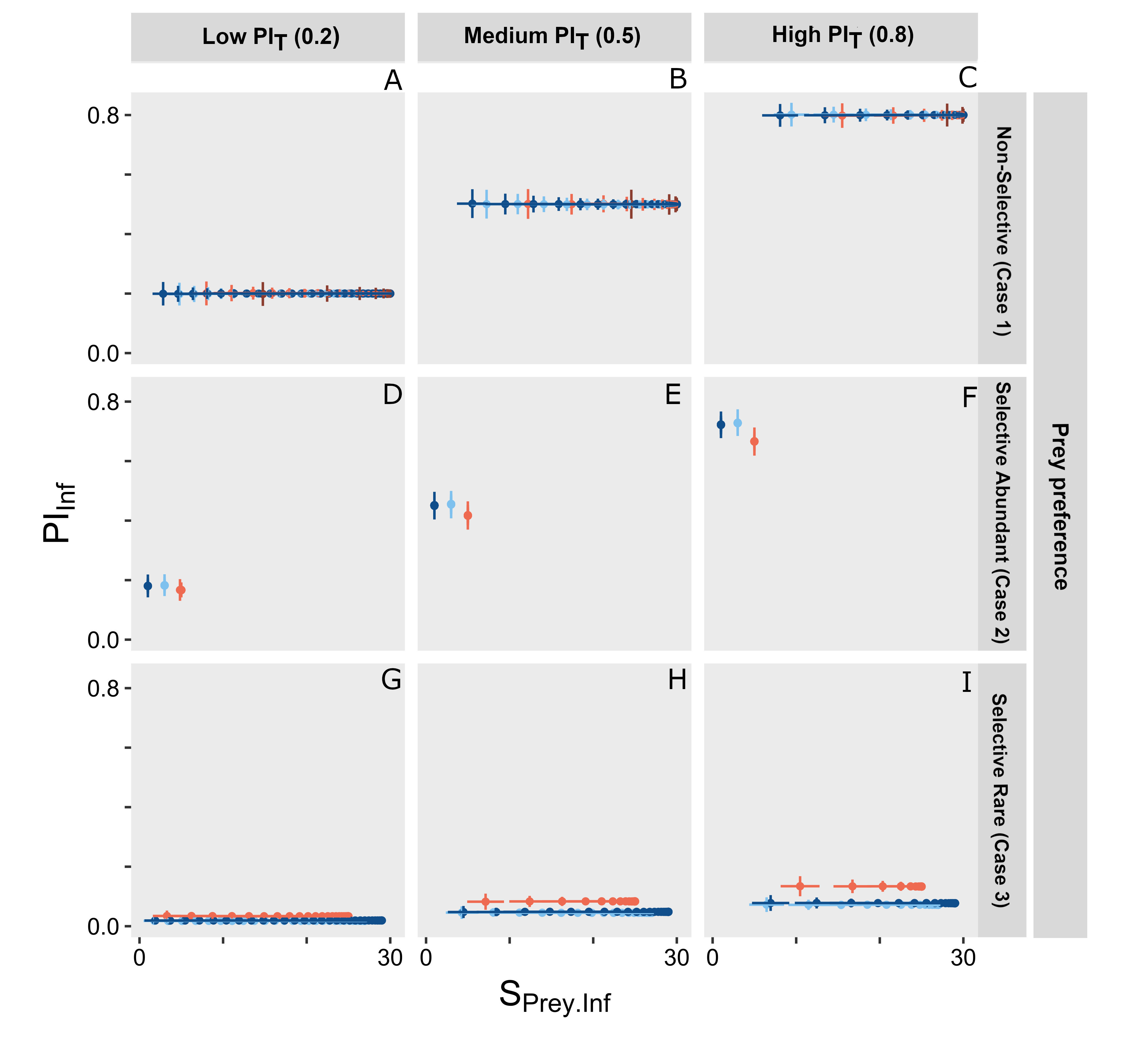
