## Supplementary R script (S3) for "Community evenness and sample size affect estimates of predation intensity and prey selection: A model-based validation"

```
## --------------------------
##
## Project: Community evenness and sample size affect estimates of
predation intensity and prey selection: A model-based validation
##
## Purpose of script: Reshape data to be used for analyses
##
## Author: Madhura Bhattacherjee, Devapriya Chattopadhyay
##
##
##
##
## Last Modified: 2022-08-23
##
## --------------------------
##
##
##
##
## --------------------------

require(vegan)
require(MASS)
require(ggplot2)

####Setting up working directory#############
setwd("F:/Madhura/Drilling frequency/Drilling frequency")


#####creating max evenness dataset for 3000 individuals (ET=1)#####
Species=array(0,dim = c(3000,1))
subsamps1=list()
for(i in 1:30)
{
  Species=rep(i,100)
  out=data.frame(Species)
  subsamps1[[i]] <-out
}
identity_species <- do.call("rbind",subsamps1)

d=c(rep(1,20),rep(0,80))
LowD=rep(d,30)
e=c(rep(1,50),rep(0,50))
MedD=rep(e,30)
f=c(rep(1,80),rep(0,20))
HighD=rep(f,30)
maxeven=data.frame(identity_species,LowD,MedD,HighD)
Individual=rep(1,3000)
vareven=data.frame(Individual,identity_species,LowD,MedD,HighD)

####Calculating evenness of the data frame#####
wideform=as.data.frame.matrix(table(vareven$Individual,vareven$Species))
evenness=(diversity(wideform)/log(specnumber(wideform)))

#########Simulation considering Low PIT (0.2)#################
DFtype="Low"
Eventype=evenness
```

```
Selection="NS"
Case=1
PI_T=0.2
prey_actual=30
a=cbind(DFtype,Eventype,Selection,Case)

randomnumber=vector(mode = "integer",length=3000)
raredata=array(0,dim = c(3000,2))
num_species1=vector(mode="integer",length=30)
num_drilled=vector(mode = "integer", length = 30)
DF=vector(mode = "integer", length = 30)
Dev=vector(mode = "integer", length = 30)
dev_numdrilled=vector(mode = "integer", length = 30)
max_evensubset=subset(maxeven,select = c(Species,LowD))
subsamps <- list()
subsamps1=list()
for (q in 1:1000)
{
  randomnumber=sample.int(3000,3000, replace = FALSE)
  raredata=max_evensubset[randomnumber[1:3000],]
  for(i in 1:30)
  {
    j=i*100
    uniquesp=unique(raredata$Species[1:j])
    num_species1[j]=length(uniquesp)
    uniquedrilled=unique(raredata$Species[which(raredata$LowD[1:j]==1)])
    DF[j]=(sum(raredata$LowD[1:j]))/j
    num_drilled[j]=length(uniquedrilled)
    Dev[j]=PI_T-DF[j]
    dev_numdrilled[j]=prey_actual-num_drilled[j]
    out=data.frame(j,num_species1[j],num_drilled[j],DF[j],Dev[j],dev_numdrilled[j])
    subsamps1[[i]] <-out
  }
  subsamples1 <- do.call("rbind",subsamps1)
  out2=data.frame(q,subsamples1)
  subsamps[[q]]=out2
  subsamples <- do.call("rbind",subsamps)
}
subsamples=as.data.frame(subsamples)



#####################################################################################
############For calculating average and variance for low PIT
(0.2)#######################################################
uniquedrilledavg=vector(mode = "integer", length = 30)
uniquedrilledvar=vector(mode = "integer", length = 30)
DFavg=vector(mode = "integer", length = 30)
DFvar=vector(mode = "integer", length = 30)
Devavg=vector(mode = "integer", length = 30)
Devvar=vector(mode = "integer", length = 30)
devpreyavg=vector(mode = "integer", length = 30)
devpreyvar=vector(mode = "integer", length = 30)

for(i in 1:30)
{
  j=i*100
```

```
  uniquedrilledavg[j]=mean(subsamples$num_drilled.j.
[which(subsamples$j==j)])
  uniquedrilledvar[j]=sd(subsamples$num_drilled.j.[which(subsamples$j==j)])
  DFavg[j]=mean(subsamples$DF.j.[which(subsamples$j==j)])
  DFvar[j]=sd(subsamples$DF.j.[which(subsamples$j==j)])
  Devavg[j]=mean(subsamples$Dev.j.[which(subsamples$j==j)])
  Devvar[j]=sd(subsamples$Dev.j.[which(subsamples$j==j)])
  devpreyavg[j]=mean(subsamples$dev_numdrilled.j.[which(subsamples$j==j)])
  devpreyvar[j]=sd(subsamples$dev_numdrilled.j.[which(subsamples$j==j)])
  out=data.frame(j,uniquedrilledavg[j],uniquedrilledvar[j],DFavg[j],DFvar[j],Devavg[j],
  subsamps1[[i]] <-out
}
case1_1even_low <- do.call("rbind",subsamps1)
Drilledsp_avg=case1_1even_low $uniquedrilledavg.j.
Drilledsp_var=case1_1even_low $uniquedrilledvar.j.
DF_avg=case1_1even_low $DFavg.j.
DF_var=case1_1even_low$DFvar.j.
DE_avg=case1_1even_low$Devavg.j.
DE_var=case1_1even_low$Devvar.j.
devpreyavg=case1_1even_low$devpreyavg.j.
devpreyavg=case1_1even_low$devpreyvar.j.


######################simulation considering medium PIT
(0.5)##################
DFtype="Medium"
Eventype=evenness
Selection="NS"
Case=1
PI_T=0.5
prey_actual=30
b=cbind(DFtype,Eventype,Selection,Case)

randomnumber=vector(mode = "integer",length=3000)
raredata=array(0,dim = c(3000,2))
num_species1=vector(mode="integer",length=30)
num_drilled=vector(mode = "integer", length = 30)
DF=vector(mode = "integer", length = 30)
Dev=vector(mode = "integer", length = 30)
dev_numdrilled=vector(mode = "integer", length = 30)
max_evensubset=subset(maxeven,select = c(Species,MedD))
subsamps <- list()
subsamps1=list()
for (q in 1:1000)
{
  randomnumber=sample.int(3000,3000, replace = FALSE)
  raredata=max_evensubset[randomnumber[1:3000],]
  for(i in 1:30)
  {
    j=i*100
    uniquesp=unique(raredata$Species[1:j])
    num_species1[j]=length(uniquesp)
    uniquedrilled=unique(raredata$Species[which(raredata$MedD[1:j]==1)])
    num_drilled[j]=length(uniquedrilled)
    DF[j]=(sum(raredata$MedD[1:j]))/j
    Dev[j]=PI_T-DF[j]
    dev_numdrilled[j]=prey_actual-num_drilled[j]
    out=data.frame(j,num_species1[j],num_drilled[j],DF[j],Dev[j],dev_numdrilled[j])
```

```
    subsamps1[[i]] <-out
  }
  subsamples1 <- do.call("rbind",subsamps1)
  out2=data.frame(q,subsamples1)
  subsamps[[q]]=out2
  subsamples <- do.call("rbind",subsamps)
}
subsamples=as.data.frame(subsamples)



#####################################################################################
############For calculating average and variance medium D medium PIT
(0.5)#########################################################
uniquedrilledavg=vector(mode = "integer", length = 30)
uniquedrilledvar=vector(mode = "integer", length = 30)
DFavg=vector(mode = "integer", length = 30)
DFvar=vector(mode = "integer", length = 30)
Devavg=vector(mode = "integer", length = 30)
Devvar=vector(mode = "integer", length = 30)
devpreyavg=vector(mode = "integer", length = 30)
devpreyvar=vector(mode = "integer", length = 30)

for(i in 1:30)
{
  j=i*100
  uniquedrilledavg[j]=mean(subsamples$num_drilled.j.
[which(subsamples$j==j)])
  uniquedrilledvar[j]=sd(subsamples$num_drilled.j.[which(subsamples$j==j)])
  DFavg[j]=mean(subsamples$DF.j.[which(subsamples$j==j)])
  DFvar[j]=sd(subsamples$DF.j.[which(subsamples$j==j)])
  Devavg[j]=mean(subsamples$Dev.j.[which(subsamples$j==j)])
  Devvar[j]=sd(subsamples$Dev.j.[which(subsamples$j==j)])
  devpreyavg[j]=mean(subsamples$dev_numdrilled.j.[which(subsamples$j==j)])
  devpreyvar[j]=sd(subsamples$dev_numdrilled.j.[which(subsamples$j==j)])
  out=data.frame(j,uniquedrilledavg[j],uniquedrilledvar[j],DFavg[j],DFvar[j],Devavg[j],
  subsamps1[[i]] <-out
}
case1_1even_med <- do.call("rbind",subsamps1)
Drilledsp_avg=case1_1even_med$uniquedrilledavg.j.
Drilledsp_var=case1_1even_med$uniquedrilledvar.j.
DF_avg=case1_1even_med$DFavg.j.
DF_var=case1_1even_med$DFvar.j.
DE_avg=case1_1even_med$Devavg.j.
DE_var=case1_1even_med$Devvar.j.
devpreyavg=case1_1even_med$devpreyavg.j.
devpreyvar=case1_1even_med$devpreyvar.j.

##############Simulation considering high PIT (0.8)#################
DFtype="High"
Eventype=evenness
Selection="NS"
Case=1
PI_T=0.8
prey_actual=30
c=cbind(DFtype,Eventype,Selection,Case)

randomnumber=vector(mode = "integer",length=3000)
```

```
raredata=array(0,dim = c(3000,2))
num_species1=vector(mode="integer",length=30)
num_drilled=vector(mode = "integer", length = 30)
DF=vector(mode = "integer", length = 30)
max_evensubset=subset(maxeven,select = c(Species,HighD))
Dev=vector(mode = "integer", length = 30)
dev_numdrilled=vector(mode = "integer", length = 30)


subsamps <- list()
subsamps1=list()
for (q in 1:1000)
{
  randomnumber=sample.int(3000,3000, replace = FALSE)
  raredata=max_evensubset[randomnumber[1:3000],]
  for(i in 1:30)
  {
    j=i*100
    uniquesp=unique(raredata$Species[1:j])
    num_species1[j]=length(uniquesp)
    uniquedrilled=unique(raredata$Species[which(raredata$HighD[1:j]==1)])
    num_drilled[j]=length(uniquedrilled)
    DF[j]=(sum(raredata$HighD[1:j]))/j
    Dev[j]=PI_T-DF[j]
    dev_numdrilled[j]=prey_actual-num_drilled[j]
    out=data.frame(j,num_species1[j],num_drilled[j],DF[j],Dev[j],dev_numdrilled[j])
    subsamps1[[i]] <-out
  }
  subsamples1 <- do.call("rbind",subsamps1)
  out2=data.frame(q,subsamples1)
  subsamps[[q]]=out2
  subsamples <- do.call("rbind",subsamps)
}
subsamples=as.data.frame(subsamples)



####################################################################################
############For calculating average and variance high D for high PIT
(0.8)#########################################################
uniquedrilledavg=vector(mode = "integer", length = 30)
uniquedrilledvar=vector(mode = "integer", length = 30)
DFavg=vector(mode = "integer", length = 30)
DFvar=vector(mode = "integer", length = 30)
Devavg=vector(mode = "integer", length = 30)
Devvar=vector(mode = "integer", length = 30)
devpreyavg=vector(mode = "integer", length = 30)
devpreyvar=vector(mode = "integer", length = 30)

for(i in 1:30)
{
  j=i*100
  uniquedrilledavg[j]=mean(subsamples$num_drilled.j.
[which(subsamples$j==j)])
  uniquedrilledvar[j]=sd(subsamples$num_drilled.j.[which(subsamples$j==j)])
  DFavg[j]=mean(subsamples$DF.j.[which(subsamples$j==j)])
  DFvar[j]=sd(subsamples$DF.j.[which(subsamples$j==j)])
  Devavg[j]=mean(subsamples$Dev.j.[which(subsamples$j==j)])
```

```
  Devvar[j]=sd(subsamples$Dev.j.[which(subsamples$j==j)])
  devpreyavg[j]=mean(subsamples$dev_numdrilled.j.[which(subsamples$j==j)])
  devpreyvar[j]=sd(subsamples$dev_numdrilled.j.[which(subsamples$j==j)])
  out=data.frame(j,uniquedrilledavg[j],uniquedrilledvar[j],DFavg[j],DFvar[j],Devavg[j],
  subsamps1[[i]] <-out
}

case1_1even_high <- do.call("rbind",subsamps1)
Drilledsp_avg=case1_1even_high$uniquedrilledavg.j.
Drilledsp_var=case1_1even_high$uniquedrilledvar.j.
DF_avg=case1_1even_high$DFavg.j.
DF_var=case1_1even_high$DFvar.j.
DE_avg=case1_1even_high$Devavg.j.
DE_var=case1_1even_high$Devvar.j.
devpreyavg=case1_1even_high$devpreyavg.j.
devpreyavg=case1_1even_high$devpreyvar.j.


##########Combining the results of all simulations in
ET=1############################################
combined_max_even_2=rbind(case1_1even_low,case1_1even_med,case1_1even_high)

##############################################################################################
######################Creating dataset for Evenness
(ET)=0.7########################################
Species=array(0,dim = c(3000,1))
subsamps1=list()
subsamps2=list()

for(i in 1:5)
{
  Species=rep(i,500)
  out=data.frame(Species)
  subsamps1[[i]] <-out
}
identity_species1 <- do.call("rbind",subsamps1)
for(i in 6:30)
{
  Species=rep(i,20)
  out=data.frame(Species)
  subsamps2[[i]] <-out
}
identity_species2 <- do.call("rbind",subsamps2)
identity_species<-rbind(identity_species1,identity_species2)
#######low predation intensity PIT (0.2) for 3000 individuals#########
d1=c(rep(1,100),rep(0,400))
LowD1=rep(d1,5)
d2=c(rep(1,4),rep(0,16))
LowD2=rep(d2,25)
LowD=c(LowD1,LowD2)
##########medium predation intensity PIT(0.5) for 3000 individuals#####
md1=c(rep(1,250),rep(0,250))
MedD1=rep(md1,5)
md2=c(rep(1,10),rep(0,10))
MedD2=rep(md2,25)
MedD=c(MedD1,MedD2)
##########high predation intensity PIT(0.8) for 3000 individuals#####
md1=c(rep(1,400),rep(0,100))
```

```
HighD1=rep(md1,5)
md2=c(rep(1,16),rep(0,4))
HighD2=rep(md2,25)
HighD=c(HighD1,HighD2)


Individual=rep(1,3000)
vareven=data.frame(Individual,identity_species,LowD,MedD,HighD)
wideform=as.data.frame.matrix(table(vareven$Individual,vareven$Species))
evenness=(diversity(wideform)/log(specnumber(wideform)))




######################################################################

#####(Case 1)Simulation for Et=0.7 at Low PIT=0.2##############
DFtype="Low"
Eventype=evenness
Selection="NS"
Case=1
PI_T=0.2
prey_actual=30
p=cbind(DFtype,Eventype,Selection,Case)


randomnumber=vector(mode = "integer",length=3000)
raredata=array(0,dim = c(3000,2))
num_species1=vector(mode="integer",length=30)
num_drilled=vector(mode = "integer", length = 30)
DF=vector(mode = "integer", length = 30)
varevensubset=subset(vareven,select = c(Species,LowD))
Dev=vector(mode = "integer", length = 30)
dev_numdrilled=vector(mode = "integer", length = 30)

subsamps <- list()
subsamps1=list()
for (q in 1:1000)
{
  randomnumber=sample.int(3000,3000, replace = FALSE)
  raredata=varevensubset[randomnumber[1:3000],] #####shuffling the
data################
  for(i in 1:30)
  {
    j=i*100
    uniquesp=unique(raredata$Species[1:j])
    num_species1[j]=length(uniquesp)
    uniquedrilled=unique(raredata$Species[which(raredata$LowD[1:j]==1)])
    num_drilled[j]=length(uniquedrilled)
    DF[j]=(sum(raredata$LowD[1:j]))/j
    Dev[j]=PI_T-DF[j]
    dev_numdrilled[j]=prey_actual-num_drilled[j]
    out=data.frame(j,num_species1[j],num_drilled[j],DF[j],Dev[j],dev_numdrilled[j])
    subsamps1[[i]] <-out
  }
  subsamples1 <- do.call("rbind",subsamps1)
  out2=data.frame(q,subsamples1)
  subsamps[[q]]=out2
```

```
  subsamples <- do.call("rbind",subsamps)
}
subsamples=as.data.frame(subsamples)



############For calculating average and variance at low
PIT##########################################################
uniquedrilledavg=vector(mode = "integer", length = 30)
uniquedrilledvar=vector(mode = "integer", length = 30)
DFavg=vector(mode = "integer", length = 30)
DFvar=vector(mode = "integer", length = 30)
Devavg=vector(mode = "integer", length = 30)
Devvar=vector(mode = "integer", length = 30)
devpreyavg=vector(mode = "integer", length = 30)
devpreyvar=vector(mode = "integer", length = 30)

for(i in 1:30)
{
  j=i*100
  uniquedrilledavg[j]=mean(subsamples$num_drilled.j.
[which(subsamples$j==j)])
  uniquedrilledvar[j]=sd(subsamples$num_drilled.j.[which(subsamples$j==j)])
  DFavg[j]=mean(subsamples$DF.j.[which(subsamples$j==j)])
  DFvar[j]=sd(subsamples$DF.j.[which(subsamples$j==j)])
  Devavg[j]=mean(subsamples$Dev.j.[which(subsamples$j==j)])
  Devvar[j]=sd(subsamples$Dev.j.[which(subsamples$j==j)])
  devpreyavg[j]=mean(subsamples$dev_numdrilled.j.[which(subsamples$j==j)])
  devpreyvar[j]=sd(subsamples$dev_numdrilled.j.[which(subsamples$j==j)])
  out=data.frame(j,uniquedrilledavg[j],uniquedrilledvar[j],DFavg[j],DFvar[j],Devavg[j],[illegible]
  subsamps1[[i]] <-out
}
case1_0.7even_low <- do.call("rbind",subsamps1)
Drilledsp_avg=case1_0.7even_low$uniquedrilledavg.j.
Drilledsp_var=case1_0.7even_low$uniquedrilledvar.j.
DF_avg=case1_0.7even_low$DFavg.j.
DF_var=case1_0.7even_low$DFvar.j.
DE_avg=case1_0.7even_low$Devavg.j.
DE_var=case1_0.7even_low$Devvar.j.
devpreyavg=case1_0.7even_low$devpreyavg.j.
devpreyvar=case1_0.7even_low$devpreyvar.j.

###########################(Case 1) Simulation for Et=0.7 at MEDIUM
PIT=0.5############################################
DFtype="Medium"
Eventype=evenness
Selection="NS"
Case=1
PI_T=0.5
prey_actual=30
qz=cbind(DFtype,Eventype,Selection,Case)

randomnumber=vector(mode = "integer",length=3000)
raredata=array(0,dim = c(3000,2))
num_species1=vector(mode="integer",length=30)
num_drilled=vector(mode = "integer", length = 30)
DF=vector(mode = "integer", length = 30)
varevensubset=subset(vareven,select = c(Species,MedD))
```

```
Dev=vector(mode = "integer", length = 30)
dev_numdrilled=vector(mode = "integer", length = 30)

subsamps <- list()
subsamps1=list()
for (q in 1:1000)
{
  randomnumber=sample.int(3000,3000, replace = FALSE)
  raredata=varevensubset[randomnumber[1:3000],]
  for(i in 1:30)
  {
    j=i*100
    uniquesp=unique(raredata$Species[1:j])
    num_species1[j]=length(uniquesp)
    uniquedrilled=unique(raredata$Species[which(raredata$MedD[1:j]==1)])
    num_drilled[j]=length(uniquedrilled)
    DF[j]=(sum(raredata$MedD[1:j]))/j
    Dev[j]=PI_T-DF[j]
    dev_numdrilled[j]=prey_actual-num_drilled[j]
    out=data.frame(j,num_species1[j],num_drilled[j],DF[j],Dev[j],dev_numdrilled[j])
    subsamps1[[i]] <-out
  }
  subsamples1 <- do.call("rbind",subsamps1)
  out2=data.frame(q,subsamples1)
  subsamps[[q]]=out2
  subsamples <- do.call("rbind",subsamps)
}
subsamples=as.data.frame(subsamples)



############For calculating average and variance at medium
PIT############################################################
uniquedrilledavg=vector(mode = "integer", length = 30)
uniquedrilledvar=vector(mode = "integer", length = 30)
DFavg=vector(mode = "integer", length = 30)
DFvar=vector(mode = "integer", length = 30)
Devavg=vector(mode = "integer", length = 30)
Devvar=vector(mode = "integer", length = 30)
devpreyavg=vector(mode = "integer", length = 30)
devpreyvar=vector(mode = "integer", length = 30)

for(i in 1:30)
{
  j=i*100
  uniquedrilledavg[j]=mean(subsamples$num_drilled.j.
[which(subsamples$j==j)])
  uniquedrilledvar[j]=sd(subsamples$num_drilled.j.[which(subsamples$j==j)])
  DFavg[j]=mean(subsamples$DF.j.[which(subsamples$j==j)])
  DFvar[j]=sd(subsamples$DF.j.[which(subsamples$j==j)])
  Devavg[j]=mean(subsamples$Dev.j.[which(subsamples$j==j)])
  Devvar[j]=sd(subsamples$Dev.j.[which(subsamples$j==j)])
  devpreyavg[j]=mean(subsamples$dev_numdrilled.j.[which(subsamples$j==j)])
  devpreyvar[j]=sd(subsamples$dev_numdrilled.j.[which(subsamples$j==j)])
  out=data.frame(j,uniquedrilledavg[j],uniquedrilledvar[j],DFavg[j],DFvar[j],Devavg[j],[illegible]
  subsamps1[[i]] <-out
}
case1_0.7even_med<- do.call("rbind",subsamps1)
```

```
Drilledsp_avg=case1_0.7even_med$uniquedrilledavg.j.
Drilledsp_var=case1_0.7even_med$uniquedrilledvar.j.
DF_avg=case1_0.7even_med$DFavg.j.
DF_var=case1_0.7even_med$DFvar.j.
DE_avg=case1_0.7even_med$Devavg.j.
DE_var=case1_0.7even_med$Devvar.j.
devpreyavg=case1_0.7even_med$devpreyavg.j.
devpreyvar=case1_0.7even_med$devpreyvar.j.



#############################(Case 1) Simulation for Et=0.7 at HIGH
PIT=0.8################################################
DFtype="High"
Eventype=evenness
Selection="NS"
Case=1
PI_T=0.8
prey_actual=30
r=cbind(DFtype,Eventype,Selection,Case)

randomnumber=vector(mode = "integer",length=3000)
raredata=array(0,dim = c(3000,2))
num_species1=vector(mode="integer",length=30)
num_drilled=vector(mode = "integer", length = 30)
DF=vector(mode = "integer", length = 30)
Dev=vector(mode = "integer", length = 30)
varevensubset=subset(vareven,select = c(Species,HighD))
dev_numdrilled=vector(mode = "integer", length = 30)

subsamps <- list()
subsamps1=list()
for (q in 1:1000)
{
  randomnumber=sample.int(3000,3000, replace = FALSE)
  raredata=varevensubset[randomnumber[1:3000],]
  for(i in 1:30)
  {
    j=i*100
    uniquesp=unique(raredata$Species[1:j])
    num_species1[j]=length(uniquesp)
    uniquedrilled=unique(raredata$Species[which(raredata$HighD[1:j]==1)])
    num_drilled[j]=length(uniquedrilled)
    DF[j]=(sum(raredata$HighD[1:j]))/j
    Dev[j]=PI_T-DF[j]
    dev_numdrilled[j]=prey_actual-num_drilled[j]
    out=data.frame(j,num_species1[j],num_drilled[j],DF[j],Dev[j],dev_numdrilled[j])
    subsamps1[[i]] <-out
  }
  subsamples1 <- do.call("rbind",subsamps1)
  out2=data.frame(q,subsamples1)
  subsamps[[q]]=out2
  subsamples <- do.call("rbind",subsamps)
}
subsamples=as.data.frame(subsamples)
```

```
##########################################################################################
############For calculating average and variance at high
PIT############################################################
uniquedrilledavg=vector(mode = "integer", length = 30)
uniquedrilledvar=vector(mode = "integer", length = 30)
DFavg=vector(mode = "integer", length = 30)
DFvar=vector(mode = "integer", length = 30)
Devavg=vector(mode = "integer", length = 30)
Devvar=vector(mode = "integer", length = 30)
devpreyavg=vector(mode = "integer", length = 30)
devpreyvar=vector(mode = "integer", length = 30)

for(i in 1:30)
{
  j=i*100
  uniquedrilledavg[j]=mean(subsamples$num_drilled.j.
[which(subsamples$j==j)])
  uniquedrilledvar[j]=sd(subsamples$num_drilled.j.[which(subsamples$j==j)])
  DFavg[j]=mean(subsamples$DF.j.[which(subsamples$j==j)])
  DFvar[j]=sd(subsamples$DF.j.[which(subsamples$j==j)])
  Devavg[j]=mean(subsamples$Dev.j.[which(subsamples$j==j)])
  Devvar[j]=sd(subsamples$Dev.j.[which(subsamples$j==j)])
  devpreyavg[j]=mean(subsamples$dev_numdrilled.j.[which(subsamples$j==j)])
  devpreyvar[j]=sd(subsamples$dev_numdrilled.j.[which(subsamples$j==j)])
  out=data.frame(j,uniquedrilledavg[j],uniquedrilledvar[j],DFavg[j],DFvar[j],Devavg[j],[illegible]
  subsamps1[[i]] <-out
}
case1_0.7even_high <- do.call("rbind",subsamps1)
Drilledsp_avg=case1_0.7even_high$uniquedrilledavg.j.
Drilledsp_var=case1_0.7even_high$uniquedrilledvar.j.
DF_avg=case1_0.7even_high$DFavg.j.
DF_var=case1_0.7even_high$DFvar.j.
DE_avg=case1_0.7even_high$Devavg.j.
DE_var=case1_0.7even_high$Devvar.j.
devpreyavg=case1_0.7even_high$devpreyavg.j.
devpreyvar=case1_0.7even_high$devpreyvar.j.



#############Case 2:Selective predation for common species(first five
species(500 individuals each) attacked)#################

vareven[2501:3000,c(3,4,5)]=0 #############Creating Case 2 scenario  in
ET=0.7 dataset--------------------------

############### (Case 2)Simulation for Et=0.7 at Low PIT=0.2---------------
DFtype="Low"
Eventype=evenness
Selection="S"
Case=2
PI_T=0.17
prey_actual=5
s=cbind(DFtype,Eventype,Selection,Case)


randomnumber=vector(mode = "integer",length=3000)
raredata=array(0,dim = c(3000,2))
num_species1=vector(mode="integer",length=30)
```

```
num_drilled=vector(mode = "integer", length = 30)
DF=vector(mode = "integer", length = 30)
Dev=vector(mode = "integer", length = 30)
dev_numdrilled=vector(mode = "integer", length = 30)

varevensubset=subset(vareven,select = c(Species,LowD))
subsamps <- list()
subsamps1=list()
for (q in 1:1000)
{
  randomnumber=sample.int(3000,3000, replace = FALSE)
  raredata=varevensubset[randomnumber[1:3000],]
  for(i in 1:30)
  {
    j=i*100
    uniquesp=unique(raredata$Species[1:j])
    num_species1[j]=length(uniquesp)
    uniquedrilled=unique(raredata$Species[which(raredata$LowD[1:j]==1)])
    num_drilled[j]=length(uniquedrilled)
    DF[j]=(sum(raredata$LowD[1:j]))/j
    Dev[j]=PI_T-DF[j]
    dev_numdrilled[j]=prey_actual-num_drilled[j]
    out=data.frame(j,num_species1[j],num_drilled[j],DF[j],Dev[j],dev_numdrilled[j])
    subsamps1[[i]] <-out
  }
  subsamples1 <- do.call("rbind",subsamps1)
  out2=data.frame(q,subsamples1)
  subsamps[[q]]=out2
  subsamples <- do.call("rbind",subsamps)
}
subsamples=as.data.frame(subsamples)




############For calculating average and variance for low
PIT#######################################################
uniquedrilledavg=vector(mode = "integer", length = 30)
uniquedrilledvar=vector(mode = "integer", length = 30)
DFavg=vector(mode = "integer", length = 30)
DFvar=vector(mode = "integer", length = 30)
Devavg=vector(mode = "integer", length = 30)
Devvar=vector(mode = "integer", length = 30)
devpreyavg=vector(mode = "integer", length = 30)
devpreyvar=vector(mode = "integer", length = 30)

for(i in 1:30)
{
  j=i*100
  uniquedrilledavg[j]=mean(subsamples$num_drilled.j.
[which(subsamples$j==j)])
  uniquedrilledvar[j]=sd(subsamples$num_drilled.j.[which(subsamples$j==j)])
  DFavg[j]=mean(subsamples$DF.j.[which(subsamples$j==j)])
  DFvar[j]=sd(subsamples$DF.j.[which(subsamples$j==j)])
  Devavg[j]=mean(subsamples$Dev.j.[which(subsamples$j==j)])
  Devvar[j]=sd(subsamples$Dev.j.[which(subsamples$j==j)])
  devpreyavg[j]=mean(subsamples$dev_numdrilled.j.[which(subsamples$j==j)])
  devpreyvar[j]=sd(subsamples$dev_numdrilled.j.[which(subsamples$j==j)])
```

```
  out=data.frame(j,uniquedrilledavg[j],uniquedrilledvar[j],DFavg[j],DFvar[j],Devavg[j],[illegible]
  subsamps1[[i]] <-out
}
case2_0.7even_low <- do.call("rbind",subsamps1)
Drilledsp_avg=case2_0.7even_low$uniquedrilledavg.j.
Drilledsp_var=case2_0.7even_low$uniquedrilledvar.j.
DF_avg=case2_0.7even_low$DFavg.j.
DF_var=case2_0.7even_low$DFvar.j.
DE_avg=case2_0.7even_low$Devavg.j.
DE_var=case2_0.7even_low$Devvar.j.
devpreyavg=case1_0.7even_low$devpreyavg.j.
devpreyvar=case1_0.7even_low$devpreyvar.j.




############(Case 2)Simulation for Et=0.7 at medium PIT=0.5---------------
DFtype="Medium"
Eventype=evenness
Selection="S"
Case=2
PI_T=0.42
prey_actual=5
t=cbind(DFtype,Eventype,Selection,Case)

randomnumber=vector(mode = "integer",length=3000)
raredata=array(0,dim = c(3000,2))
num_species1=vector(mode="integer",length=30)
num_drilled=vector(mode = "integer", length = 30)
DF=vector(mode = "integer", length = 30)
Dev=vector(mode = "integer", length = 30)
dev_numdrilled=vector(mode = "integer", length = 30)

varevensubset=subset(vareven,select = c(Species,MedD))
subsamps <- list()
subsamps1=list()
for (q in 1:1000)
{
  randomnumber=sample.int(3000,3000, replace = FALSE)
  raredata=varevensubset[randomnumber[1:3000],]
  for(i in 1:30)
  {
    j=i*100
    uniquesp=unique(raredata$Species[1:j])
    num_species1[j]=length(uniquesp)
    uniquedrilled=unique(raredata$Species[which(raredata$MedD[1:j]==1)])
    num_drilled[j]=length(uniquedrilled)
    DF[j]=(sum(raredata$MedD[1:j]))/j
    Dev[j]=PI_T-DF[j]
    dev_numdrilled[j]=prey_actual-num_drilled[j]
    out=data.frame(j,num_species1[j],num_drilled[j],DF[j],Dev[j],dev_numdrilled[j])
    subsamps1[[i]] <-out
  }
  subsamples1 <- do.call("rbind",subsamps1)
  out2=data.frame(q,subsamples1)
  subsamps[[q]]=out2
  subsamples <- do.call("rbind",subsamps)
}
```

```
subsamples=as.data.frame(subsamples)


############For calculating average and variance at medium
PIT############################################################
uniquedrilledavg=vector(mode = "integer", length = 30)
uniquedrilledvar=vector(mode = "integer", length = 30)
DFavg=vector(mode = "integer", length = 30)
DFvar=vector(mode = "integer", length = 30)
Devavg=vector(mode = "integer", length = 30)
Devvar=vector(mode = "integer", length = 30)
devpreyavg=vector(mode = "integer", length = 30)
devpreyvar=vector(mode = "integer", length = 30)

for(i in 1:30)
{
  j=i*100
  uniquedrilledavg[j]=mean(subsamples$num_drilled.j.
[which(subsamples$j==j)])
  uniquedrilledvar[j]=sd(subsamples$num_drilled.j.[which(subsamples$j==j)])
  DFavg[j]=mean(subsamples$DF.j.[which(subsamples$j==j)])
  DFvar[j]=sd(subsamples$DF.j.[which(subsamples$j==j)])
  Devavg[j]=mean(subsamples$Dev.j.[which(subsamples$j==j)])
  Devvar[j]=sd(subsamples$Dev.j.[which(subsamples$j==j)])
  devpreyavg[j]=mean(subsamples$dev_numdrilled.j.[which(subsamples$j==j)])
  devpreyvar[j]=sd(subsamples$dev_numdrilled.j.[which(subsamples$j==j)])
  out=data.frame(j,uniquedrilledavg[j],uniquedrilledvar[j],DFavg[j],DFvar[j],Devavg[j],[illegible]
  subsamps1[[i]] <-out
}
case2_0.7even_med<- do.call("rbind",subsamps1)
Drilledsp_avg=case2_0.7even_med$uniquedrilledavg.j.
Drilledsp_var=case2_0.7even_med$uniquedrilledvar.j.
DF_avg=case2_0.7even_med$DFavg.j.
DF_var=case2_0.7even_med$DFvar.j.
DE_avg=case2_0.7even_med$Devavg.j.
DE_var=case2_0.7even_med$Devvar.j.
devpreyavg=case1_0.7even_med$devpreyavg.j.
devpreyvar=case1_0.7even_med$devpreyvar.j.



#############################(Case 2)Simulation for Et=0.7 at high
PIT=0.8---------------
DFtype="High"
Eventype=evenness
Selection="S"
Case=2
PI_T=0.67
prey_actual=5
u=cbind(DFtype,Eventype,Selection,Case)

randomnumber=vector(mode = "integer",length=3000)
raredata=array(0,dim = c(3000,2))
num_species1=vector(mode="integer",length=30)
num_drilled=vector(mode = "integer", length = 30)
DF=vector(mode = "integer", length = 30)
Dev=vector(mode = "integer", length = 30)
dev_numdrilled=vector(mode = "integer", length = 30)
```

```
varevensubset=subset(vareven,select = c(Species,HighD))
subsamps <- list()
subsamps1=list()
for (q in 1:1000)
{
  randomnumber=sample.int(3000,3000, replace = FALSE)
  raredata=varevensubset[randomnumber[1:3000],]
  for(i in 1:30)
  {
    j=i*100
    uniquesp=unique(raredata$Species[1:j])
    num_species1[j]=length(uniquesp)
    uniquedrilled=unique(raredata$Species[which(raredata$HighD[1:j]==1)])
    num_drilled[j]=length(uniquedrilled)
    DF[j]=(sum(raredata$HighD[1:j]))/j
    Dev[j]=PI_T-DF[j]
    dev_numdrilled[j]=prey_actual-num_drilled[j]
    out=data.frame(j,num_species1[j],num_drilled[j],DF[j],Dev[j],dev_numdrilled[j])
    subsamps1[[i]] <-out
  }
  subsamples1 <- do.call("rbind",subsamps1)
  out2=data.frame(q,subsamples1)
  subsamps[[q]]=out2
  subsamples <- do.call("rbind",subsamps)
}
subsamples=as.data.frame(subsamples)



############For calculating average and variance at high
PIT#########################################################
uniquedrilledavg=vector(mode = "integer", length = 30)
uniquedrilledvar=vector(mode = "integer", length = 30)
DFavg=vector(mode = "integer", length = 30)
DFvar=vector(mode = "integer", length = 30)
Devavg=vector(mode = "integer", length = 30)
Devvar=vector(mode = "integer", length = 30)
devpreyavg=vector(mode = "integer", length = 30)
devpreyvar=vector(mode = "integer", length = 30)

for(i in 1:30)
{
  j=i*100
  uniquedrilledavg[j]=mean(subsamples$num_drilled.j.
[which(subsamples$j==j)])
  uniquedrilledvar[j]=sd(subsamples$num_drilled.j.[which(subsamples$j==j)])
  DFavg[j]=mean(subsamples$DF.j.[which(subsamples$j==j)])
  DFvar[j]=sd(subsamples$DF.j.[which(subsamples$j==j)])
  Devavg[j]=mean(subsamples$Dev.j.[which(subsamples$j==j)])
  Devvar[j]=sd(subsamples$Dev.j.[which(subsamples$j==j)])
  devpreyavg[j]=mean(subsamples$dev_numdrilled.j.[which(subsamples$j==j)])
  devpreyvar[j]=sd(subsamples$dev_numdrilled.j.[which(subsamples$j==j)])
  out=data.frame(j,uniquedrilledavg[j],uniquedrilledvar[j],DFavg[j],DFvar[j],Devavg[j],[illegible]
  subsamps1[[i]] <-out
}
case2_0.7even_high <- do.call("rbind",subsamps1)
Drilledsp_avg=case2_0.7even_high$uniquedrilledavg.j.
```

```
Drilledsp_var=case2_0.7even_high$uniquedrilledvar.j.
DF_avg=case2_0.7even_high$DFavg.j.
DF_var=case2_0.7even_high$DFvar.j.
DE_avg=case2_0.7even_high$Devavg.j.
DE_var=case2_0.7even_high$Devvar.j.
devpreyavg=case2_0.7even_high$devpreyavg.j.
devpreyvar=case2_0.7even_high$devpreyvar.j.



##########Re running 0.7 evenness
dataset################################################################
Species=array(0,dim = c(3000,1))
subsamps1=list()
subsamps2=list()

for(i in 1:5)
{
  Species=rep(i,500)
  out=data.frame(Species)
  subsamps1[[i]] <-out
}
identity_species1 <- do.call("rbind",subsamps1)
for(i in 6:30)
{
  Species=rep(i,20)
  out=data.frame(Species)
  subsamps2[[i]] <-out
}
identity_species2 <- do.call("rbind",subsamps2)
identity_species<-rbind(identity_species1,identity_species2)
#############low PIT (0.2) for 3000 individuals##################
d1=c(rep(1,100),rep(0,400))
LowD1=rep(d1,5)
d2=c(rep(1,4),rep(0,16))
LowD2=rep(d2,25)
LowD=c(LowD1,LowD2)
##########medium PIT (0.5) for 3000 individuals#####
md1=c(rep(1,250),rep(0,250))
MedD1=rep(md1,5)
md2=c(rep(1,10),rep(0,10))
MedD2=rep(md2,25)
MedD=c(MedD1,MedD2)
##########high PIT (0.8) for 3000 individuals#####
md1=c(rep(1,400),rep(0,100))
HighD1=rep(md1,5)
md2=c(rep(1,16),rep(0,4))
HighD2=rep(md2,25)
HighD=c(HighD1,HighD2)


Individual=rep(1,3000)
vareven=data.frame(Individual,identity_species,LowD,MedD,HighD)
wideform=as.data.frame.matrix(table(vareven$Individual,vareven$Species))
evenness=(diversity(wideform)/log(specnumber(wideform)))
```

```
#############Case 2:Selective predation for  rare species(25 species(20
individuals each) attacked################
vareven[1:2500,c(3,4,5)]=0 ####creating case3 scenario in E=0.7
Dastaset#####

#####(Case 3)Simulation for Et=0.7 at low PIT=0.2---------------
DFtype="Low"
Eventype=evenness
Selection="S"
Case=3
PI_T=0.03
prey_actual=25
v=cbind(DFtype,Eventype,Selection,Case)


randomnumber=vector(mode = "integer",length=3000)
raredata=array(0,dim = c(3000,2))
num_species1=vector(mode="integer",length=30)
num_drilled=vector(mode = "integer", length = 30)
DF=vector(mode = "integer", length = 30)
Dev=vector(mode = "integer", length = 30)
dev_numdrilled=vector(mode = "integer", length = 30)

varevensubset=subset(vareven,select = c(Species,LowD))
subsamps <- list()
subsamps1=list()
for (q in 1:1000)
{
  randomnumber=sample.int(3000,3000, replace = FALSE)
  raredata=varevensubset[randomnumber[1:3000],]
  for(i in 1:30)
  {
    j=i*100
    uniquesp=unique(raredata$Species[1:j])
    num_species1[j]=length(uniquesp)
    uniquedrilled=unique(raredata$Species[which(raredata$LowD[1:j]==1)])
    num_drilled[j]=length(uniquedrilled)
    DF[j]=(sum(raredata$LowD[1:j]))/j
    Dev[j]=PI_T-DF[j]
    dev_numdrilled[j]=prey_actual-num_drilled[j]
    out=data.frame(j,num_species1[j],num_drilled[j],DF[j],Dev[j],dev_numdrilled[j])
    subsamps1[[i]] <-out
  }
  subsamples1 <- do.call("rbind",subsamps1)
  out2=data.frame(q,subsamples1)
  subsamps[[q]]=out2
  subsamples <- do.call("rbind",subsamps)
}
subsamples=as.data.frame(subsamples)


############For calculating average and variance at low
PIT#########################################################
uniquedrilledavg=vector(mode = "integer", length = 30)
uniquedrilledvar=vector(mode = "integer", length = 30)
DFavg=vector(mode = "integer", length = 30)
DFvar=vector(mode = "integer", length = 30)
```

```
Devavg=vector(mode = "integer", length = 30)
Devvar=vector(mode = "integer", length = 30)
devpreyavg=vector(mode = "integer", length = 30)
devpreyvar=vector(mode = "integer", length = 30)


for(i in 1:30)
{
  j=i*100
  uniquedrilledavg[j]=mean(subsamples$num_drilled.j.
[which(subsamples$j==j)])
  uniquedrilledvar[j]=sd(subsamples$num_drilled.j.[which(subsamples$j==j)])
  DFavg[j]=mean(subsamples$DF.j.[which(subsamples$j==j)])
  DFvar[j]=sd(subsamples$DF.j.[which(subsamples$j==j)])
  Devavg[j]=mean(subsamples$Dev.j.[which(subsamples$j==j)])
  Devvar[j]=sd(subsamples$Dev.j.[which(subsamples$j==j)])
  devpreyavg[j]=mean(subsamples$dev_numdrilled.j.[which(subsamples$j==j)])
  devpreyvar[j]=sd(subsamples$dev_numdrilled.j.[which(subsamples$j==j)])
  out=data.frame(j,uniquedrilledavg[j],uniquedrilledvar[j],DFavg[j],DFvar[j],Devavg[j],[illegible]
  subsamps1[[i]] <-out
}
case3_0.7even_low<- do.call("rbind",subsamps1)
Drilledsp_avg=case3_0.7even_low$uniquedrilledavg.j.
Drilledsp_var=case3_0.7even_low$uniquedrilledvar.j.
DF_avg=case3_0.7even_low$DFavg.j.
DF_var=case3_0.7even_low$DFvar.j.
DE_avg=case3_0.7even_low$Devavg.j.
DE_var=case3_0.7even_low$Devvar.j.
devpreyavg=case3_0.7even_low$devpreyavg.j.
devpreyvar=case3_0.7even_low$devpreyvar.j.



###########################(Case 3)Simulation for Et=0.7 at medium
PIT=0.5---------------
DFtype="Medium"
Eventype=evenness
Selection="S"
Case=3
PI_T=0.08
prey_actual=25
w=cbind(DFtype,Eventype,Selection,Case)

randomnumber=vector(mode = "integer",length=3000)
raredata=array(0,dim = c(3000,2))
num_species1=vector(mode="integer",length=30)
num_drilled=vector(mode = "integer", length = 30)
DF=vector(mode = "integer", length = 30)
varevensubset=subset(vareven,select = c(Species,MedD))
Dev=vector(mode = "integer", length = 30)
dev_numdrilled=vector(mode = "integer", length = 30)

subsamps <- list()
subsamps1=list()
for (q in 1:1000)
{
  randomnumber=sample.int(3000,3000, replace = FALSE)
  raredata=varevensubset[randomnumber[1:3000],]
```

```
  for(i in 1:30)
  {
    j=i*100
    uniquesp=unique(raredata$Species[1:j])
    num_species1[j]=length(uniquesp)
    uniquedrilled=unique(raredata$Species[which(raredata$MedD[1:j]==1)])
    num_drilled[j]=length(uniquedrilled)
    DF[j]=(sum(raredata$MedD[1:j]))/j
    Dev[j]=PI_T-DF[j]
    dev_numdrilled[j]=prey_actual-num_drilled[j]
    out=data.frame(j,num_species1[j],num_drilled[j],DF[j],Dev[j],dev_numdrilled[j])
    subsamps1[[i]] <-out
  }
  subsamples1 <- do.call("rbind",subsamps1)
  out2=data.frame(q,subsamples1)
  subsamps[[q]]=out2
  subsamples <- do.call("rbind",subsamps)
}
subsamples=as.data.frame(subsamples)



############For calculating average and variance for medium
PIT#####################################################
uniquedrilledavg=vector(mode = "integer", length = 30)
uniquedrilledvar=vector(mode = "integer", length = 30)
DFavg=vector(mode = "integer", length = 30)
DFvar=vector(mode = "integer", length = 30)
Devavg=vector(mode = "integer", length = 30)
Devvar=vector(mode = "integer", length = 30)
devpreyavg=vector(mode = "integer", length = 30)
devpreyvar=vector(mode = "integer", length = 30)

for(i in 1:30)
{
  j=i*100
  uniquedrilledavg[j]=mean(subsamples$num_drilled.j.
[which(subsamples$j==j)])
  uniquedrilledvar[j]=sd(subsamples$num_drilled.j.[which(subsamples$j==j)])
  DFavg[j]=mean(subsamples$DF.j.[which(subsamples$j==j)])
  DFvar[j]=sd(subsamples$DF.j.[which(subsamples$j==j)])
  Devavg[j]=mean(subsamples$Dev.j.[which(subsamples$j==j)])
  Devvar[j]=sd(subsamples$Dev.j.[which(subsamples$j==j)])
  devpreyavg[j]=mean(subsamples$dev_numdrilled.j.[which(subsamples$j==j)])
  devpreyvar[j]=sd(subsamples$dev_numdrilled.j.[which(subsamples$j==j)])
  out=data.frame(j,uniquedrilledavg[j],uniquedrilledvar[j],DFavg[j],DFvar[j],Devavg[j],[illegible]
  subsamps1[[i]] <-out
}
case3_0.7even_med<- do.call("rbind",subsamps1)
Drilledsp_avg=case3_0.7even_med$uniquedrilledavg.j.
Drilledsp_var=case3_0.7even_med$uniquedrilledvar.j.
DF_avg=case3_0.7even_med$DFavg.j.
DF_var=case3_0.7even_med$DFvar.j.
DE_avg=case3_0.7even_med$Devavg.j.
DE_var=case3_0.7even_med$Devvar.j.
devpreyavg=case3_0.7even_med$devpreyavg.j.
devpreyvar=case3_0.7even_med$devpreyvar.j.
```

```
############################(Case 3)Simulation for Et=0.7 at high
PIT=0.8---------------
DFtype="High"
Eventype=evenness
Selection="S"
Case=3
PI_T=0.13
prey_actual=25
xi=cbind(DFtype,Eventype,Selection,Case)

randomnumber=vector(mode = "integer",length=3000)
raredata=array(0,dim = c(3000,2))
num_species1=vector(mode="integer",length=30)
num_drilled=vector(mode = "integer", length = 30)
DF=vector(mode = "integer", length = 30)
Dev=vector(mode = "integer", length = 30)
dev_numdrilled=vector(mode = "integer", length = 30)

varevensubset=subset(vareven,select = c(Species,HighD))
subsamps <- list()
subsamps1=list()
for (q in 1:1000)
{
  randomnumber=sample.int(3000,3000, replace = FALSE)
  raredata=varevensubset[randomnumber[1:3000],]
  for(i in 1:30)
  {
    j=i*100
    uniquesp=unique(raredata$Species[1:j])
    num_species1[j]=length(uniquesp)
    uniquedrilled=unique(raredata$Species[which(raredata$HighD[1:j]==1)])
    num_drilled[j]=length(uniquedrilled)
    DF[j]=(sum(raredata$HighD[1:j]))/j
    Dev[j]=PI_T-DF[j]
    dev_numdrilled[j]=prey_actual-num_drilled[j]
    out=data.frame(j,num_species1[j],num_drilled[j],DF[j],Dev[j],dev_numdrilled[j])
    subsamps1[[i]] <-out
  }
  subsamples1 <- do.call("rbind",subsamps1)
  out2=data.frame(q,subsamples1)
  subsamps[[q]]=out2
  subsamples <- do.call("rbind",subsamps)
}
subsamples=as.data.frame(subsamples)



############For calculating average and variance at high
PIT###########################################################
uniquedrilledavg=vector(mode = "integer", length = 30)
uniquedrilledvar=vector(mode = "integer", length = 30)
DFavg=vector(mode = "integer", length = 30)
DFvar=vector(mode = "integer", length = 30)
Devavg=vector(mode = "integer", length = 30)
Devvar=vector(mode = "integer", length = 30)
```

```
devpreyavg=vector(mode = "integer", length = 30)
devpreyvar=vector(mode = "integer", length = 30)

for(i in 1:30)
{
  j=i*100
  uniquedrilledavg[j]=mean(subsamples$num_drilled.j.
[which(subsamples$j==j)])
  uniquedrilledvar[j]=sd(subsamples$num_drilled.j.[which(subsamples$j==j)])
  DFavg[j]=mean(subsamples$DF.j.[which(subsamples$j==j)])
  DFvar[j]=sd(subsamples$DF.j.[which(subsamples$j==j)])
  Devavg[j]=mean(subsamples$Dev.j.[which(subsamples$j==j)])
  Devvar[j]=sd(subsamples$Dev.j.[which(subsamples$j==j)])
  devpreyavg[j]=mean(subsamples$dev_numdrilled.j.[which(subsamples$j==j)])
  devpreyvar[j]=sd(subsamples$dev_numdrilled.j.[which(subsamples$j==j)])
  out=data.frame(j,uniquedrilledavg[j],uniquedrilledvar[j],DFavg[j],DFvar[j],Devavg[j],[illegible]
  subsamps1[[i]] <-out
}
case3_0.7even_high <- do.call("rbind",subsamps1)
Drilledsp_avg=case3_0.7even_high$uniquedrilledavg.j.
Drilledsp_var=case3_0.7even_high$uniquedrilledvar.j.
DF_avg=case3_0.7even_high$DFavg.j.
DF_var=case3_0.7even_high$DFvar.j.
DE_avg=case3_0.7even_high$Devavg.j.
DE_var=case3_0.7even_high$Devvar.j.
devpreyavg=case3_0.7even_high$devpreyavg.j.
devpreyvar=case3_0.7even_high$devpreyvar.j.

############Combining the results of all simulations in
ET=0.7########################################
combined_medeven=rbind(case1_0.7even_low,case1_0.7even_med,case1_0.7even_high,
case2_0.7even_low,case2_0.7even_med,case2_0.7even_high,case3_0.7even_low,case3_0.7even_[illegible]

##############################################################################################[illegible]
#################Creating dataset for Evenness(ET)
=0.5############################################
Species=array(0,dim = c(3000,1))
subsamps1=list()
subsamps2=list()

for(i in 1:3)
{
  Species=rep(i,910)
  out=data.frame(Species)
  subsamps1[[i]] <-out
}
identity_species1 <- do.call("rbind",subsamps1)
for(i in 4:30)
{
  Species=rep(i,10)
  out=data.frame(Species)
  subsamps2[[i]] <-out
}
identity_species2 <- do.call("rbind",subsamps2)
identity_species<-rbind(identity_species1,identity_species2)

#######low predation intensity PIT (0.2) for 3000 individuals#########
d1=c(rep(1,182),rep(0,728))
```

```
LowD1=rep(d1,3)
d2=c(rep(1,2),rep(0,8))
LowD2=rep(d2,27)
LowD=c(LowD1,LowD2)
##########medium predation intensity PIT (0.5) for 3000 individuals#####
md1=c(rep(1,455),rep(0,455))
MedD1=rep(md1,3)
md2=c(rep(1,5),rep(0,5))
MedD2=rep(md2,27)
MedD=c(MedD1,MedD2)
##########high predation intensity PIT (0.8) for 3000 individuals#####
md1=c(rep(1,728),rep(0,182))
HighD1=rep(md1,3)
md2=c(rep(1,8),rep(0,2))
HighD2=rep(md2,27)
HighD=c(HighD1,HighD2)
Individual=rep(1,3000)
vareven=data.frame(Individual,identity_species,LowD,MedD,HighD)
wideform=as.data.frame.matrix(table(vareven$Individual,vareven$Species))
evenness=(diversity(wideform)/log(specnumber(wideform)))




#####(Case 1)Simulation for Et=0.5 at low PIT=0.2---------------
DFtype="Low"
Eventype=evenness
Selection="NS"
Case=1
PI_T=0.2
prey_actual=30
g=cbind(DFtype,Eventype,Selection,Case)

randomnumber=vector(mode = "integer",length=3000)
raredata=array(0,dim = c(3000,2))
num_species1=vector(mode="integer",length=30)
num_drilled=vector(mode = "integer", length = 30)
DF=vector(mode = "integer", length = 30)
varevensubset=subset(vareven,select = c(Species,LowD))
Dev=vector(mode = "integer", length = 30)
dev_numdrilled=vector(mode = "integer", length = 30)

subsamps <- list()
subsamps1=list()
for (q in 1:1000)
{
  randomnumber=sample.int(3000,3000, replace = FALSE)
  raredata=varevensubset[randomnumber[1:3000],]
  for(i in 1:30)
  {
    j=i*100
    uniquesp=unique(raredata$Species[1:j])
    num_species1[j]=length(uniquesp)
    uniquedrilled=unique(raredata$Species[which(raredata$LowD[1:j]==1)])
    num_drilled[j]=length(uniquedrilled)
    DF[j]=(sum(raredata$LowD[1:j]))/j
    Dev[j]=PI_T-DF[j]
    dev_numdrilled[j]=prey_actual-num_drilled[j]
```

```
    out=data.frame(j,num_species1[j],num_drilled[j],DF[j],Dev[j],dev_numdrilled[j])
    subsamps1[[i]] <-out
  }
  subsamples1 <- do.call("rbind",subsamps1)
  out2=data.frame(q,subsamples1)
  subsamps[[q]]=out2
  subsamples <- do.call("rbind",subsamps)
}
subsamples=as.data.frame(subsamples)


############For calculating average and variance at low
PIT#######################################################
uniquedrilledavg=vector(mode = "integer", length = 30)
uniquedrilledvar=vector(mode = "integer", length = 30)
DFavg=vector(mode = "integer", length = 30)
DFvar=vector(mode = "integer", length = 30)
Devavg=vector(mode = "integer", length = 30)
Devvar=vector(mode = "integer", length = 30)
devpreyavg=vector(mode = "integer", length = 30)
devpreyvar=vector(mode = "integer", length = 30)

for(i in 1:30)
{
  j=i*100
  uniquedrilledavg[j]=mean(subsamples$num_drilled.j.
[which(subsamples$j==j)])
  uniquedrilledvar[j]=sd(subsamples$num_drilled.j.[which(subsamples$j==j)])
  DFavg[j]=mean(subsamples$DF.j.[which(subsamples$j==j)])
  DFvar[j]=sd(subsamples$DF.j.[which(subsamples$j==j)])
  Devavg[j]=mean(subsamples$Dev.j.[which(subsamples$j==j)])
  Devvar[j]=sd(subsamples$Dev.j.[which(subsamples$j==j)])
  devpreyavg[j]=mean(subsamples$dev_numdrilled.j.[which(subsamples$j==j)])
  devpreyvar[j]=sd(subsamples$dev_numdrilled.j.[which(subsamples$j==j)])
  out=data.frame(j,uniquedrilledavg[j],uniquedrilledvar[j],DFavg[j],DFvar[j],Devavg[j],[illegible]
  subsamps1[[i]] <-out
}
case1_0.5even_low <- do.call("rbind",subsamps1)
Drilledsp_avg=case1_0.5even_low$uniquedrilledavg.j.
Drilledsp_var=case1_0.5even_low$uniquedrilledvar.j.
DF_avg=case1_0.5even_low$DFavg.j.
DF_var=case1_0.5even_low$DFvar.j.
DE_avg=case1_0.5even_low$Devavg.j.
DE_var=case1_0.5even_low$Devvar.j.
devpreyavg=case1_0.5even_low$devpreyavg.j.
devpreyvar=case1_0.5even_low$devpreyvar.j.




###########################(Case 1)Simulation for Et=0.5 at medium
PIT=0.5---------------
DFtype="Medium"
Eventype=evenness
Selection="NS"
PI_T=0.5
Case=1
prey_actual=30
```

```
h=cbind(DFtype,Eventype,Selection,Case)

randomnumber=vector(mode = "integer",length=3000)
raredata=array(0,dim = c(3000,2))
num_species1=vector(mode="integer",length=30)
num_drilled=vector(mode = "integer", length = 30)
DF=vector(mode = "integer", length = 30)
Dev=vector(mode = "integer", length = 30)
varevensubset=subset(vareven,select = c(Species,MedD))
dev_numdrilled=vector(mode = "integer", length = 30)

subsamps <- list()
subsamps1=list()
for (q in 1:1000)
{
  randomnumber=sample.int(3000,3000, replace = FALSE)
  raredata=varevensubset[randomnumber[1:3000],]
  for(i in 1:30)
  {
    j=i*100
    uniquesp=unique(raredata$Species[1:j])
    num_species1[j]=length(uniquesp)
    uniquedrilled=unique(raredata$Species[which(raredata$MedD[1:j]==1)])
    num_drilled[j]=length(uniquedrilled)
    DF[j]=(sum(raredata$MedD[1:j]))/j
    Dev[j]=PI_T-DF[j]
    dev_numdrilled[j]=prey_actual-num_drilled[j]
    out=data.frame(j,num_species1[j],num_drilled[j],DF[j],Dev[j],dev_numdrilled[j])
    subsamps1[[i]] <-out
  }
  subsamples1 <- do.call("rbind",subsamps1)
  out2=data.frame(q,subsamples1)
  subsamps[[q]]=out2
  subsamples <- do.call("rbind",subsamps)
}
subsamples=as.data.frame(subsamples)

############For calculating average and variance for medium
PIT#########################################################
uniquedrilledavg=vector(mode = "integer", length = 30)
uniquedrilledvar=vector(mode = "integer", length = 30)
DFavg=vector(mode = "integer", length = 30)
DFvar=vector(mode = "integer", length = 30)
Devavg=vector(mode = "integer", length = 30)
Devvar=vector(mode = "integer", length = 30)
devpreyavg=vector(mode = "integer", length = 30)
devpreyvar=vector(mode = "integer", length = 30)

for(i in 1:30)
{
  j=i*100
  uniquedrilledavg[j]=mean(subsamples$num_drilled.j.
[which(subsamples$j==j)])
  uniquedrilledvar[j]=sd(subsamples$num_drilled.j.[which(subsamples$j==j)])
  DFavg[j]=mean(subsamples$DF.j.[which(subsamples$j==j)])
  DFvar[j]=sd(subsamples$DF.j.[which(subsamples$j==j)])
  Devavg[j]=mean(subsamples$Dev.j.[which(subsamples$j==j)])
  Devvar[j]=sd(subsamples$Dev.j.[which(subsamples$j==j)])
```

```
  devpreyavg[j]=mean(subsamples$dev_numdrilled.j.[which(subsamples$j==j)])
  devpreyvar[j]=sd(subsamples$dev_numdrilled.j.[which(subsamples$j==j)])
  out=data.frame(j,uniquedrilledavg[j],uniquedrilledvar[j],DFavg[j],DFvar[j],Devavg[j],
  subsamps1[[i]] <-out
}
case1_0.5even_med <- do.call("rbind",subsamps1)
Drilledsp_avg=case1_0.5even_med$uniquedrilledavg.j.
Drilledsp_var=case1_0.5even_med$uniquedrilledvar.j.
DF_avg=case1_0.5even_med$DFavg.j.
DF_var=case1_0.5even_med$DFvar.j.
DE_avg=case1_0.5even_med$Devavg.j.
DE_var=case1_0.5even_med$Devvar.j.
devpreyavg=case1_0.5even_med$devpreyavg.j.
devpreyvar=case1_0.5even_med$devpreyvar.j.



#############################(Case 1)Simulation for Et=0.5 at high
PIT=0.8---------------
DFtype="High"
Eventype=evenness
Selection="NS"
Case=1
PI_T=0.8
prey_actual=30
i1=cbind(DFtype,Eventype,Selection,Case)

randomnumber=vector(mode = "integer",length=3000)
raredata=array(0,dim = c(3000,2))
num_species1=vector(mode="integer",length=30)
num_drilled=vector(mode = "integer", length = 30)
DF=vector(mode = "integer", length = 30)
varevensubset=subset(vareven,select = c(Species,HighD))
Dev=vector(mode = "integer", length = 30)
dev_numdrilled=vector(mode = "integer", length = 30)

subsamps <- list()
subsamps1=list()
for (q in 1:1000)
{
  randomnumber=sample.int(3000,3000, replace = FALSE)
  raredata=varevensubset[randomnumber[1:3000],]
  for(i in 1:30)
  {
    j=i*100
    uniquesp=unique(raredata$Species[1:j])
    num_species1[j]=length(uniquesp)
    uniquedrilled=unique(raredata$Species[which(raredata$HighD[1:j]==1)])
    num_drilled[j]=length(uniquedrilled)
    DF[j]=(sum(raredata$HighD[1:j]))/j
    Dev[j]=PI_T-DF[j]
    dev_numdrilled[j]=prey_actual-num_drilled[j]
    out=data.frame(j,num_species1[j],num_drilled[j],DF[j],Dev[j],dev_numdrilled[j])
    subsamps1[[i]] <-out
  }
  subsamples1 <- do.call("rbind",subsamps1)
  out2=data.frame(q,subsamples1)
  subsamps[[q]]=out2
```

```
  subsamples <- do.call("rbind",subsamps)
}
subsamples=as.data.frame(subsamples)



########################################################################################
############For calculating average and variance at high
PIT#########################################################
uniquedrilledavg=vector(mode = "integer", length = 30)
uniquedrilledvar=vector(mode = "integer", length = 30)
DFavg=vector(mode = "integer", length = 30)
DFvar=vector(mode = "integer", length = 30)
Devavg=vector(mode = "integer", length = 30)
Devvar=vector(mode = "integer", length = 30)
devpreyavg=vector(mode = "integer", length = 30)
devpreyvar=vector(mode = "integer", length = 30)

for(i in 1:30)
{
  j=i*100
  uniquedrilledavg[j]=mean(subsamples$num_drilled.j.
[which(subsamples$j==j)])
  uniquedrilledvar[j]=sd(subsamples$num_drilled.j.[which(subsamples$j==j)])
  DFavg[j]=mean(subsamples$DF.j.[which(subsamples$j==j)])
  DFvar[j]=sd(subsamples$DF.j.[which(subsamples$j==j)])
  Devavg[j]=mean(subsamples$Dev.j.[which(subsamples$j==j)])
  Devvar[j]=sd(subsamples$Dev.j.[which(subsamples$j==j)])
  devpreyavg[j]=mean(subsamples$dev_numdrilled.j.[which(subsamples$j==j)])
  devpreyvar[j]=sd(subsamples$dev_numdrilled.j.[which(subsamples$j==j)])
  out=data.frame(j,uniquedrilledavg[j],uniquedrilledvar[j],DFavg[j],DFvar[j],Devavg[j],
  subsamps1[[i]] <-out
}
case1_0.5even_high <- do.call("rbind",subsamps1)
Drilledsp_avg=case1_0.5even_high$uniquedrilledavg.j.
Drilledsp_var=case1_0.5even_high$uniquedrilledvar.j.
DF_avg=case1_0.5even_high$DFavg.j.
DF_var=case1_0.5even_high$DFvar.j.
DE_avg=case1_0.5even_high$Devavg.j.
DE_var=case1_0.5even_high$Devvar.j.
devpreyavg=case1_0.5even_high$devpreyavg.j.
devpreyvar=case1_0.5even_high$devpreyvar.j.





#################################Case 2:Selective predation for common
species(first three species(2731 individuals each)
attacked#################################################################
vareven[2731:3000,c(3,4,5)]=0

#####(Case 2)Simulation for Et=0.5 at Low PIT=0.2####################
DFtype="Low"
Eventype=evenness
Selection="S"
PI_T=0.18
Case=2
```

```
prey_actual=3
j1=cbind(DFtype,Eventype,Selection,Case)

randomnumber=vector(mode = "integer",length=3000)
raredata=array(0,dim = c(3000,2))
num_species1=vector(mode="integer",length=30)
num_drilled=vector(mode = "integer", length = 30)
DF=vector(mode = "integer", length = 30)
varevensubset=subset(vareven,select = c(Species,LowD))
Dev=vector(mode = "integer", length = 30)
dev_numdrilled=vector(mode = "integer", length = 30)

subsamps <- list()
subsamps1=list()
for (q in 1:1000)
{
  randomnumber=sample.int(3000,3000, replace = FALSE)
  raredata=varevensubset[randomnumber[1:3000],]
  for(i in 1:30)
  {
    j=i*100
    uniquesp=unique(raredata$Species[1:j])
    num_species1[j]=length(uniquesp)
    uniquedrilled=unique(raredata$Species[which(raredata$LowD[1:j]==1)])
    num_drilled[j]=length(uniquedrilled)
    DF[j]=(sum(raredata$LowD[1:j]))/j
    Dev[j]=PI_T-DF[j]
    dev_numdrilled[j]=prey_actual-num_drilled[j]
    out=data.frame(j,num_species1[j],num_drilled[j],DF[j],Dev[j],dev_numdrilled[j])
    subsamps1[[i]] <-out
  }
  subsamples1 <- do.call("rbind",subsamps1)
  out2=data.frame(q,subsamples1)
  subsamps[[q]]=out2
  subsamples <- do.call("rbind",subsamps)
}
subsamples=as.data.frame(subsamples)


############For calculating average and variance at low
PIT##########################################################
uniquedrilledavg=vector(mode = "integer", length = 30)
uniquedrilledvar=vector(mode = "integer", length = 30)
DFavg=vector(mode = "integer", length = 30)
DFvar=vector(mode = "integer", length = 30)
Devavg=vector(mode = "integer", length = 30)
Devvar=vector(mode = "integer", length = 30)
devpreyavg=vector(mode = "integer", length = 30)
devpreyvar=vector(mode = "integer", length = 30)


for(i in 1:30)
{
  j=i*100
  uniquedrilledavg[j]=mean(subsamples$num_drilled.j.
[which(subsamples$j==j)])
  uniquedrilledvar[j]=sd(subsamples$num_drilled.j.[which(subsamples$j==j)])
  DFavg[j]=mean(subsamples$DF.j.[which(subsamples$j==j)])
```

```
  DFvar[j]=sd(subsamples$DF.j.[which(subsamples$j==j)])
  Devavg[j]=mean(subsamples$Dev.j.[which(subsamples$j==j)])
  Devvar[j]=sd(subsamples$Dev.j.[which(subsamples$j==j)])
  devpreyavg[j]=mean(subsamples$dev_numdrilled.j.[which(subsamples$j==j)])
  devpreyvar[j]=sd(subsamples$dev_numdrilled.j.[which(subsamples$j==j)])
  out=data.frame(j,uniquedrilledavg[j],uniquedrilledvar[j],DFavg[j],DFvar[j],Devavg[j],
  subsamps1[[i]] <-out
}
case2_0.5even_low <- do.call("rbind",subsamps1)
Drilledsp_avg=case2_0.5even_low$uniquedrilledavg.j.
Drilledsp_var=case2_0.5even_low$uniquedrilledvar.j.
DF_avg=case2_0.5even_low$DFavg.j.
DF_var=case2_0.5even_low$DFvar.j.
DE_avg=case2_0.5even_low$Devavg.j.
DE_var=case2_0.5even_low$Devvar.j.
devpreyavg=case2_0.5even_low$devpreyavg.j.
devpreyvar=case2_0.5even_low$devpreyvar.j.




################(Case 2)Simulation for Et=0.5 at medium
PIT=0.5#############################################
DFtype="Medium"
Eventype=evenness
Selection="S"
Case=2
PI_T=0.46
prey_actual=3
k=cbind(DFtype,Eventype,Selection,Case)

randomnumber=vector(mode = "integer",length=3000)
raredata=array(0,dim = c(3000,2))
num_species1=vector(mode="integer",length=30)
num_drilled=vector(mode = "integer", length = 30)
DF=vector(mode = "integer", length = 30)
varevensubset=subset(vareven,select = c(Species,MedD))
Dev=vector(mode = "integer", length = 30)
dev_numdrilled=vector(mode = "integer", length = 30)

subsamps <- list()
subsamps1=list()
for (q in 1:1000)
{
  randomnumber=sample.int(3000,3000, replace = FALSE)
  raredata=varevensubset[randomnumber[1:3000],]
  for(i in 1:30)
  {
    j=i*100
    uniquesp=unique(raredata$Species[1:j])
    num_species1[j]=length(uniquesp)
    uniquedrilled=unique(raredata$Species[which(raredata$MedD[1:j]==1)])
    num_drilled[j]=length(uniquedrilled)
    DF[j]=(sum(raredata$MedD[1:j]))/j
    Dev[j]=PI_T-DF[j]
    dev_numdrilled[j]=prey_actual-num_drilled[j]
    out=data.frame(j,num_species1[j],num_drilled[j],DF[j],Dev[j],dev_numdrilled[j])
    subsamps1[[i]] <-out
```

```
  }
  subsamples1 <- do.call("rbind",subsamps1)
  out2=data.frame(q,subsamples1)
  subsamps[[q]]=out2
  subsamples <- do.call("rbind",subsamps)
}
subsamples=as.data.frame(subsamples)


############For calculating average and variance at medium
PIT#########################################################
uniquedrilledavg=vector(mode = "integer", length = 30)
uniquedrilledvar=vector(mode = "integer", length = 30)
DFavg=vector(mode = "integer", length = 30)
DFvar=vector(mode = "integer", length = 30)
Devavg=vector(mode = "integer", length = 30)
Devvar=vector(mode = "integer", length = 30)
devpreyavg=vector(mode = "integer", length = 30)
devpreyvar=vector(mode = "integer", length = 30)

for(i in 1:30)
{
  j=i*100
  uniquedrilledavg[j]=mean(subsamples$num_drilled.j.
[which(subsamples$j==j)])
  uniquedrilledvar[j]=sd(subsamples$num_drilled.j.[which(subsamples$j==j)])
  DFavg[j]=mean(subsamples$DF.j.[which(subsamples$j==j)])
  DFvar[j]=sd(subsamples$DF.j.[which(subsamples$j==j)])
  Devavg[j]=mean(subsamples$Dev.j.[which(subsamples$j==j)])
  Devvar[j]=sd(subsamples$Dev.j.[which(subsamples$j==j)])
  devpreyavg[j]=mean(subsamples$dev_numdrilled.j.[which(subsamples$j==j)])
  devpreyvar[j]=sd(subsamples$dev_numdrilled.j.[which(subsamples$j==j)])
  out=data.frame(j,uniquedrilledavg[j],uniquedrilledvar[j],DFavg[j],DFvar[j],Devavg[j],[illegible]
  subsamps1[[i]] <-out
}
case2_0.5even_med <- do.call("rbind",subsamps1)
Drilledsp_avg=case2_0.5even_med$uniquedrilledavg.j.
Drilledsp_var=case2_0.5even_med$uniquedrilledvar.j.
DF_avg=case2_0.5even_med$DFavg.j.
DF_var=case2_0.5even_med$DFvar.j.
DE_avg=case2_0.5even_med$Devavg.j.
DE_var=case2_0.5even_med$Devvar.j.
devpreyavg=case2_0.5even_med$devpreyavg.j.
devpreyvar=case2_0.5even_med$devpreyvar.j.




#############################(Case 2)Simulation for Et=0.5 at high
PIT=0.8################################################
DFtype="High"
Eventype=evenness
Selection="S"
Case=2
PI_T=0.73
prey_actual=3
l=cbind(DFtype,Eventype,Selection,Case)
```

```
randomnumber=vector(mode = "integer",length=3000)
raredata=array(0,dim = c(3000,2))
num_species1=vector(mode="integer",length=30)
num_drilled=vector(mode = "integer", length = 30)
DF=vector(mode = "integer", length = 30)
varevensubset=subset(vareven,select = c(Species,HighD))
Dev=vector(mode = "integer", length = 30)
dev_numdrilled=vector(mode = "integer", length = 30)

subsamps <- list()
subsamps1=list()
for (q in 1:1000)
{
  randomnumber=sample.int(3000,3000, replace = FALSE)
  raredata=varevensubset[randomnumber[1:3000],]
  for(i in 1:30)
  {
    j=i*100
    uniquesp=unique(raredata$Species[1:j])
    num_species1[j]=length(uniquesp)
    uniquedrilled=unique(raredata$Species[which(raredata$HighD[1:j]==1)])
    num_drilled[j]=length(uniquedrilled)
    DF[j]=(sum(raredata$HighD[1:j]))/j
    Dev[j]=PI_T-DF[j]
    dev_numdrilled[j]=prey_actual-num_drilled[j]
    out=data.frame(j,num_species1[j],num_drilled[j],DF[j],Dev[j],dev_numdrilled[j])
    subsamps1[[i]] <-out
  }
  subsamples1 <- do.call("rbind",subsamps1)
  out2=data.frame(q,subsamples1)
  subsamps[[q]]=out2
  subsamples <- do.call("rbind",subsamps)
}
subsamples=as.data.frame(subsamples)



#####################################################################################
############For calculating average and variance at high
PIT######################################################
uniquedrilledavg=vector(mode = "integer", length = 30)
uniquedrilledvar=vector(mode = "integer", length = 30)
DFavg=vector(mode = "integer", length = 30)
DFvar=vector(mode = "integer", length = 30)
Devavg=vector(mode = "integer", length = 30)
Devvar=vector(mode = "integer", length = 30)
devpreyavg=vector(mode = "integer", length = 30)
devpreyvar=vector(mode = "integer", length = 30)

for(i in 1:30)
{
  j=i*100
  uniquedrilledavg[j]=mean(subsamples$num_drilled.j.
[which(subsamples$j==j)])
  uniquedrilledvar[j]=sd(subsamples$num_drilled.j.[which(subsamples$j==j)])
  DFavg[j]=mean(subsamples$DF.j.[which(subsamples$j==j)])
  DFvar[j]=sd(subsamples$DF.j.[which(subsamples$j==j)])
  Devavg[j]=mean(subsamples$Dev.j.[which(subsamples$j==j)])
```

```
  Devvar[j]=sd(subsamples$Dev.j.[which(subsamples$j==j)])
  devpreyavg[j]=mean(subsamples$dev_numdrilled.j.[which(subsamples$j==j)])
  devpreyvar[j]=sd(subsamples$dev_numdrilled.j.[which(subsamples$j==j)])
  out=data.frame(j,uniquedrilledavg[j],uniquedrilledvar[j],DFavg[j],DFvar[j],Devavg[j],[illegible]
  subsamps1[[i]] <-out
}
case2_0.5even_high <- do.call("rbind",subsamps1)
Drilledsp_avg=case2_0.5even_high$uniquedrilledavg.j.
Drilledsp_var=case2_0.5even_high$uniquedrilledvar.j.
DF_avg=case2_0.5even_high$DFavg.j.
DF_var=case2_0.5even_high$DFvar.j.
DE_avg=case2_0.5even_high$Devavg.j.
DE_var=case2_0.5even_high$Devvar.j.
devpreyavg=case2_0.5even_high$devpreyavg.j.
devpreyvar=case2_0.5even_high$devpreyvar.j.
#####################################################################################




#################Re-running the ET=0.5 evenness
dataset#############################################
Species=array(0,dim = c(3000,1))
subsamps1=list()
subsamps2=list()

for(i in 1:3)
{
  Species=rep(i,910)
  out=data.frame(Species)
  subsamps1[[i]] <-out
}
identity_species1 <- do.call("rbind",subsamps1)
for(i in 4:30)
{
  Species=rep(i,10)
  out=data.frame(Species)
  subsamps2[[i]] <-out
}
identity_species2 <- do.call("rbind",subsamps2)
identity_species<-rbind(identity_species1,identity_species2)

#######low PIT (0.2) for 3000 individuals#########
d1=c(rep(1,182),rep(0,728))
LowD1=rep(d1,3)
d2=c(rep(1,2),rep(0,8))
LowD2=rep(d2,27)
LowD=c(LowD1,LowD2)
##########medium PIT (0.5) for 3000 individuals#####
md1=c(rep(1,455),rep(0,455))
MedD1=rep(md1,3)
md2=c(rep(1,5),rep(0,5))
MedD2=rep(md2,27)
MedD=c(MedD1,MedD2)
##########high PIT (0.8) for 3000 individuals#####
md1=c(rep(1,728),rep(0,182))
HighD1=rep(md1,3)
```

```
md2=c(rep(1,8),rep(0,2))
HighD2=rep(md2,27)
HighD=c(HighD1,HighD2)
Individual=rep(1,3000)
vareven=data.frame(Individual,identity_species,LowD,MedD,HighD)
wideform=as.data.frame.matrix(table(vareven$Individual,vareven$Species))
evenness=(diversity(wideform)/log(specnumber(wideform)))

########Case 3:Selective predation for rare species(first three species(270
individuals each) attacked#########################
vareven[1:2730,c(3,4,5)]=0
#####(Case 3)Simulation for Et=0.5 at low PIT=0.2####################
DFtype="Low"
Eventype=evenness
Selection="S"
Case=3
PI_T=0.02
prey_actual=27
m=cbind(DFtype,Eventype,Selection,Case)

randomnumber=vector(mode = "integer",length=3000)
raredata=array(0,dim = c(3000,2))
num_species1=vector(mode="integer",length=30)
num_drilled=vector(mode = "integer", length = 30)
DF=vector(mode = "integer", length = 30)
Dev=vector(mode = "integer", length = 30)
varevensubset=subset(vareven,select = c(Species,LowD))
dev_numdrilled=vector(mode = "integer", length = 30)

subsamps <- list()
subsamps1=list()
for (q in 1:1000)
{
  randomnumber=sample.int(3000,3000, replace = FALSE)
  raredata=varevensubset[randomnumber[1:3000],]
  for(i in 1:30)
  {
    j=i*100
    uniquesp=unique(raredata$Species[1:j])
    num_species1[j]=length(uniquesp)
    uniquedrilled=unique(raredata$Species[which(raredata$LowD[1:j]==1)])
    num_drilled[j]=length(uniquedrilled)
    DF[j]=(sum(raredata$LowD[1:j]))/j
    Dev[j]=PI_T-DF[j]
    dev_numdrilled[j]=prey_actual-num_drilled[j]
    out=data.frame(j,num_species1[j],num_drilled[j],DF[j],Dev[j],dev_numdrilled[j])
    subsamps1[[i]] <-out
  }
  subsamples1 <- do.call("rbind",subsamps1)
  out2=data.frame(q,subsamples1)
  subsamps[[q]]=out2
  subsamples <- do.call("rbind",subsamps)
}
subsamples=as.data.frame(subsamples)


###########For calculating average and variance for low
PIT########################################################
```

```
uniquedrilledavg=vector(mode = "integer", length = 30)
uniquedrilledvar=vector(mode = "integer", length = 30)
DFavg=vector(mode = "integer", length = 30)
DFvar=vector(mode = "integer", length = 30)
Devavg=vector(mode = "integer", length = 30)
Devvar=vector(mode = "integer", length = 30)
devpreyavg=vector(mode = "integer", length = 30)
devpreyvar=vector(mode = "integer", length = 30)

for(i in 1:30)
{
  j=i*100
  uniquedrilledavg[j]=mean(subsamples$num_drilled.j.
[which(subsamples$j==j)])
  uniquedrilledvar[j]=sd(subsamples$num_drilled.j.[which(subsamples$j==j)])
  DFavg[j]=mean(subsamples$DF.j.[which(subsamples$j==j)])
  DFvar[j]=sd(subsamples$DF.j.[which(subsamples$j==j)])
  Devavg[j]=mean(subsamples$Dev.j.[which(subsamples$j==j)])
  Devvar[j]=sd(subsamples$Dev.j.[which(subsamples$j==j)])
  devpreyavg[j]=mean(subsamples$dev_numdrilled.j.[which(subsamples$j==j)])
  devpreyvar[j]=sd(subsamples$dev_numdrilled.j.[which(subsamples$j==j)])
  out=data.frame(j,uniquedrilledavg[j],uniquedrilledvar[j],DFavg[j],DFvar[j],Devavg[j],[illegible]
  subsamps1[[i]] <-out
}
case3_0.5even_low <- do.call("rbind",subsamps1)
Drilledsp_avg=case3_0.5even_low$uniquedrilledavg.j.
Drilledsp_var=case3_0.5even_low$uniquedrilledvar.j.
DF_avg=case3_0.5even_low$DFavg.j.
DF_var=case3_0.5even_low$DFvar.j.
DE_avg=case3_0.5even_low$Devavg.j.
DE_var=case3_0.5even_low$Devvar.j.
devpreyavg=case3_0.5even_low$devpreyavg.j.
devpreyvar=case3_0.5even_low$devpreyvar.j.





##############(Case 3)Simulation for Et=0.5 at medium
PIT=0.5###############################################
DFtype="Medium"
Eventype=evenness
Selection="S"
Case=3
PI_T=0.05
prey_actual=27
n=cbind(DFtype,Eventype,Selection,Case)


randomnumber=vector(mode = "integer",length=3000)
raredata=array(0,dim = c(3000,2))
num_species1=vector(mode="integer",length=30)
num_drilled=vector(mode = "integer", length = 30)
DF=vector(mode = "integer", length = 30)
Dev=vector(mode = "integer", length = 30)
varevensubset=subset(vareven,select = c(Species,MedD))
dev_numdrilled=vector(mode = "integer", length = 30)
```

```
subsamps <- list()
subsamps1=list()
for (q in 1:1000)
{
  randomnumber=sample.int(3000,3000, replace = FALSE)
  raredata=varevensubset[randomnumber[1:3000],]
  for(i in 1:30)
  {
    j=i*100
    uniquesp=unique(raredata$Species[1:j])
    num_species1[j]=length(uniquesp)
    uniquedrilled=unique(raredata$Species[which(raredata$MedD[1:j]==1)])
    num_drilled[j]=length(uniquedrilled)
    DF[j]=(sum(raredata$MedD[1:j]))/j
    Dev[j]=PI_T-DF[j]
    dev_numdrilled[j]=prey_actual-num_drilled[j]
    out=data.frame(j,num_species1[j],num_drilled[j],DF[j],Dev[j],dev_numdrilled[j])
    subsamps1[[i]] <-out
  }
  subsamples1 <- do.call("rbind",subsamps1)
  out2=data.frame(q,subsamples1)
  subsamps[[q]]=out2
  subsamples <- do.call("rbind",subsamps)
}
subsamples=as.data.frame(subsamples)




############For calculating average and variance medium
PIT#######################################################
uniquedrilledavg=vector(mode = "integer", length = 30)
uniquedrilledvar=vector(mode = "integer", length = 30)
DFavg=vector(mode = "integer", length = 30)
DFvar=vector(mode = "integer", length = 30)
Devavg=vector(mode = "integer", length = 30)
Devvar=vector(mode = "integer", length = 30)
devpreyavg=vector(mode = "integer", length = 30)
devpreyvar=vector(mode = "integer", length = 30)

for(i in 1:30)
{
  j=i*100
  uniquedrilledavg[j]=mean(subsamples$num_drilled.j.
[which(subsamples$j==j)])
  uniquedrilledvar[j]=sd(subsamples$num_drilled.j.[which(subsamples$j==j)])
  DFavg[j]=mean(subsamples$DF.j.[which(subsamples$j==j)])
  DFvar[j]=sd(subsamples$DF.j.[which(subsamples$j==j)])
  Devavg[j]=mean(subsamples$Dev.j.[which(subsamples$j==j)])
  Devvar[j]=sd(subsamples$Dev.j.[which(subsamples$j==j)])
  devpreyavg[j]=mean(subsamples$dev_numdrilled.j.[which(subsamples$j==j)])
  devpreyvar[j]=sd(subsamples$dev_numdrilled.j.[which(subsamples$j==j)])
  out=data.frame(j,uniquedrilledavg[j],uniquedrilledvar[j],DFavg[j],DFvar[j],Devavg[j],[illegible]
  subsamps1[[i]] <-out
}
case3_0.5even_med <- do.call("rbind",subsamps1)
Drilledsp_avg=case3_0.5even_med$uniquedrilledavg.j.
Drilledsp_var=case3_0.5even_med$uniquedrilledvar.j.
```

```
DF_avg=case3_0.5even_med$DFavg.j.
DF_var=case3_0.5even_med$DFvar.j.
DE_avg=case3_0.5even_med$Devavg.j.
DE_var=case3_0.5even_med$Devvar.j.
devpreyavg=case3_0.5even_med$devpreyavg.j.
devpreyvar=case3_0.5even_med$devpreyvar.j.




#########################(Case 3)Simulation for Et=0.5 at high
PIT=0.2###############################################
DFtype="High"
Eventype=evenness
Selection="S"
Case=3
PI_T=0.07
prey_actual=27
op=cbind(DFtype,Eventype,Selection,Case)

randomnumber=vector(mode = "integer",length=3000)
raredata=array(0,dim = c(3000,2))
num_species1=vector(mode="integer",length=30)
num_drilled=vector(mode = "integer", length = 30)
DF=vector(mode = "integer", length = 30)
Dev=vector(mode = "integer", length = 30)
dev_numdrilled=vector(mode = "integer", length = 30)

varevensubset=subset(vareven,select = c(Species,HighD))
subsamps <- list()
subsamps1=list()


for (q in 1:1000)
{
  randomnumber=sample.int(3000,3000, replace = FALSE)
  raredata=varevensubset[randomnumber[1:3000],]
  for(i in 1:30)
  {
    j=i*100
    uniquesp=unique(raredata$Species[1:j])
    num_species1[j]=length(uniquesp)
    uniquedrilled=unique(raredata$Species[which(raredata$HighD[1:j]==1)])
    num_drilled[j]=length(uniquedrilled)
    DF[j]=(sum(raredata$HighD[1:j]))/j
    Dev[j]=PI_T-DF[j]
    dev_numdrilled[j]=prey_actual-num_drilled[j]
    out=data.frame(j,num_species1[j],num_drilled[j],DF[j],Dev[j],dev_numdrilled[j])
    subsamps1[[i]] <-out
  }
  subsamples1 <- do.call("rbind",subsamps1)
  out2=data.frame(q,subsamples1)
  subsamps[[q]]=out2
  subsamples <- do.call("rbind",subsamps)
}
subsamples=as.data.frame(subsamples)
```

```
#####################################################################################
############For calculating average and variance with high
PT#########################################################
uniquedrilledavg=vector(mode = "integer", length = 30)
uniquedrilledvar=vector(mode = "integer", length = 30)
DFavg=vector(mode = "integer", length = 30)
DFvar=vector(mode = "integer", length = 30)
Devavg=vector(mode = "integer", length = 30)
Devvar=vector(mode = "integer", length = 30)
devpreyavg=vector(mode = "integer", length = 30)
devpreyvar=vector(mode = "integer", length = 30)

for(i in 1:30)
{
  j=i*100
  uniquedrilledavg[j]=mean(subsamples$num_drilled.j.
[which(subsamples$j==j)])
  uniquedrilledvar[j]=sd(subsamples$num_drilled.j.[which(subsamples$j==j)])
  DFavg[j]=mean(subsamples$DF.j.[which(subsamples$j==j)])
  DFvar[j]=sd(subsamples$DF.j.[which(subsamples$j==j)])
  Devavg[j]=mean(subsamples$Dev.j.[which(subsamples$j==j)])
  Devvar[j]=sd(subsamples$Dev.j.[which(subsamples$j==j)])
  devpreyavg[j]=mean(subsamples$dev_numdrilled.j.[which(subsamples$j==j)])
  devpreyvar[j]=sd(subsamples$dev_numdrilled.j.[which(subsamples$j==j)])
  out=data.frame(j,uniquedrilledavg[j],uniquedrilledvar[j],DFavg[j],DFvar[j],Devavg[j],[illegible]
  subsamps1[[i]] <-out
}
case3_0.5even_high <- do.call("rbind",subsamps1)
Drilledsp_avg=case3_0.5even_high$uniquedrilledavg.j.
Drilledsp_var=case3_0.5even_high$uniquedrilledvar.j.
DF_avg=case3_0.5even_high$DFavg.j.
DF_var=case3_0.5even_high$DFvar.j.
DE_avg=case3_0.5even_high$Devavg.j.
DE_var=case3_0.5even_high$Devvar.j.
devpreyavg=case3_0.5even_med$devpreyavg.j.
devpreyvar=case3_0.5even_med$devpreyvar.j.


#####################################################################################
##########combining the results of all simulations at
ET=0.5#################################################################
combined_loweven=rbind(case1_0.5even_low,case1_0.5even_med,case1_0.5even_high,case2_0.5[illegible]


##################Creating dataset for
evennes(ET)=0.2########################################################
Species=array(0,dim = c(3000,1))
subsamps1=list()
subsamps2=list()

for(i in 1:29)
{
  Species=rep(i,10)
  out=data.frame(Species)
  subsamps1[[i]] <-out
}
identity_species1 <- do.call("rbind",subsamps1)
```

```
identity_species2=data.frame(rep(30,2710))
colnames(identity_species2)="Species"

identity_species<-rbind(identity_species1,identity_species2)

#######low PIT(0.2) for 3000 individuals#########
d1=c(rep(1,2),rep(0,8))
LowD1=rep(d1,29)
d2=c(rep(1,542),rep(0,2168))
LowD2=rep(d2,1)
LowD=c(LowD1,LowD2)
##########medium PIT(0.5) for 3000 individuals#####
md1=c(rep(1,5),rep(0,5))
MedD1=rep(md1,29)
md2=c(rep(1,1355),rep(0,1355))
MedD2=rep(md2,1)
MedD=c(MedD1,MedD2)
##########high PIT(0.8) for 3000 individuals#####
md1=c(rep(1,8),rep(0,2))
HighD1=rep(md1,29)
md2=c(rep(1,2168),rep(0,542))
HighD2=rep(md2,1)
HighD=c(HighD1,HighD2)

Individual=rep(1,3000)
vareven=data.frame(Individual,identity_species,LowD,MedD,HighD)
wideform=as.data.frame.matrix(table(vareven$Individual,vareven$Species))
evenness=(diversity(wideform)/log(specnumber(wideform)))




##############(Case 1)Simulation for Et=0.2 at low
PIT=0.2###############################################################

DFtype="Low"
Eventype=evenness
Selection="NS"
Case=1
PI_T=0.2
prey_actual=30
yi=cbind(DFtype,Eventype,Selection,Case)

randomnumber=vector(mode = "integer",length=3000)
raredata=array(0,dim = c(3000,2))
num_species1=vector(mode="integer",length=30)
num_drilled=vector(mode = "integer", length = 30)
DF=vector(mode = "integer", length = 30)
Dev=vector(mode = "integer", length = 30)
varevensubset=subset(vareven,select = c(Species,LowD))
dev_numdrilled=vector(mode = "integer", length = 30)

subsamps <- list()
subsamps1=list()
for (q in 1:1000)
{
  randomnumber=sample.int(3000,3000, replace = FALSE)
  raredata=varevensubset[randomnumber[1:3000],]
```

```
  for(i in 1:30)
  {
    j=i*100
    uniquesp=unique(raredata$Species[1:j])
    num_species1[j]=length(uniquesp)
    uniquedrilled=unique(raredata$Species[which(raredata$LowD[1:j]==1)])
    num_drilled[j]=length(uniquedrilled)
    DF[j]=(sum(raredata$LowD[1:j]))/j
    Dev[j]=PI_T-DF[j]
    dev_numdrilled[j]=prey_actual-num_drilled[j]
    out=data.frame(j,num_species1[j],num_drilled[j],DF[j],Dev[j],dev_numdrilled[j])
    subsamps1[[i]] <-out
  }
  subsamples1 <- do.call("rbind",subsamps1)
  out2=data.frame(q,subsamples1)
  subsamps[[q]]=out2
  subsamples <- do.call("rbind",subsamps)
}
subsamples=as.data.frame(subsamples)



############For calculating average and variance for
PIT#########################################################
uniquedrilledavg=vector(mode = "integer", length = 30)
uniquedrilledvar=vector(mode = "integer", length = 30)
DFavg=vector(mode = "integer", length = 30)
DFvar=vector(mode = "integer", length = 30)
Devavg=vector(mode = "integer", length = 30)
Devvar=vector(mode = "integer", length = 30)
devpreyavg=vector(mode = "integer", length = 30)
devpreyvar=vector(mode = "integer", length = 30)


for(i in 1:30)
{
  j=i*100
  uniquedrilledavg[j]=mean(subsamples$num_drilled.j.
[which(subsamples$j==j)])
  uniquedrilledvar[j]=sd(subsamples$num_drilled.j.[which(subsamples$j==j)])
  DFavg[j]=mean(subsamples$DF.j.[which(subsamples$j==j)])
  DFvar[j]=sd(subsamples$DF.j.[which(subsamples$j==j)])
  Devavg[j]=mean(subsamples$Dev.j.[which(subsamples$j==j)])
  Devvar[j]=sd(subsamples$Dev.j.[which(subsamples$j==j)])
  devpreyavg[j]=mean(subsamples$dev_numdrilled.j.[which(subsamples$j==j)])
  devpreyvar[j]=sd(subsamples$dev_numdrilled.j.[which(subsamples$j==j)])
  out=data.frame(j,uniquedrilledavg[j],uniquedrilledvar[j],DFavg[j],DFvar[j],Devavg[j],[illegible]
  subsamps1[[i]] <-out
}
case1_0.1even_low <- do.call("rbind",subsamps1)
Drilledsp_avg=case1_0.1even_low$uniquedrilledavg.j.
Drilledsp_var=case1_0.1even_low$uniquedrilledvar.j.
DF_avg=case1_0.1even_low$DFavg.j.
DF_var=case1_0.1even_low$DFvar.j.
DE_avg=case1_0.1even_low$Devavg.j.
DE_var=case1_0.1even_low$Devvar.j.
devpreyavg=case1_0.1even_low$devpreyavg.j.
devpreyvar=case1_0.1even_low$devpreyvar.j.
```

```
#####################(Case 1)Simulation for Et=0.2 at medium
PIT=0.5############################################
DFtype="Medium"
Eventype=evenness
Selection="NS"
Case=1
PI_T=0.5
prey_actual=30
z=cbind(DFtype,Eventype,Selection,Case)


randomnumber=vector(mode = "integer",length=3000)
raredata=array(0,dim = c(3000,2))
num_species1=vector(mode="integer",length=30)
num_drilled=vector(mode = "integer", length = 30)
DF=vector(mode = "integer", length = 30)
Dev=vector(mode = "integer", length = 30)
varevensubset=subset(vareven,select = c(Species,MedD))
dev_numdrilled=vector(mode = "integer", length = 30)

subsamps <- list()
subsamps1=list()
for (q in 1:1000)
{
  randomnumber=sample.int(3000,3000, replace = FALSE)
  raredata=varevensubset[randomnumber[1:3000],]
  for(i in 1:30)
  {
    j=i*100
    uniquesp=unique(raredata$Species[1:j])
    num_species1[j]=length(uniquesp)
    uniquedrilled=unique(raredata$Species[which(raredata$MedD[1:j]==1)])
    num_drilled[j]=length(uniquedrilled)
    DF[j]=(sum(raredata$MedD[1:j]))/j
    Dev[j]=PI_T-DF[j]
    dev_numdrilled[j]=prey_actual-num_drilled[j]
    out=data.frame(j,num_species1[j],num_drilled[j],DF[j],Dev[j],dev_numdrilled[j])
    subsamps1[[i]] <-out
  }
  subsamples1 <- do.call("rbind",subsamps1)
  out2=data.frame(q,subsamples1)
  subsamps[[q]]=out2
  subsamples <- do.call("rbind",subsamps)
}
subsamples=as.data.frame(subsamples)



############For calculating average and variance for medium
PIT#########################################################
uniquedrilledavg=vector(mode = "integer", length = 30)
uniquedrilledvar=vector(mode = "integer", length = 30)
DFavg=vector(mode = "integer", length = 30)
DFvar=vector(mode = "integer", length = 30)
Devavg=vector(mode = "integer", length = 30)
Devvar=vector(mode = "integer", length = 30)
devpreyavg=vector(mode = "integer", length = 30)
```

```
devpreyvar=vector(mode = "integer", length = 30)

for(i in 1:30)
{
  j=i*100
  uniquedrilledavg[j]=mean(subsamples$num_drilled.j.
[which(subsamples$j==j)])
  uniquedrilledvar[j]=sd(subsamples$num_drilled.j.[which(subsamples$j==j)])
  DFavg[j]=mean(subsamples$DF.j.[which(subsamples$j==j)])
  DFvar[j]=sd(subsamples$DF.j.[which(subsamples$j==j)])
  Devavg[j]=mean(subsamples$Dev.j.[which(subsamples$j==j)])
  Devvar[j]=sd(subsamples$Dev.j.[which(subsamples$j==j)])
  devpreyavg[j]=mean(subsamples$dev_numdrilled.j.[which(subsamples$j==j)])
  devpreyvar[j]=sd(subsamples$dev_numdrilled.j.[which(subsamples$j==j)])
  out=data.frame(j,uniquedrilledavg[j],uniquedrilledvar[j],DFavg[j],DFvar[j],Devavg[j],
  subsamps1[[i]] <-out
}
case1_0.1even_med <- do.call("rbind",subsamps1)
Drilledsp_avg=case1_0.1even_med$uniquedrilledavg.j.
Drilledsp_var=case1_0.1even_med$uniquedrilledvar.j.
DF_avg=case1_0.1even_med$DFavg.j.
DF_var=case1_0.1even_med$DFvar.j.
DE_avg=case1_0.1even_med$Devavg.j.
DE_var=case1_0.1even_med$Devvar.j.
devpreyavg=case1_0.1even_med$devpreyavg.j.
devpreyvar=case1_0.1even_med$devpreyvar.j.



##############(Case 1)Simulation for Et=0.2 at high
PIT=0.8###############################################
########################################################################
DFtype="High"
Eventype=evenness
Selection="NS"
Case=1
PI_T=0.8
prey_actual=30
ab=cbind(DFtype,Eventype,Selection,Case)

randomnumber=vector(mode = "integer",length=3000)
raredata=array(0,dim = c(3000,2))
num_species1=vector(mode="integer",length=30)
num_drilled=vector(mode = "integer", length = 30)
DF=vector(mode = "integer", length = 30)
Dev=vector(mode = "integer", length = 30)
dev_numdrilled=vector(mode = "integer", length = 30)

varevensubset=subset(vareven,select = c(Species,HighD))
subsamps <- list()
subsamps1=list()
for (q in 1:1000)
{
  randomnumber=sample.int(3000,3000, replace = FALSE)
  raredata=varevensubset[randomnumber[1:3000],]
  for(i in 1:30)
  {
    j=i*100
```

```
    uniquesp=unique(raredata$Species[1:j])
    num_species1[j]=length(uniquesp)
    uniquedrilled=unique(raredata$Species[which(raredata$HighD[1:j]==1)])
    num_drilled[j]=length(uniquedrilled)
    DF[j]=(sum(raredata$HighD[1:j]))/j
    Dev[j]=PI_T-DF[j]
    dev_numdrilled[j]=prey_actual-num_drilled[j]
    out=data.frame(j,num_species1[j],num_drilled[j],DF[j],Dev[j],dev_numdrilled[j])
    subsamps1[[i]] <-out
  }
  subsamples1 <- do.call("rbind",subsamps1)
  out2=data.frame(q,subsamples1)
  subsamps[[q]]=out2
  subsamples <- do.call("rbind",subsamps)
}
subsamples=as.data.frame(subsamples)



########################################################################################
############For calculating average and variance for high
PIT#######################################################
uniquedrilledavg=vector(mode = "integer", length = 30)
uniquedrilledvar=vector(mode = "integer", length = 30)
DFavg=vector(mode = "integer", length = 30)
DFvar=vector(mode = "integer", length = 30)
Devavg=vector(mode = "integer", length = 30)
Devvar=vector(mode = "integer", length = 30)
devpreyavg=vector(mode = "integer", length = 30)
devpreyvar=vector(mode = "integer", length = 30)

for(i in 1:30)
{
  j=i*100
  uniquedrilledavg[j]=mean(subsamples$num_drilled.j.
[which(subsamples$j==j)])
  uniquedrilledvar[j]=sd(subsamples$num_drilled.j.[which(subsamples$j==j)])
  DFavg[j]=mean(subsamples$DF.j.[which(subsamples$j==j)])
  DFvar[j]=sd(subsamples$DF.j.[which(subsamples$j==j)])
  Devavg[j]=mean(subsamples$Dev.j.[which(subsamples$j==j)])
  Devvar[j]=sd(subsamples$Dev.j.[which(subsamples$j==j)])
  devpreyavg[j]=mean(subsamples$dev_numdrilled.j.[which(subsamples$j==j)])
  devpreyvar[j]=sd(subsamples$dev_numdrilled.j.[which(subsamples$j==j)])
  out=data.frame(j,uniquedrilledavg[j],uniquedrilledvar[j],DFavg[j],DFvar[j],Devavg[j],[illegible]
  subsamps1[[i]] <-out
}
case1_0.1even_high <- do.call("rbind",subsamps1)
Drilledsp_avg=case1_0.1even_high$uniquedrilledavg.j.
Drilledsp_var=case1_0.1even_high$uniquedrilledvar.j.
DF_avg=case1_0.1even_high$DFavg.j.
DF_var=case1_0.1even_high$DFvar.j.
DE_avg=case1_0.1even_high$Devavg.j.
DE_var=case1_0.1even_high$Devvar.j.
devpreyavg=case1_0.1even_high$devpreyavg.j.
devpreyvar=case1_0.1even_high$devpreyvar.j.
```

```
#############CASE 2: Selective drilling for common species(only one species
with 2710 individuals drilled)#################
vareven[1:290,c(3,4,5)]=0

#####(Case 2)Simulation for Et=0.2 at low PIT=0.2####################
DFtype="Low"
Eventype=evenness
Selection="S"
Case=2
PI_T=0.18
prey_actual=1
ac=cbind(DFtype,Eventype,Selection,Case)

randomnumber=vector(mode = "integer",length=3000)
raredata=array(0,dim = c(3000,2))
num_species1=vector(mode="integer",length=30)
num_drilled=vector(mode = "integer", length = 30)
DF=vector(mode = "integer", length = 30)
Dev=vector(mode = "integer", length = 30)
dev_numdrilled=vector(mode = "integer", length = 30)

varevensubset=subset(vareven,select = c(Species,LowD))
subsamps <- list()
subsamps1=list()
for (q in 1:1000)
{
  randomnumber=sample.int(3000,3000, replace = FALSE)
  raredata=varevensubset[randomnumber[1:3000],]
  for(i in 1:30)
  {
    j=i*100
    uniquesp=unique(raredata$Species[1:j])
    num_species1[j]=length(uniquesp)
    uniquedrilled=unique(raredata$Species[which(raredata$LowD[1:j]==1)])
    num_drilled[j]=length(uniquedrilled)
    DF[j]=(sum(raredata$LowD[1:j]))/j
    Dev[j]=PI_T-DF[j]
    dev_numdrilled[j]=prey_actual-num_drilled[j]
    out=data.frame(j,num_species1[j],num_drilled[j],DF[j],Dev[j],dev_numdrilled[j])
    subsamps1[[i]] <-out
  }
  subsamples1 <- do.call("rbind",subsamps1)
  out2=data.frame(q,subsamples1)
  subsamps[[q]]=out2
  subsamples <- do.call("rbind",subsamps)
}
subsamples=as.data.frame(subsamples)




############For calculating average and variance for low
PIT#########################################################
uniquedrilledavg=vector(mode = "integer", length = 30)
uniquedrilledvar=vector(mode = "integer", length = 30)
DFavg=vector(mode = "integer", length = 30)
DFvar=vector(mode = "integer", length = 30)
Devavg=vector(mode = "integer", length = 30)
```

```
Devvar=vector(mode = "integer", length = 30)
devpreyavg=vector(mode = "integer", length = 30)
devpreyvar=vector(mode = "integer", length = 30)

for(i in 1:30)
{
  j=i*100
  uniquedrilledavg[j]=mean(subsamples$num_drilled.j.
[which(subsamples$j==j)])
  uniquedrilledvar[j]=sd(subsamples$num_drilled.j.[which(subsamples$j==j)])
  DFavg[j]=mean(subsamples$DF.j.[which(subsamples$j==j)])
  DFvar[j]=sd(subsamples$DF.j.[which(subsamples$j==j)])
  Devavg[j]=mean(subsamples$Dev.j.[which(subsamples$j==j)])
  Devvar[j]=sd(subsamples$Dev.j.[which(subsamples$j==j)])
  devpreyavg[j]=mean(subsamples$dev_numdrilled.j.[which(subsamples$j==j)])
  devpreyvar[j]=sd(subsamples$dev_numdrilled.j.[which(subsamples$j==j)])
  out=data.frame(j,uniquedrilledavg[j],uniquedrilledvar[j],DFavg[j],DFvar[j],Devavg[j],
  subsamps1[[i]] <-out
}
case2_0.1even_low <- do.call("rbind",subsamps1)
Drilledsp_avg=case2_0.1even_low$uniquedrilledavg.j.
Drilledsp_var=case2_0.1even_low$uniquedrilledvar.j.
DF_avg=case2_0.1even_low$DFavg.j.
DF_var=case2_0.1even_low$DFvar.j.
DE_avg=case2_0.1even_low$Devavg.j.
DE_var=case2_0.1even_low$Devvar.j.
devpreyavg=case2_0.1even_low$devpreyavg.j.
devpreyvar=case2_0.1even_low$devpreyvar.j.


#########################(Case 2)Simulation for Et=0.2 at medium
PIT=0.5#############################################
DFtype="Medium"
Eventype=evenness
Selection="S"
PI_T=0.45
Case=2
prey_actual=1
ad=cbind(DFtype,Eventype,Selection,Case)


randomnumber=vector(mode = "integer",length=3000)
raredata=array(0,dim = c(3000,2))
num_species1=vector(mode="integer",length=30)
num_drilled=vector(mode = "integer", length = 30)
DF=vector(mode = "integer", length = 30)
varevensubset=subset(vareven,select = c(Species,MedD))
Dev=vector(mode = "integer", length = 30)
dev_numdrilled=vector(mode = "integer", length = 30)

subsamps <- list()
subsamps1=list()
for (q in 1:1000)
{
  randomnumber=sample.int(3000,3000, replace = FALSE)
  raredata=varevensubset[randomnumber[1:3000],]
  for(i in 1:30)
  {
```

```
    j=i*100
    uniquesp=unique(raredata$Species[1:j])
    num_species1[j]=length(uniquesp)
    uniquedrilled=unique(raredata$Species[which(raredata$MedD[1:j]==1)])
    num_drilled[j]=length(uniquedrilled)
    DF[j]=(sum(raredata$MedD[1:j]))/j
    Dev[j]=PI_T-DF[j]
    dev_numdrilled[j]=prey_actual-num_drilled[j]
    out=data.frame(j,num_species1[j],num_drilled[j],DF[j],Dev[j],dev_numdrilled[j])
    subsamps1[[i]] <-out
  }
  subsamples1 <- do.call("rbind",subsamps1)
  out2=data.frame(q,subsamples1)
  subsamps[[q]]=out2
  subsamples <- do.call("rbind",subsamps)
}
subsamples=as.data.frame(subsamples)

############For calculating average and variance for medium
PIT##########################################################
uniquedrilledavg=vector(mode = "integer", length = 30)
uniquedrilledvar=vector(mode = "integer", length = 30)
DFavg=vector(mode = "integer", length = 30)
DFvar=vector(mode = "integer", length = 30)
Devavg=vector(mode = "integer", length = 30)
Devvar=vector(mode = "integer", length = 30)
devpreyavg=vector(mode = "integer", length = 30)
devpreyvar=vector(mode = "integer", length = 30)

for(i in 1:30)
{
  j=i*100
  uniquedrilledavg[j]=mean(subsamples$num_drilled.j.
[which(subsamples$j==j)])
  uniquedrilledvar[j]=sd(subsamples$num_drilled.j.[which(subsamples$j==j)])
  DFavg[j]=mean(subsamples$DF.j.[which(subsamples$j==j)])
  DFvar[j]=sd(subsamples$DF.j.[which(subsamples$j==j)])
  Devavg[j]=mean(subsamples$Dev.j.[which(subsamples$j==j)])
  Devvar[j]=sd(subsamples$Dev.j.[which(subsamples$j==j)])
  devpreyavg[j]=mean(subsamples$dev_numdrilled.j.[which(subsamples$j==j)])
  devpreyvar[j]=sd(subsamples$dev_numdrilled.j.[which(subsamples$j==j)])
  out=data.frame(j,uniquedrilledavg[j],uniquedrilledvar[j],DFavg[j],DFvar[j],Devavg[j],[illegible]
  subsamps1[[i]] <-out
}
case2_0.1even_med <- do.call("rbind",subsamps1)
Drilledsp_avg=case2_0.1even_med$uniquedrilledavg.j.
Drilledsp_var=case2_0.1even_med$uniquedrilledvar.j.
DF_avg=case2_0.1even_med$DFavg.j.
DF_var=case2_0.1even_med$DFvar.j.
DE_avg=case2_0.1even_med$Devavg.j.
DE_var=case2_0.1even_med$Devvar.j.
devpreyavg=case2_0.1even_med$devpreyavg.j.
devpreyvar=case2_0.1even_med$devpreyvar.j.



#############(Case 2)Simulation for Et=0.2 at high
PIT=0.8##########################
```

```
DFtype="High"
Eventype=evenness
Selection="S"
Case=2
PI_T=0.72
prey_actual=1
ae=cbind(DFtype,Eventype,Selection,Case)

randomnumber=vector(mode = "integer",length=3000)
raredata=array(0,dim = c(3000,2))
num_species1=vector(mode="integer",length=30)
num_drilled=vector(mode = "integer", length = 30)
DF=vector(mode = "integer", length = 30)
varevensubset=subset(vareven,select = c(Species,HighD))
Dev=vector(mode = "integer", length = 30)
dev_numdrilled=vector(mode = "integer", length = 30)

subsamps <- list()
subsamps1=list()
for (q in 1:1000)
{
  randomnumber=sample.int(3000,3000, replace = FALSE)
  raredata=varevensubset[randomnumber[1:3000],]
  for(i in 1:30)
  {
    j=i*100
    uniquesp=unique(raredata$Species[1:j])
    num_species1[j]=length(uniquesp)
    uniquedrilled=unique(raredata$Species[which(raredata$HighD[1:j]==1)])
    num_drilled[j]=length(uniquedrilled)
    DF[j]=(sum(raredata$HighD[1:j]))/j
    Dev[j]=PI_T-DF[j]
    dev_numdrilled[j]=prey_actual-num_drilled[j]
    out=data.frame(j,num_species1[j],num_drilled[j],DF[j],Dev[j],dev_numdrilled[j])
    subsamps1[[i]] <-out
  }
  subsamples1 <- do.call("rbind",subsamps1)
  out2=data.frame(q,subsamples1)
  subsamps[[q]]=out2
  subsamples <- do.call("rbind",subsamps)
}
subsamples=as.data.frame(subsamples)


#####################################################################################
############For calculating average and variance for high
PIT#########################################################
uniquedrilledavg=vector(mode = "integer", length = 30)
uniquedrilledvar=vector(mode = "integer", length = 30)
DFavg=vector(mode = "integer", length = 30)
DFvar=vector(mode = "integer", length = 30)
Devavg=vector(mode = "integer", length = 30)
Devvar=vector(mode = "integer", length = 30)
devpreyavg=vector(mode = "integer", length = 30)
devpreyvar=vector(mode = "integer", length = 30)

for(i in 1:30)
```

```
{
  j=i*100
  uniquedrilledavg[j]=mean(subsamples$num_drilled.j.
[which(subsamples$j==j)])
  uniquedrilledvar[j]=sd(subsamples$num_drilled.j.[which(subsamples$j==j)])
  DFavg[j]=mean(subsamples$DF.j.[which(subsamples$j==j)])
  DFvar[j]=sd(subsamples$DF.j.[which(subsamples$j==j)])
  Devavg[j]=mean(subsamples$Dev.j.[which(subsamples$j==j)])
  Devvar[j]=sd(subsamples$Dev.j.[which(subsamples$j==j)])
  devpreyavg[j]=mean(subsamples$dev_numdrilled.j.[which(subsamples$j==j)])
  devpreyvar[j]=sd(subsamples$dev_numdrilled.j.[which(subsamples$j==j)])
  out=data.frame(j,uniquedrilledavg[j],uniquedrilledvar[j],DFavg[j],DFvar[j],Devavg[j],[illegible]
  subsamps1[[i]] <-out
}
case2_0.1even_high <- do.call("rbind",subsamps1)
Drilledsp_avg=case2_0.1even_high$uniquedrilledavg.j.
Drilledsp_var=case2_0.1even_high$uniquedrilledvar.j.
DF_avg=case2_0.1even_high$DFavg.j.
DF_var=case2_0.1even_high$DFvar.j.
DE_avg=case2_0.1even_high$Devavg.j.
DE_var=case2_0.1even_high$Devvar.j.
devpreyavg=case2_0.1even_high$devpreyavg.j.
devpreyvar=case2_0.1even_high$devpreyvar.j.




###############Re running the ET=0.2 evenness
dataset#################################################################
Species=array(0,dim = c(3000,1))
subsamps1=list()
subsamps2=list()

for(i in 1:29)
{
  Species=rep(i,10)
  out=data.frame(Species)
  subsamps1[[i]] <-out
}
identity_species1 <- do.call("rbind",subsamps1)
identity_species2=data.frame(rep(30,2710))
colnames(identity_species2)="Species"

identity_species<-rbind(identity_species1,identity_species2)

#######low PIT (0.2) for 3000 individuals#########
d1=c(rep(1,2),rep(0,8))
LowD1=rep(d1,29)
d2=c(rep(1,542),rep(0,2168))
LowD2=rep(d2,1)
LowD=c(LowD1,LowD2)
##########medium PIT (0.5) for 3000 individuals#####
md1=c(rep(1,5),rep(0,5))
MedD1=rep(md1,29)
md2=c(rep(1,1355),rep(0,1355))
MedD2=rep(md2,1)
MedD=c(MedD1,MedD2)
##########high PIT (0.8) for 3000 individuals#####
```

```
md1=c(rep(1,8),rep(0,2))
HighD1=rep(md1,29)
md2=c(rep(1,2168),rep(0,542))
HighD2=rep(md2,1)
HighD=c(HighD1,HighD2)

Individual=rep(1,3000)
vareven=data.frame(Individual,identity_species,LowD,MedD,HighD)
wideform=as.data.frame.matrix(table(vareven$Individual,vareven$Species))
evenness=(diversity(wideform)/log(specnumber(wideform)))




#############Case 3:Selective drilling for rare species(sp 1-29) with 10
individuals each drilled)#################
vareven[291:3000,c(3,4,5)]=0


#####(Case 3)Simulation for Et=0.2 at low PIT=0.2####################
DFtype="Low"
Eventype=evenness
Selection="S"
Case=3
PI_T=0.02
prey_actual=29
af=cbind(DFtype,Eventype,Selection,Case)

randomnumber=vector(mode = "integer",length=3000)
raredata=array(0,dim = c(3000,2))
num_species1=vector(mode="integer",length=30)
num_drilled=vector(mode = "integer", length = 30)
DF=vector(mode = "integer", length = 30)
Dev=vector(mode = "integer", length = 30)
dev_numdrilled=vector(mode = "integer", length = 30)

varevensubset=subset(vareven,select = c(Species,LowD))
subsamps <- list()
subsamps1=list()
for (q in 1:1000)
{
  randomnumber=sample.int(3000,3000, replace = FALSE)
  raredata=varevensubset[randomnumber[1:3000],]
  for(i in 1:30)
  {
    j=i*100
    uniquesp=unique(raredata$Species[1:j])
    num_species1[j]=length(uniquesp)
    uniquedrilled=unique(raredata$Species[which(raredata$LowD[1:j]==1)])
    num_drilled[j]=length(uniquedrilled)
    DF[j]=(sum(raredata$LowD[1:j]))/j
    Dev[j]=PI_T-DF[j]
    dev_numdrilled[j]=prey_actual-num_drilled[j]
    out=data.frame(j,num_species1[j],num_drilled[j],DF[j],Dev[j],dev_numdrilled[j])
    subsamps1[[i]] <-out
  }
  subsamples1 <- do.call("rbind",subsamps1)
  out2=data.frame(q,subsamples1)
```

```
  subsamps[[q]]=out2
  subsamples <- do.call("rbind",subsamps)
}
subsamples=as.data.frame(subsamples)




############For calculating average and variance for low
PIT######################################################
uniquedrilledavg=vector(mode = "integer", length = 30)
uniquedrilledvar=vector(mode = "integer", length = 30)
DFavg=vector(mode = "integer", length = 30)
DFvar=vector(mode = "integer", length = 30)
Devavg=vector(mode = "integer", length = 30)
Devvar=vector(mode = "integer", length = 30)
devpreyavg=vector(mode = "integer", length = 30)
devpreyvar=vector(mode = "integer", length = 30)

for(i in 1:30)
{
  j=i*100
  uniquedrilledavg[j]=mean(subsamples$num_drilled.j.
[which(subsamples$j==j)])
  uniquedrilledvar[j]=sd(subsamples$num_drilled.j.[which(subsamples$j==j)])
  DFavg[j]=mean(subsamples$DF.j.[which(subsamples$j==j)])
  DFvar[j]=sd(subsamples$DF.j.[which(subsamples$j==j)])
  Devavg[j]=mean(subsamples$Dev.j.[which(subsamples$j==j)])
  Devvar[j]=sd(subsamples$Dev.j.[which(subsamples$j==j)])
  devpreyavg[j]=mean(subsamples$dev_numdrilled.j.[which(subsamples$j==j)])
  devpreyvar[j]=sd(subsamples$dev_numdrilled.j.[which(subsamples$j==j)])
  out=data.frame(j,uniquedrilledavg[j],uniquedrilledvar[j],DFavg[j],DFvar[j],Devavg[j],[illegible]
  subsamps1[[i]] <-out
}
case3_0.1even_low <- do.call("rbind",subsamps1)
Drilledsp_avg=case3_0.1even_low$uniquedrilledavg.j.
Drilledsp_var=case3_0.1even_low$uniquedrilledvar.j.
DF_avg=case3_0.1even_low$DFavg.j.
DF_var=case3_0.1even_low$DFvar.j.
DE_avg=case3_0.1even_low$Devavg.j.
DE_var=case3_0.1even_low$Devvar.j.
devpreyavg=case3_0.1even_low$devpreyavg.j.
devpreyvar=case3_0.1even_low$devpreyvar.j.


#######################(Case 3)Simulation for Et=0.2 at medium
PIT=0.2###############################################
DFtype="Medium"
Eventype=evenness
Selection="S"
Case=3
PI_T=0.05
prey_actual=29
ag=cbind(DFtype,Eventype,Selection,Case)


randomnumber=vector(mode = "integer",length=3000)
raredata=array(0,dim = c(3000,2))
```

```
num_species1=vector(mode="integer",length=30)
num_drilled=vector(mode = "integer", length = 30)
DF=vector(mode = "integer", length = 30)
Dev=vector(mode = "integer", length = 30)
dev_numdrilled=vector(mode = "integer", length = 30)

varevensubset=subset(vareven,select = c(Species,MedD))
subsamps <- list()
subsamps1=list()
for (q in 1:1000)
{
  randomnumber=sample.int(3000,3000, replace = FALSE)
  raredata=varevensubset[randomnumber[1:3000],]
  for(i in 1:30)
  {
    j=i*100
    uniquesp=unique(raredata$Species[1:j])
    num_species1[j]=length(uniquesp)
    uniquedrilled=unique(raredata$Species[which(raredata$MedD[1:j]==1)])
    num_drilled[j]=length(uniquedrilled)
    DF[j]=(sum(raredata$MedD[1:j]))/j
    Dev[j]=PI_T-DF[j]
    dev_numdrilled[j]=prey_actual-num_drilled[j]
    out=data.frame(j,num_species1[j],num_drilled[j],DF[j],Dev[j],dev_numdrilled[j])
    subsamps1[[i]] <-out
  }
  subsamples1 <- do.call("rbind",subsamps1)
  out2=data.frame(q,subsamples1)
  subsamps[[q]]=out2
  subsamples <- do.call("rbind",subsamps)
}
subsamples=as.data.frame(subsamples)


############For calculating average and variance for medium
PIT##########################################################
uniquedrilledavg=vector(mode = "integer", length = 30)
uniquedrilledvar=vector(mode = "integer", length = 30)
DFavg=vector(mode = "integer", length = 30)
DFvar=vector(mode = "integer", length = 30)
Devavg=vector(mode = "integer", length = 30)
Devvar=vector(mode = "integer", length = 30)
devpreyavg=vector(mode = "integer", length = 30)
devpreyvar=vector(mode = "integer", length = 30)

for(i in 1:30)
{
  j=i*100
  uniquedrilledavg[j]=mean(subsamples$num_drilled.j.
[which(subsamples$j==j)])
  uniquedrilledvar[j]=sd(subsamples$num_drilled.j.[which(subsamples$j==j)])
  DFavg[j]=mean(subsamples$DF.j.[which(subsamples$j==j)])
  DFvar[j]=sd(subsamples$DF.j.[which(subsamples$j==j)])
  Devavg[j]=mean(subsamples$Dev.j.[which(subsamples$j==j)])
  Devvar[j]=sd(subsamples$Dev.j.[which(subsamples$j==j)])
  devpreyavg[j]=mean(subsamples$dev_numdrilled.j.[which(subsamples$j==j)])
  devpreyvar[j]=sd(subsamples$dev_numdrilled.j.[which(subsamples$j==j)])
  out=data.frame(j,uniquedrilledavg[j],uniquedrilledvar[j],DFavg[j],DFvar[j],Devavg[j],
```

```
  subsamps1[[i]] <-out
}
case3_0.1even_med <- do.call("rbind",subsamps1)
Drilledsp_avg=case3_0.1even_med$uniquedrilledavg.j.
Drilledsp_var=case3_0.1even_med$uniquedrilledvar.j.
DF_avg=case3_0.1even_med$DFavg.j.
DF_var=case3_0.1even_med$DFvar.j.
DE_avg=case3_0.1even_med$Devavg.j.
DE_var=case3_0.1even_med$Devvar.j.
devpreyavg=case3_0.1even_med$devpreyavg.j.
devpreyvar=case3_0.1even_med$devpreyvar.j.



###########(Case 3)Simulation for Et=0.2 at high
PIT=0.8##############################################

DFtype="High"
Eventype=evenness
Selection="S"
Case=3
PI_T=0.08
prey_actual=29
ah=cbind(DFtype,Eventype,Selection,Case)

randomnumber=vector(mode = "integer",length=3000)
raredata=array(0,dim = c(3000,2))
num_species1=vector(mode="integer",length=30)
num_drilled=vector(mode = "integer", length = 30)
DF=vector(mode = "integer", length = 30)
Dev=vector(mode = "integer", length = 30)
dev_numdrilled=vector(mode = "integer", length = 30)

varevensubset=subset(vareven,select = c(Species,HighD))
subsamps <- list()
subsamps1=list()
for (q in 1:1000)
{
  randomnumber=sample.int(3000,3000, replace = FALSE)
  raredata=varevensubset[randomnumber[1:3000],]
  for(i in 1:30)
  {
    j=i*100
    uniquesp=unique(raredata$Species[1:j])
    num_species1[j]=length(uniquesp)
    uniquedrilled=unique(raredata$Species[which(raredata$HighD[1:j]==1)])
    num_drilled[j]=length(uniquedrilled)
    DF[j]=(sum(raredata$HighD[1:j]))/j
    Dev[j]=PI_T-DF[j]
    dev_numdrilled[j]=prey_actual-num_drilled[j]
    out=data.frame(j,num_species1[j],num_drilled[j],DF[j],Dev[j],dev_numdrilled[j])
    subsamps1[[i]] <-out
  }
  subsamples1 <- do.call("rbind",subsamps1)
  out2=data.frame(q,subsamples1)
  subsamps[[q]]=out2
  subsamples <- do.call("rbind",subsamps)
}
```

```
subsamples=as.data.frame(subsamples)



#######################################################################################
############For calculating average and variance for high
PIT#######################################################
uniquedrilledavg=vector(mode = "integer", length = 30)
uniquedrilledvar=vector(mode = "integer", length = 30)
DFavg=vector(mode = "integer", length = 30)
DFvar=vector(mode = "integer", length = 30)
Devavg=vector(mode = "integer", length = 30)
Devvar=vector(mode = "integer", length = 30)
devpreyavg=vector(mode = "integer", length = 30)
devpreyvar=vector(mode = "integer", length = 30)

for(i in 1:30)
{
  j=i*100
  uniquedrilledavg[j]=mean(subsamples$num_drilled.j.
[which(subsamples$j==j)])
  uniquedrilledvar[j]=sd(subsamples$num_drilled.j.[which(subsamples$j==j)])
  DFavg[j]=mean(subsamples$DF.j.[which(subsamples$j==j)])
  DFvar[j]=sd(subsamples$DF.j.[which(subsamples$j==j)])
  Devavg[j]=mean(subsamples$Dev.j.[which(subsamples$j==j)])
  Devvar[j]=sd(subsamples$Dev.j.[which(subsamples$j==j)])
  devpreyavg[j]=mean(subsamples$dev_numdrilled.j.[which(subsamples$j==j)])
  devpreyvar[j]=sd(subsamples$dev_numdrilled.j.[which(subsamples$j==j)])
  out=data.frame(j,uniquedrilledavg[j],uniquedrilledvar[j],DFavg[j],DFvar[j],Devavg[j],[illegible]
  subsamps1[[i]] <-out
}
case3_0.1even_high <- do.call("rbind",subsamps1)
Drilledsp_avg=case3_0.1even_high$uniquedrilledavg.j.
Drilledsp_var=case3_0.1even_high$uniquedrilledvar.j.
DF_avg=case3_0.1even_high$DFavg.j.
DF_var=case3_0.1even_high$DFvar.j.
DE_avg=case3_0.1even_high$Devavg.j.
DE_var=case3_0.1even_high$Devvar.j.
devpreyavg=case3_0.1even_high$devpreyavg.j.
devpreyvar=case3_0.1even_high$devpreyvar.j.



##########combining the results of all simulations for
ET=0.2##################
combined_vloweven=rbind(case1_0.1even_low,case1_0.1even_med,case1_0.1even_high,
case2_0.1even_low,case2_0.1even_med,case2_0.1even_high,case3_0.1even_low,case3_0.1even_[illegible]

##########################################################################################[illegible]
##########combining all the assemblages
together#####################################################
combineddata2=rbind(combined_max_even_2,combined_medeven,combined_loweven,combined_vlow[illegible]
#write.table(combineddata2,file="combined2022_mod_2.csv")
combinedata=read.csv(file="combined2022_mod_2.csv",header=T)
#combinedata$Case <- replace(combinedata$Case, (combinedata$Eventype ==
"")&(combinedata$Selection=="S"), "1")####make selective drilling in
maxeven case 1 from case 0#######
require(dplyr)
```

```
require(ggplot2)
combinedata$Eventype=round(combinedata[,10],1)
neworderevenness <- c("0.2","0.5","0.7","1")
combined2 <-
arrange(transform(combinedata,Eventype=factor(Eventype,levels=neworderevenness)),Eventy
the data to sequentially plot low , medium high in ggplot2###############

neworder <- c("Low","Medium","High")
combined2 <-
arrange(transform(combinedata,DFtype=factor(DFtype,levels=neworder)),DFtype)########rea
the data to sequentially plot low , medium high in ggplot2###############

combined2=combinedata
Drilledsp_avg=combined2$uniquedrilledavg.j.
Drilledsp_var=combined2$uniquedrilledvar.j.
DF_avg=combined2$DFavg.j.
DF_var=combined2$DFvar.j.
DE_avg=combined2$Devavg.j.
DE_var=combined2$Devvar.j.

j=combined2$j
DFtype=combined2$DFtype
#Eventype=combined2$Eventype
#Case=combined2$Case
factor_Eventype=factor(Eventype)
DFtype=factor(DFtype)
######Figure 4##################################################
ggplot(combined2,aes(j,DF_avg,gcoroup=factor_Eventype,color=factor_Eventype))
+  xlab("Sample size")  + ylab(bquote(PI[Inf]))+scale_colour_manual(values =
c("dodgerblue4","skyblue2","coral2","coral4"))+geom_point(size = 1)
+geom_errorbar(aes(ymin = DF_avg-DF_var, ymax = DF_avg+DF_var))
+facet_grid(vars(Case), vars(DFtype))+ theme(strip.text.x = element_text(size
= 12, face = "bold"),strip.text.y = element_text(size = 12, face = "bold" ))
+theme(axis.title =  element_text(size = 15))+theme(axis.text =
element_text(size = 10))+theme(aspect.ratio=1)+ theme(panel.grid.major =
element_blank(),

panel.grid.minor = element_blank())+theme(legend.position= "none")
+theme(axis.text.x=element_text(color=c("black","transparent","transparent","black")))
+theme(axis.text.y=element_text(color=c("black","transparent","transparent","transparen

################Figure 8###########################################
ggplot(combined2,aes(j,Drilledsp_avg,color=factor_Eventype)) +  xlab("Sample size")
+ylab(bquote(S[Prey.Inf]))+scale_colour_manual(values =
c("dodgerblue4","skyblue2","coral2","coral4"))+geom_point(size = 1)
+geom_errorbar(aes(ymin = Drilledsp_avg-Drilledsp_var, ymax =
Drilledsp_avg+Drilledsp_var))+facet_grid(vars(Case),vars(DFtype))+ theme(strip.text.x
= element_text(size = 12, face = "bold"),strip.text.y = element_text(size = 12, face =
"bold" ))+theme(axis.title =  element_text(size = 15))+theme(axis.text =
element_text(size = 10))+theme(aspect.ratio=1)+ theme(panel.grid.major =
element_blank(),panel.grid.minor = element_blank())+theme(legend.position= "none")
+theme(axis.text.x=element_text(color=c("black","transparent","transparent","black")))
+theme(axis.text.y=element_text(color=c("black","transparent","transparent","black")))



#############Figure 12#################################################
```

```
ggplot(combined2,aes(Drilledsp_avg,DF_avg,color=factor_Eventype)) +
xlab(bquote(S[Prey.Inf])) + ylab(bquote(PI[Inf]))
+scale_colour_manual(values =
c("dodgerblue4","skyblue2","coral2","coral4"))+geom_point(size = 1)+
geom_errorbar(aes(ymin = DF_avg-DF_var, ymax = DF_avg+DF_var))
+geom_errorbarh(aes(xmin = Drilledsp_avg-Drilledsp_var, xmax =
Drilledsp_avg+Drilledsp_var))+facet_grid(vars(Case),vars(DFtype))
+theme(strip.text.x = element_text(size = 12, face = "bold"),strip.text.y =
element_text(size = 12, face = "bold" ))+theme(axis.title =
element_text(size = 15))+theme(axis.text =  element_text(size = 10))
+theme(aspect.ratio=1)+ theme(panel.grid.major = element_blank(),

panel.grid.minor = element_blank())+theme(legend.position= "none")
+theme(axis.text.x=element_text(color=c("black","transparent","transparent","black")))
+theme(axis.text.y=element_text(color=c("black","transparent","transparent","transparen

####making subset data for case 1###########
combinedcase1=subset(combined2,combined2$Case=="Case 1")
j=combinedcase1$j
DFtype=combinedcase1$DFtype
Eventype=combinedcase1$Eventype
Case=combinedcase1$Case
factor_Eventype=factor(Eventype)
DFtype=factor(DFtype)
Drilledsp_avg=combinedcase1$uniquedrilledavg.j.
Drilledsp_var=combinedcase1$uniquedrilledvar.j.
DF_avg=combinedcase1$DFavg.j.
DF_var=combinedcase1$DFvar.j.
DE_avg=combinedcase1$Devavg.j.
DE_var=combinedcase1$Devvar.j.
devpreyavg=combinedcase1$devpreyavg.j.
devpreyvar=combinedcase1$devpreyvar.j.
combinedcase1$Eventype=round(combinedcase1[,10],1)
ggplot(combinedcase1,aes(j,DE_avg,gcoroup=factor_Eventype,color=factor_Eventype))
+  xlab("Sample size") + ylab("Deviation")+geom_hline(yintercept=0,
color="red",size=1)+scale_colour_manual(values =
c("dodgerblue4","skyblue2","salmon2","orangered4"))+geom_point(size = 1)
+geom_errorbar(aes(ymin = DE_avg-DE_var, ymax = DE_avg+DE_var))
+facet_grid(vars(Eventype), vars(DFtype))+ theme(strip.text.x = element_text(size
= 12, face = "bold"),strip.text.y = element_text(size = 12, face = "bold" ))
+theme(axis.title =  element_text(size = 15))+theme(axis.text =
element_text(size = 10))+theme(aspect.ratio=1)+theme(
  panel.grid.major = element_blank(),
  panel.grid.minor = element_blank())+theme(legend.position= "none")

#######figure 5#########################################
ggplot(combinedcase1,aes(j,DE_avg,gcoroup=factor_Eventype,color=factor_Eventype))
+coord_cartesian(xlim = c(100,3000),ylim =c(-0.04,0.04))+  xlab("Sample size") +
ylab(bquote(Dev[PI]))+geom_hline(yintercept=0, color="red",size=0.5)
+theme(panel.spacing.x = unit(4, "mm"))
+theme(axis.text.x=element_text(color=c("black","transparent","transparent","black")))
+scale_colour_manual(values = c("grey","grey","grey","grey"))+geom_point(size = 1)
+geom_errorbar(aes(ymin = DE_avg-DE_var, ymax = DE_avg+DE_var))
+facet_grid(vars(Eventype),vars(DFtype))+ theme(strip.text.x = element_text(size = 12,
face = "bold"),strip.text.y = element_text(size = 12, face = "bold" ))
+theme(axis.title =  element_text(size = 15))+theme(axis.text =  element_text(size =
10))+theme(aspect.ratio=1)+theme(
  panel.grid.major = element_blank(),
```

```
  panel.grid.minor = element_blank())+theme(panel.background = element_blank())
+theme(legend.position= "none")+
theme(axis.text.y=element_text(color=c("black","transparent","black","transparent","bla

#######figure 9###########################################
ggplot(combinedcase1,aes(j,devpreyavg,gcoroup=factor_Eventype,color=factor_Eventype))
+coord_cartesian(xlim = c(100,3000),ylim=c(0,30))+  xlab("Sample size") +
ylab(bquote(Dev[S]))+
theme(axis.text.x=element_text(color=c("black","transparent","transparent","black")))
+geom_hline(yintercept=0, color="red",size=0.5)+scale_colour_manual(values =
c("grey","grey","grey","grey"))+geom_point(size = 1)+geom_errorbar(aes(ymin =
devpreyavg-devpreyvar, ymax = devpreyavg+devpreyvar))+facet_grid(vars(Eventype),
vars(DFtype))+ theme(strip.text.x = element_text(size = 12, face =
"bold"),strip.text.y = element_text(size = 12, face = "bold" ))+theme(axis.title =
element_text(size = 15))+theme(axis.text =  element_text(size = 10))
+theme(aspect.ratio=1)+theme(

  panel.grid.major = element_blank(),
  panel.grid.minor = element_blank())+theme(panel.background = element_blank())
+theme(legend.position= "none")+theme(panel.spacing.x = unit(4, "mm"))+
theme(axis.text.y=element_text(color=c("black","transparent","transparent","black")))


####making subset data for case 2###############
combinedcase2=subset(combined2,combined2$Case=="Case 2")
j=combinedcase2$j
DFtype=combinedcase2$DFtype
Eventype=combinedcase2$Eventype
Case=combinedcase2$Case
factor_Eventype=factor(Eventype)
DFtype=factor(DFtype)
Drilledsp_avg=combinedcase2$uniquedrilledavg.j.
Drilledsp_var=combinedcase2$uniquedrilledvar.j.
DF_avg=combinedcase2$DFavg.j.
DF_var=combinedcase2$DFvar.j.
DE_avg=combinedcase2$Devavg.j.
DE_var=combinedcase2$Devvar.j.
devpreyavg=combinedcase2$devpreyavg.j.
devpreyvar=combinedcase2$devpreyvar.j.
combinedcase2$Eventype=round(combinedcase2[,10],1)

#######figure 6###########################################
ggplot(combinedcase2,aes(j,DE_avg,gcoroup=factor_Eventype,color=factor_Eventype))
+coord_cartesian(xlim = c(100,3000),ylim =c(-0.04,0.04))+  xlab("Sample size") +
ylab(bquote(Dev[PI]))+geom_hline(yintercept=0, color="red",size=0.5)
+theme(panel.spacing.x = unit(4, "mm"))
+theme(axis.text.x=element_text(color=c("black","transparent","transparent","black")))
+scale_colour_manual(values = c("grey","grey","grey","grey"))+geom_point(size = 1)
+geom_errorbar(aes(ymin = DE_avg-DE_var, ymax = DE_avg+DE_var))
+facet_grid(vars(Eventype),vars(DFtype))+ theme(strip.text.x = element_text(size = 12,
face = "bold"),strip.text.y = element_text(size = 12, face = "bold" ))
+theme(axis.title =  element_text(size = 15))+theme(axis.text =  element_text(size =
10))+theme(aspect.ratio=1)+theme(
  panel.grid.major = element_blank(),
  panel.grid.minor = element_blank())+theme(panel.background = element_blank())
+theme(legend.position= "none")
+theme(axis.text.y=element_text(color=c("black","transparent","black","transparent","bl
```

```
ggplot(combinedcase2,aes(j,devpreyavg,gcoroup=factor_Eventype,color=factor_Eventype))
+coord_cartesian(xlim = c(100,3000),ylim=c(-0.4, 0.4))+xlab("Sample size") +
theme(axis.text.x=element_text(color=c("black","transparent","transparent","black")))
+ylab(bquote(Dev[S]))+geom_hline(yintercept=0, color="red",size=0.5)
+scale_colour_manual(values = c("grey","grey","grey","grey"))+geom_point(size = 1)
+geom_errorbar(aes(ymin = devpreyavg-devpreyvar, ymax = devpreyavg+devpreyvar))
+facet_grid(vars(Eventype), vars(DFtype))+ theme(strip.text.x = element_text(size =
12, face = "bold"),strip.text.y = element_text(size = 12, face = "bold" ))
+theme(axis.title =  element_text(size = 15))+theme(axis.text =  element_text(size =
10))+theme(aspect.ratio=1)+theme(


  panel.grid.major = element_blank(),
  panel.grid.minor = element_blank())+theme(panel.background = element_blank())+
theme(axis.text.y=element_text(color=c("black","transparent","black","transparent","bla
+theme(panel.spacing.x = unit(4, "mm"))


###################figure 10###########################################
ggplot(combinedcase2,aes(j,devpreyavg,gcoroup=factor_Eventype,color=factor_Eventype))
+coord_cartesian(xlim = c(100,3000),ylim=c(-0.4, 0.4))+xlab("Sample size") +
ylab(bquote(Dev[S]))+
theme(axis.text.x=element_text(color=c("black","transparent","transparent","black")))
+geom_hline(yintercept=0, color="red",size=0.5)+scale_colour_manual(values =
c("grey","grey","grey","grey"))+geom_point(size = 1)+geom_errorbar(aes(ymin =
devpreyavg-devpreyvar, ymax = devpreyavg+devpreyvar))+facet_grid(vars(Eventype),
vars(DFtype))+ theme(strip.text.x = element_text(size = 12, face =
"bold"),strip.text.y = element_text(size = 12, face = "bold" ))+theme(axis.title =
element_text(size = 15))+theme(axis.text =  element_text(size = 10))
+theme(aspect.ratio=1)+theme(

  panel.grid.major = element_blank(),
  panel.grid.minor = element_blank())+theme(panel.background = element_blank())
+theme(legend.position= "none")+theme(panel.spacing.x = unit(4, "mm"))+
theme(axis.text.y=element_text(color=c("black","transparent","black","transparent","bla


#####creating subset for case
3##########################################################################
combinedcase3=subset(combined2,combined2$Case=="Case 3")
j=combinedcase3$j
DFtype=combinedcase3$DFtype
Eventype=combinedcase3$Eventype
Case=combinedcase3$Case
factor_Eventype=factor(Eventype)
DFtype=factor(DFtype)
Drilledsp_avg=combinedcase3$uniquedrilledavg.j.
Drilledsp_var=combinedcase3$uniquedrilledvar.j.
DF_avg=combinedcase3$DFavg.j.
DF_var=combinedcase3$DFvar.j.
DE_avg=combinedcase3$Devavg.j.
DE_var=combinedcase3$Devvar.j.
devpreyavg=combinedcase3$devpreyavg.j.
devpreyvar=combinedcase3$devpreyvar.j.

combinedcase3$Eventype=round(combinedcase3[,10],1)

###############################Figure 7################################
```

```
ggplot(combinedcase3,aes(j,DE_avg,gcoroup=factor_Eventype,color=factor_Eventype))
+coord_cartesian(xlim = c(100,3000),ylim =c(-0.04,0.04))+  xlab("Sample size") +
ylab(bquote(Dev[PI]))+geom_hline(yintercept=0, color="red",size=0.5)
+theme(panel.spacing.x = unit(4, "mm"))
+theme(axis.text.x=element_text(color=c("black","transparent","transparent","black")))
+scale_colour_manual(values = c("grey","grey","grey","grey"))+geom_point(size = 1)
+geom_errorbar(aes(ymin = DE_avg-DE_var, ymax = DE_avg+DE_var))
+facet_grid(vars(Eventype),vars(DFtype))+ theme(strip.text.x = element_text(size = 12,
face = "bold"),strip.text.y = element_text(size = 12, face = "bold" ))
+theme(axis.title =  element_text(size = 15))+theme(axis.text =  element_text(size =
10))+theme(aspect.ratio=1)+theme(
  panel.grid.major = element_blank(),
  panel.grid.minor = element_blank())+theme(panel.background = element_blank())
+theme(panel.spacing.x = unit(4, "mm"))+theme(legend.position= "none")
+theme(axis.text.y=element_text(color=c("black","transparent","black","transparent","bl


########figure 11###########################################
ggplot(combinedcase3,aes(j,devpreyavg,gcoroup=factor_Eventype,color=factor_Eventype))
+coord_cartesian(xlim = c(100,3000),ylim=c(0,30))
+theme(axis.text.x=element_text(color=c("black","transparent","transparent","black")))
+ xlab("Sample size") + ylab(bquote(Dev[S]))+ geom_hline(yintercept=0,
color="red",size=0.5)+scale_colour_manual(values = c("grey","grey","grey","grey"))
+geom_point(size = 1)+geom_errorbar(aes(ymin = devpreyavg-devpreyvar, ymax =
devpreyavg+devpreyvar))+facet_grid(vars(Eventype), vars(DFtype))+ theme(strip.text.x =
element_text(size = 12, face = "bold"),strip.text.y = element_text(size = 12, face =
"bold" ))+theme(axis.title =  element_text(size = 15))+theme(axis.text =
element_text(size = 10))+theme(aspect.ratio=1)+theme(

  panel.grid.major = element_blank(),
  panel.grid.minor = element_blank())+theme(panel.background = element_blank())
+theme(panel.spacing.x = unit(4, "mm"))+theme(legend.position= "none")+
theme(axis.text.y=element_text(color=c("black","transparent","transparent","black")))



########################Florida
locations##########################################################################
require(vegan)
###########Punta Gorda#####################################

neuseriver=read.csv("Punta Gorda.csv",header=T)
Individual=rep(1,2418)
neuseriver2=data.frame(Individual,neuseriver)
wideform=as.data.frame.matrix(table(neuseriver2$Individual,neuseriver2$Species))
evenness=(diversity(wideform)/log(specnumber(wideform)))

###evenness=0.467#######
######Drilling frequency###########################
Eventype=evenness
Location="Punta Gorda"
a=cbind(Eventype,Location)


randomnumber=vector(mode = "integer",length=2418)
raredata=array(0,dim = c(2418,2))
num_species1=vector(mode="integer",length=length(unique(neuseriver2$Species)))
```

```
num_drilled=vector(mode = "integer", length =
length(unique(neuseriver2$Species)))
DF=vector(mode = "integer", length = length(unique(neuseriver2$Species)))
neuseriver_subset=subset(neuseriver2,select = c(Species,Drilled))
subsamps <- list()
subsamps1=list()
subsamps2=list()
for (q in 1:1000)
{
  randomnumber=sample.int(2418,2418, replace = FALSE)
  raredata=neuseriver_subset[randomnumber[1:2418],]
  for(i in 1:24)
  {
    j=(i*100)
    {
      if((2418-j)==18)
      {
        c=j+18
        uniquesp=unique(raredata$Species[1:c])
        num_species1[c]=length(uniquesp)
        uniquedrilled=unique(raredata$Species[which(raredata$Drilled[1:c]==1)])
        DF[c]=(sum(raredata$Drilled[1:c]))/c
        num_drilled[c]=length(uniquedrilled)
        oute=data.frame(c,num_species1[c],num_drilled[c],DF[c])
        subsamps2[[c]] <-oute
      }
      else
      {
        uniquesp=unique(raredata$Species[1:j])
        num_species1[j]=length(uniquesp)
        uniquedrilled=unique(raredata$Species[which(raredata$Drilled[1:j]==1)])
        DF[j]=(sum(raredata$Drilled[1:j]))/j
        num_drilled[j]=length(uniquedrilled)
        out=data.frame(j,num_species1[j],num_drilled[j],DF[j])
        subsamps1[[i]] <-out
      }
    }
  }
  subsamples1 <- do.call("rbind",subsamps1)
  subsamples2<-do.call("rbind",subsamps2)
  colnames(subsamples2)=colnames(subsamples1)
  subsamplesfinal=rbind(subsamples1,subsamples2)
  out2=data.frame(q,subsamplesfinal)
  subsamps[[q]]=out2
  subsamples <- do.call("rbind",subsamps)
}
subsamples_punta=as.data.frame(subsamples)
subsamples_punta[is.na(subsamples_punta)] = 0



#######################################################################################
############For calculating average and
variance#####################################################
uniquedrilledavg=vector(mode = "integer", length =
length(unique(neuseriver2$Species)))
uniquedrilledvar=vector(mode = "integer", length =
length(unique(neuseriver2$Species)))
```

```
DFavg=vector(mode = "integer", length =
length(unique(neuseriver2$Species)))
DFvar=vector(mode = "integer", length =
length(unique(neuseriver2$Species)))
subsamps1=list()
subsamps2=list()

for(i in 1:24)
{
  j=(i*100)
  {
    if((2418-j)==18)
    {
      c=j+18
      uniquedrilledavg[c]=mean(subsamples_punta$num_drilled.j.
[which(subsamples_punta$j==c)])
      uniquedrilledvar[c]=sd(subsamples_punta$num_drilled.j.
[which(subsamples_punta$j==c)])
      DFavg[c]=mean(subsamples_punta$DF.j.[which(subsamples_punta$j==c)])
      DFvar[c]=sd(subsamples_punta$DF.j.[which(subsamples_punta$j==c)])
      oute=data.frame(c,uniquedrilledavg[c],uniquedrilledvar[c],DFavg[c],DFvar[c],a)
      subsamps2[[i]] <-oute
    }
    else
    {
      uniquedrilledavg[j]=mean(subsamples_punta$num_drilled.j.
[which(subsamples_punta$j==j)])
      uniquedrilledvar[j]=sd(subsamples_punta$num_drilled.j.
[which(subsamples_punta$j==j)])
      DFavg[j]=mean(subsamples_punta$DF.j.[which(subsamples_punta$j==j)])
      DFvar[j]=sd(subsamples_punta$DF.j.[which(subsamples_punta$j==j)])
      out=data.frame(j,uniquedrilledavg[j],uniquedrilledvar[j],DFavg[j],DFvar[j],a)
      subsamps1[[i]] <-out
    }
  }
}
subsamples3<- do.call("rbind",subsamps1)
subsamples4=do.call("rbind",subsamps2)
colnames(subsamples4)=colnames(subsamples3)
subsamplesfinal1=rbind(subsamples3,subsamples4)

Drilledsp_avg=subsamplesfinal1$uniquedrilledavg.j.
Drilledsp_var=subsamplesfinal1$uniquedrilledvar.j.
DF_avg=subsamplesfinal1$DFavg.j.
DF_var=subsamplesfinal1$DFvar.j.



###########Repair frequency###################
Eventype=evenness
Location="Punta Gorda"
a=cbind(Eventype,Location)

randomnumber=vector(mode = "integer",length=2418)
raredata=array(0,dim = c(2418,2))
num_species1=vector(mode="integer",length=length(unique(neuseriver2$Species)))
num_repaired=vector(mode = "integer", length =
length(unique(neuseriver2$Species)))
```

```
RF=vector(mode = "integer", length = length(unique(neuseriver2$Species)))
neuseriver_subset_2=subset(neuseriver2,select = c(Species,Repaired))
subsamps <- list()
subsamps1=list()
subsamps2=list()

for (q in 1:1000)
{
  randomnumber=sample.int(2418,2418, replace = FALSE)
  raredata=neuseriver_subset_2[randomnumber[1:2418],]
  for(i in 1:24)
  {
    j=(i*100)
    {
      if((2418-j)==18)
      {
        c=j+18
        uniquesp=unique(raredata$Species[1:c])
        num_species1[c]=length(uniquesp)
        uniquerepaired=unique(raredata$Species[which(raredata$Repaired[1:c]==1)])
        RF[c]=(sum(raredata$Repaired[1:c]))/c
        num_repaired[c]=length(uniquerepaired)
        oute=data.frame(c,num_species1[c],num_repaired[c],RF[c])
        subsamps2[[c]] <-oute
      }
      else
      {
        uniquesp=unique(raredata$Species[1:j])
        num_species1[j]=length(uniquesp)
        uniquerepaired=unique(raredata$Species[which(raredata$Repaired[1:j]==1)])
        num_repaired[j]=length(uniquerepaired)
        RF[j]=(sum(raredata$Repaired[1:j]))/j
        out=data.frame(j,num_species1[j],num_repaired[j],RF[j])
        subsamps1[[i]] <-out
      }
    }
  }
  subsamples1 <- do.call("rbind",subsamps1)
  subsamples2<-do.call("rbind",subsamps2)
  colnames(subsamples2)=colnames(subsamples1)
  subsamplesfinal3=rbind(subsamples1,subsamples2)
  out2=data.frame(q,subsamplesfinal3)
  subsamps[[q]]=out2
  subsamples <- do.call("rbind",subsamps)
}
subsamples_punta_repair=as.data.frame(subsamples)
subsamples_punta_repair[is.na(subsamples_punta_repair)] = 0



#####################################################################################
############For calculating average and
variance#####################################################
uniquerepairedavg=vector(mode = "integer", length =
length(unique(neuseriver2$Species)))
uniquerepairedvar=vector(mode = "integer", length =
length(unique(neuseriver2$Species)))
```

```
RFavg=vector(mode = "integer", length =
length(unique(neuseriver2$Species)))
RFvar=vector(mode = "integer", length =
length(unique(neuseriver2$Species)))
subsamps <- list()
subsamps1=list()
subsamps2=list()


for(i in 1:24)
{
  j=(i*100)
  {
    if((2418-j)==18)
    {
      c=j+18
      uniquerepairedavg[c]=mean(subsamples_punta_repair$num_repaired.j.
[which(subsamples_punta_repair$j==c)])
      uniquerepairedvar[c]=sd(subsamples_punta_repair$num_repaired.j.
[which(subsamples_punta_repair$j==c)])
      RFavg[c]=mean(subsamples_punta_repair$RF.j.
[which(subsamples_punta_repair$j==c)])
      RFvar[c]=sd(subsamples_punta_repair$RF.j.
[which(subsamples_punta_repair$j==c)])
      oute=data.frame(c,uniquerepairedavg[c],uniquerepairedvar[c],RFavg[c],RFvar[c],a)
      subsamps2[[i]] <-oute
    }
    else
    {
      uniquerepairedavg[j]=mean(subsamples_punta_repair$num_repaired.j.
[which(subsamples_punta_repair$j==j)])
      uniquerepairedvar[j]=sd(subsamples_punta_repair$num_repaired.j.
[which(subsamples_punta_repair$j==j)])
      RFavg[j]=mean(subsamples_punta_repair$RF.j.
[which(subsamples_punta_repair$j==j)])
      RFvar[j]=sd(subsamples_punta_repair$RF.j.
[which(subsamples_punta_repair$j==j)])
      out=data.frame(j,uniquerepairedavg[j],uniquerepairedvar[j],RFavg[j],RFvar[j],a)
      subsamps1[[i]] <-out
    }
  }
}
subsamples3<- do.call("rbind",subsamps1)
subsamples4=do.call("rbind",subsamps2)
colnames(subsamples4)=colnames(subsamples3)
subsamplesfinal4=rbind(subsamples3,subsamples4)

Repairedsp_avg=subsamplesfinal4$uniquerepairedavg.j.
Repairedsp_var=subsamplesfinal4$uniquerepairedvar.j.
RF_avg=subsamplesfinal4$RFavg.j.
RF_var=subsamplesfinal4$RFvar.j.

###subsamples3 for df###
####subsamples4 for rf#######

###################Miami canal######################################

neuseriver=read.csv("Miami canal.csv",header=T)
```

```
Individual=rep(1,4794)
neuseriver2=data.frame(Individual,neuseriver)
wideform=as.data.frame.matrix(table(neuseriver2$Individual,neuseriver2$Species))
evenness=(diversity(wideform)/log(specnumber(wideform)))

###evenness=0.315#######
######Drilling frequency############################
Eventype=evenness
Location="Miami canal"
a=cbind(Eventype,Location)


randomnumber=vector(mode = "integer",length=4794)
raredata=array(0,dim = c(4794,2))
num_species1=vector(mode="integer",length=length(unique(neuseriver2$Species)))
num_drilled=vector(mode = "integer", length =
length(unique(neuseriver2$Species)))
DF=vector(mode = "integer", length = length(unique(neuseriver2$Species)))
neuseriver_subset=subset(neuseriver2,select = c(Species,Drilled))
subsamps <- list()
subsamps1=list()
subsamps2=list()
for (q in 1:1000)
{
  randomnumber=sample.int(4794,4794, replace = FALSE)
  raredata=neuseriver_subset[randomnumber[1:4794],]
  for(i in 1:47)
  {
    j=(i*100)
    {
      if((4794-j)==94)
      {
        c=j+94
        uniquesp=unique(raredata$Species[1:c])
        num_species1[c]=length(uniquesp)
        uniquedrilled=unique(raredata$Species[which(raredata$Drilled[1:c]==1)])
        DF[c]=(sum(raredata$Drilled[1:c]))/c
        num_drilled[c]=length(uniquedrilled)
        oute=data.frame(c,num_species1[c],num_drilled[c],DF[c])
        subsamps2[[c]] <-oute
      }
      else
      {
        uniquesp=unique(raredata$Species[1:j])
        num_species1[j]=length(uniquesp)
        uniquedrilled=unique(raredata$Species[which(raredata$Drilled[1:j]==1)])
        DF[j]=(sum(raredata$Drilled[1:j]))/j
        num_drilled[j]=length(uniquedrilled)
        out=data.frame(j,num_species1[j],num_drilled[j],DF[j])
        subsamps1[[i]] <-out
      }
    }
  }
  subsamples1 <- do.call("rbind",subsamps1)
  subsamples2<-do.call("rbind",subsamps2)
  colnames(subsamples2)=colnames(subsamples1)
  subsamplesfinal5=rbind(subsamples1,subsamples2)
  out2=data.frame(q,subsamplesfinal5)
```

```
  subsamps[[q]]=out2
  subsamples <- do.call("rbind",subsamps)
}
subsamples_miami=as.data.frame(subsamples)
subsamples_miami[is.na(subsamples_miami)] = 0




#####################################################################################
############For calculating average and
variance########################################################
uniquedrilledavg=vector(mode = "integer", length =
length(unique(neuseriver2$Species)))
uniquedrilledvar=vector(mode = "integer", length =
length(unique(neuseriver2$Species)))
DFavg=vector(mode = "integer", length =
length(unique(neuseriver2$Species)))
DFvar=vector(mode = "integer", length =
length(unique(neuseriver2$Species)))
subsamps1=list()
subsamps2=list()


for(i in 1:47)
{
  j=(i*100)
  {
    if((4794-j)==94)
    {
      c=j+94
      uniquedrilledavg[c]=mean(subsamples_miami$num_drilled.j.
[which(subsamples_miami$j==c)])
      uniquedrilledvar[c]=sd(subsamples_miami$num_drilled.j.
[which(subsamples_miami$j==c)])
      DFavg[c]=mean(subsamples_miami$DF.j.[which(subsamples_miami$j==c)])
      DFvar[c]=sd(subsamples_miami$DF.j.[which(subsamples_miami$j==c)])
      oute=data.frame(c,uniquedrilledavg[c],uniquedrilledvar[c],DFavg[c],DFvar[c],a)
      subsamps2[[i]] <-oute
    }
    else
    {
      uniquedrilledavg[j]=mean(subsamples_miami$num_drilled.j.
[which(subsamples_miami$j==j)])
      uniquedrilledvar[j]=sd(subsamples_miami$num_drilled.j.
[which(subsamples_miami$j==j)])
      DFavg[j]=mean(subsamples_miami$DF.j.[which(subsamples_miami$j==j)])
      DFvar[j]=sd(subsamples_miami$DF.j.[which(subsamples_miami$j==j)])
      out=data.frame(j,uniquedrilledavg[j],uniquedrilledvar[j],DFavg[j],DFvar[j],a)
      subsamps1[[i]] <-out
    }
  }
}
subsamples3<- do.call("rbind",subsamps1)
subsamples4=do.call("rbind",subsamps2)
colnames(subsamples4)=colnames(subsamples3)
subsamplesfinal6=rbind(subsamples3,subsamples4)
Drilledsp_avg=subsamplesfinal6$uniquedrilledavg.j.
```

```
Drilledsp_var=subsamplesfinal6$uniquedrilledvar.j.
DF_avg=subsamplesfinal6$DFavg.j.
DF_var=subsamplesfinal6$DFvar.j.


###########Repair frequency###################
Eventype=evenness
Location="Miami canal"
a=cbind(Eventype,Location)

randomnumber=vector(mode = "integer",length=4794)
raredata=array(0,dim = c(4794,2))
num_species1=vector(mode="integer",length=length(unique(neuseriver2$Species)))
num_repaired=vector(mode = "integer", length =
length(unique(neuseriver2$Species)))
RF=vector(mode = "integer", length = length(unique(neuseriver2$Species)))
neuseriver_subset_2=subset(neuseriver2,select = c(Species,Repaired))
subsamps <- list()
subsamps1=list()
subsamps2=list()
for (q in 1:1000)
{
  randomnumber=sample.int(4794,4794, replace = FALSE)
  raredata=neuseriver_subset_2[randomnumber[1:4794],]
  for(i in 1:47)
  {
    j=(i*100)
    {
      if((4794-j)==94)
      {
        c=j+94
        uniquesp=unique(raredata$Species[1:c])
        num_species1[c]=length(uniquesp)
        uniquedrilled=unique(raredata$Species[which(raredata$Repaired[1:c]==1)])
        RF[c]=(sum(raredata$Repaired[1:c]))/c
        num_repaired[c]=length(uniquerepaired)
        oute=data.frame(c,num_species1[c],num_repaired[c],RF[c])
        subsamps2[[c]] <-oute
      }
      else
      {
        uniquesp=unique(raredata$Species[1:j])
        num_species1[j]=length(uniquesp)
        uniquerepaired=unique(raredata$Species[which(raredata$Repaired[1:j]==1)])
        num_repaired[j]=length(uniquerepaired)
        RF[j]=(sum(raredata$Repaired[1:j]))/j
        out=data.frame(j,num_species1[j],num_repaired[j],RF[j])
        subsamps1[[i]] <-out
      }
    }
  }
  subsamples1 <- do.call("rbind",subsamps1)
  subsamples2<-do.call("rbind",subsamps2)
  colnames(subsamples2)=colnames(subsamples1)
  subsamplesfinal7=rbind(subsamples1,subsamples2)
  out2=data.frame(q,subsamplesfinal7)
  subsamps[[q]]=out2
  subsamples <- do.call("rbind",subsamps)
```

```
}
subsamples_miami_repair=as.data.frame(subsamples)
subsamples_miami_repair[is.na(subsamples_miami_repair)] = 0



########################################################################################
############For calculating average and
variance##########################################################
uniquerepairedavg=vector(mode = "integer", length =
length(unique(neuseriver2$Species)))
uniquerepairedvar=vector(mode = "integer", length =
length(unique(neuseriver2$Species)))
RFavg=vector(mode = "integer", length =
length(unique(neuseriver2$Species)))
RFvar=vector(mode = "integer", length =
length(unique(neuseriver2$Species)))
subsamps1=list()
subsamps2=list()

for(i in 1:47)
{
  j=(i*100)
  {
    if((4794-j)==94)
    {
      c=j+94
      uniquerepairedavg[c]=mean(subsamples_miami_repair$num_repaired.j.
[which(subsamples_miami_repair$j==c)])
      uniquerepairedvar[c]=sd(subsamples_miami_repair$num_repaired.j.
[which(subsamples_miami_repair$j==c)])
      RFavg[c]=mean(subsamples_miami_repair$RF.j.
[which(subsamples_miami_repair$j==c)])
      RFvar[c]=sd(subsamples_miami_repair$RF.j.
[which(subsamples_miami_repair$j==c)])
      oute=data.frame(c,uniquerepairedavg[c],uniquerepairedvar[c],RFavg[c],RFvar[c],a)
      subsamps2[[i]] <-oute
    }
    else
    {
      uniquerepairedavg[j]=mean(subsamples_miami_repair$num_repaired.j.
[which(subsamples_miami_repair$j==j)])
      uniquerepairedvar[j]=sd(subsamples_miami_repair$num_repaired.j.
[which(subsamples_miami_repair$j==j)])
      RFavg[j]=mean(subsamples_miami_repair$RF.j.
[which(subsamples_miami_repair$j==j)])
      RFvar[j]=sd(subsamples_miami_repair$RF.j.
[which(subsamples_miami_repair$j==j)])
      out=data.frame(j,uniquerepairedavg[j],uniquerepairedvar[j],RFavg[j],RFvar[j],a)
      subsamps1[[i]] <-out
    }
  }
}
subsamples3<- do.call("rbind",subsamps1)
subsamples4=do.call("rbind",subsamps2)
colnames(subsamples4)=colnames(subsamples3)
subsamplesfinal8=rbind(subsamples3,subsamples4)
```

```
Repairedsp_avg=subsamplesfinal8$uniquerepairedavg.j.
Repairedsp_var=subsamplesfinal8$uniquerepairedvar.j.
RF_avg=subsamplesfinal8$RFavg.j.
RF_var=subsamplesfinal8$RFvar.j.

###subsamples5 for df###
####subsamples6 for rf#######

##########Mc queen##############################
neuseriver=read.csv("Mcqueen.csv",header=T)
Individual=rep(1,659)
neuseriver2=data.frame(Individual,neuseriver)
wideform=as.data.frame.matrix(table(neuseriver2$Individual,neuseriver2$Species))
evenness=(diversity(wideform)/log(specnumber(wideform)))

###evenness=0.315#######
######Drilling frequency###########################
Eventype=evenness
Location="Mc Queens pit"
a=cbind(Eventype,Location)


randomnumber=vector(mode = "integer",length=659)
raredata=array(0,dim = c(659,2))
num_species1=vector(mode="integer",length=length(unique(neuseriver2$Species)))
num_drilled=vector(mode = "integer", length =
length(unique(neuseriver2$Species)))
DF=vector(mode = "integer", length = length(unique(neuseriver2$Species)))
neuseriver_subset=subset(neuseriver2,select = c(Species,Drilled))
subsamps <- list()
subsamps1=list()
subsamps2=list()
for (q in 1:1000)
{
  randomnumber=sample.int(659,659, replace = FALSE)
  raredata=neuseriver_subset[randomnumber[1:659],]
  for(i in 1:6)
  {
    j=(i*100)
    {
      if((659-j)==59)
      {
        c=j+59
        uniquesp=unique(raredata$Species[1:c])
        num_species1[c]=length(uniquesp)
        uniquedrilled=unique(raredata$Species[which(raredata$Drilled[1:c]==1)])
        DF[c]=(sum(raredata$Drilled[1:c]))/c
        num_drilled[c]=length(uniquedrilled)
        oute=data.frame(c,num_species1[c],num_drilled[c],DF[c])
        subsamps2[[c]] <-oute
      }
      else
      {
        uniquesp=unique(raredata$Species[1:j])
        num_species1[j]=length(uniquesp)
        uniquedrilled=unique(raredata$Species[which(raredata$Drilled[1:j]==1)])
        DF[j]=(sum(raredata$Drilled[1:j]))/j
```

```
        num_drilled[j]=length(uniquedrilled)
        out=data.frame(j,num_species1[j],num_drilled[j],DF[j])
        subsamps1[[i]] <-out
      }
    }
  }
  subsamples1 <- do.call("rbind",subsamps1)
  subsamples2<-do.call("rbind",subsamps2)
  colnames(subsamples2)=colnames(subsamples1)
  subsamplesfinal9=rbind(subsamples1,subsamples2)
  out2=data.frame(q,subsamplesfinal9)
  subsamps[[q]]=out2
  subsamples <- do.call("rbind",subsamps)
}
subsamples_mcqueen=as.data.frame(subsamples)
subsamples_mcqueen[is.na(subsamples_mcqueen)] = 0


#################################################################################
############For calculating average and
variance#########################################################
uniquedrilledavg=vector(mode = "integer", length =
length(unique(neuseriver2$Species)))
uniquedrilledvar=vector(mode = "integer", length =
length(unique(neuseriver2$Species)))
DFavg=vector(mode = "integer", length =
length(unique(neuseriver2$Species)))
DFvar=vector(mode = "integer", length =
length(unique(neuseriver2$Species)))
subsamps1=list()
subsamps2=list()
for(i in 1:6)
{
  j=(i*100)
  {
    if((659-j)==59)
    {
      c=j+59
      uniquedrilledavg[c]=mean(subsamples_mcqueen$num_drilled.j.
[which(subsamples_mcqueen$j==c)])
      uniquedrilledvar[c]=sd(subsamples_mcqueen$num_drilled.j.
[which(subsamples_mcqueen$j==c)])
      DFavg[c]=mean(subsamples_mcqueen$DF.j.
[which(subsamples_mcqueen$j==c)])
      DFvar[c]=sd(subsamples_mcqueen$DF.j.[which(subsamples_mcqueen$j==c)])
      oute=data.frame(c,uniquedrilledavg[c],uniquedrilledvar[c],DFavg[c],DFvar[c],a)
      subsamps2[[i]] <-oute
    }
    else
    {
      uniquedrilledavg[j]=mean(subsamples_mcqueen$num_drilled.j.
[which(subsamples_mcqueen$j==j)])
      uniquedrilledvar[j]=sd(subsamples_mcqueen$num_drilled.j.
[which(subsamples_mcqueen$j==j)])
      DFavg[j]=mean(subsamples_mcqueen$DF.j.
[which(subsamples_mcqueen$j==j)])
      DFvar[j]=sd(subsamples_mcqueen$DF.j.[which(subsamples_mcqueen$j==j)])
      out=data.frame(j,uniquedrilledavg[j],uniquedrilledvar[j],DFavg[j],DFvar[j],a)
```

```
      subsamps1[[i]] <-out
    }
  }
}
subsamples3<- do.call("rbind",subsamps1)
subsamples4=do.call("rbind",subsamps2)
colnames(subsamples4)=colnames(subsamples3)
subsamplesfinal10=rbind(subsamples3,subsamples4)

Drilledsp_avg=subsamplesfinal10$uniquedrilledavg.j.
Drilledsp_var=subsamplesfinal10$uniquedrilledvar.j.
DF_avg=subsamplesfinal10$DFavg.j.
DF_var=subsamplesfinal10$DFvar.j.


###########Repair frequency####################
Eventype=evenness
Location="Mc Queens pit"
a=cbind(Eventype,Location)

randomnumber=vector(mode = "integer",length=659)
raredata=array(0,dim = c(659,2))
num_species1=vector(mode="integer",length=length(unique(neuseriver2$Species)))
num_repaired=vector(mode = "integer", length =
length(unique(neuseriver2$Species)))
RF=vector(mode = "integer", length = length(unique(neuseriver2$Species)))
neuseriver_subset_2=subset(neuseriver2,select = c(Species,Repaired))
subsamps <- list()
subsamps1=list()
subsamps2=list()
for (q in 1:1000)
{
  randomnumber=sample.int(659,659, replace = FALSE)
  raredata=neuseriver_subset_2[randomnumber[1:659],]
  for(i in 1:6)
  {
    j=(i*100)
    {
      if((659-j)==59)
      {
        c=j+59
        uniquesp=unique(raredata$Species[1:c])
        num_species1[c]=length(uniquesp)
        uniquedrilled=unique(raredata$Species[which(raredata$Repaired[1:c]==1)])
        RF[c]=(sum(raredata$Repaired[1:c]))/c
        num_repaired[c]=length(uniquerepaired)
        oute=data.frame(c,num_species1[c],num_repaired[c],RF[c])
        subsamps2[[c]] <-oute
      }
      else
      {
        uniquesp=unique(raredata$Species[1:j])
        num_species1[j]=length(uniquesp)
        uniquerepaired=unique(raredata$Species[which(raredata$Repaired[1:j]==1)])
        num_repaired[j]=length(uniquerepaired)
        RF[j]=(sum(raredata$Repaired[1:j]))/j
        out=data.frame(j,num_species1[j],num_repaired[j],RF[j])
        subsamps1[[i]] <-out
```

```
      }
    }
  }
  subsamples1 <- do.call("rbind",subsamps1)
  subsamples2<-do.call("rbind",subsamps2)
  colnames(subsamples2)=colnames(subsamples1)
  subsamplesfinal11=rbind(subsamples1,subsamples2)
  out2=data.frame(q,subsamplesfinal11)
  subsamps[[q]]=out2
  subsamples <- do.call("rbind",subsamps)
}
subsamples_mcqueen_repair=as.data.frame(subsamples)
subsamples_mcqueen_repair[is.na(subsamples_mcqueen_repair)] = 0



#####################################################################################
############For calculating average and
variance#########################################################
uniquerepairedavg=vector(mode = "integer", length =
length(unique(neuseriver2$Species)))
uniquerepairedvar=vector(mode = "integer", length =
length(unique(neuseriver2$Species)))
RFavg=vector(mode = "integer", length =
length(unique(neuseriver2$Species)))
RFvar=vector(mode = "integer", length =
length(unique(neuseriver2$Species)))
subsamps1=list()
subsamps2=list()

for(i in 1:6)
{
  j=(i*100)
  {
    if((659-j)==59)
    {
      c=j+59
      uniquerepairedavg[c]=mean(subsamples_mcqueen_repair$num_repaired.j.
[which(subsamples_mcqueen_repair$j==c)])
      uniquerepairedvar[c]=sd(subsamples_mcqueen_repair$num_repaired.j.
[which(subsamples_mcqueen_repair$j==c)])
      RFavg[c]=mean(subsamples_mcqueen_repair$RF.j.
[which(subsamples_mcqueen_repair$j==c)])
      RFvar[c]=sd(subsamples_mcqueen_repair$RF.j.
[which(subsamples_mcqueen_repair$j==c)])
      oute=data.frame(c,uniquerepairedavg[c],uniquerepairedvar[c],RFavg[c],RFvar[c],a)
      subsamps2[[i]] <-oute
    }
    else
    {
      uniquerepairedavg[j]=mean(subsamples_mcqueen_repair$num_repaired.j.
[which(subsamples_mcqueen_repair$j==j)])
      uniquerepairedvar[j]=sd(subsamples_mcqueen_repair$num_repaired.j.
[which(subsamples_mcqueen_repair$j==j)])
      RFavg[j]=mean(subsamples_mcqueen_repair$RF.j.
[which(subsamples_mcqueen_repair$j==j)])
      RFvar[j]=sd(subsamples_mcqueen_repair$RF.j.
[which(subsamples_mcqueen_repair$j==j)])
```

```
      out=data.frame(j,uniquerepairedavg[j],uniquerepairedvar[j],RFavg[j],RFvar[j],a)
      subsamps1[[i]] <-out
    }
  }
}
subsamples3<- do.call("rbind",subsamps1)
subsamples4=do.call("rbind",subsamps2)
colnames(subsamples4)=colnames(subsamples3)
subsamplesfinal12=rbind(subsamples3,subsamples4)


Repairedsp_avg=subsamplesfinal12$uniquerepairedavg.j.
Repairedsp_var=subsamplesfinal12$uniquerepairedvar.j.
RF_avg=subsamplesfinal12$RFavg.j.
RF_var=subsamplesfinal12$RFvar.j.

###subsamples7 for df###
####subsamples8 for rf#######


####################Chiquita################################

neuseriver=read.csv("Chiquita.csv",header=T)
Individual=rep(1,894)
neuseriver2=data.frame(Individual,neuseriver)
wideform=as.data.frame.matrix(table(neuseriver2$Individual,neuseriver2$Species))
evenness=(diversity(wideform)/log(specnumber(wideform)))


###########evenness=0.743#########################################


Eventype=evenness
Location="Chiquita"
a=cbind(Eventype,Location)


randomnumber=vector(mode = "integer",length=894)
raredata=array(0,dim = c(894,2))
num_species1=vector(mode="integer",length=length(unique(neuseriver2$Species)))
num_drilled=vector(mode = "integer", length =
length(unique(neuseriver2$Species)))
DF=vector(mode = "integer", length = length(unique(neuseriver2$Species)))
neuseriver_subset=subset(neuseriver2,select = c(Species,Drilled))
subsamps <- list()
subsamps1=list()
subsamps2=list()
for (q in 1:1000)
{
  randomnumber=sample.int(894,894, replace = FALSE)
  raredata=neuseriver_subset[randomnumber[1:894],]
  for(i in 1:8)
  {
    j=(i*100)
    {
      if((894-j)==94)
      {
        c=j+94
```

```
        uniquesp=unique(raredata$Species[1:c])
        num_species1[c]=length(uniquesp)
        uniquedrilled=unique(raredata$Species[which(raredata$Drilled[1:c]==1)])
        DF[c]=(sum(raredata$Drilled[1:c]))/c
        num_drilled[c]=length(uniquedrilled)
        oute=data.frame(c,num_species1[c],num_drilled[c],DF[c])
        subsamps2[[c]] <-oute
      }
      else
      {
        uniquesp=unique(raredata$Species[1:j])
        num_species1[j]=length(uniquesp)
        uniquedrilled=unique(raredata$Species[which(raredata$Drilled[1:j]==1)])
        DF[j]=(sum(raredata$Drilled[1:j]))/j
        num_drilled[j]=length(uniquedrilled)
        out=data.frame(j,num_species1[j],num_drilled[j],DF[j])
        subsamps1[[i]] <-out
      }
    }
  }
  subsamples1 <- do.call("rbind",subsamps1)
  subsamples2<-do.call("rbind",subsamps2)
  colnames(subsamples2)=colnames(subsamples1)
  subsamplesfinal13=rbind(subsamples1,subsamples2)
  out2=data.frame(q,subsamplesfinal13)
  subsamps[[q]]=out2
  subsamples <- do.call("rbind",subsamps)
}
subsamples_chiquita=as.data.frame(subsamples)
subsamples_chiquita[is.na(subsamples_chiquita)] = 0



#######################################################################################
############For calculating average and
variance#########################################################
uniquedrilledavg=vector(mode = "integer", length =
length(unique(neuseriver2$Species)))
uniquedrilledvar=vector(mode = "integer", length =
length(unique(neuseriver2$Species)))
DFavg=vector(mode = "integer", length =
length(unique(neuseriver2$Species)))
DFvar=vector(mode = "integer", length =
length(unique(neuseriver2$Species)))
subsamps1=list()
subsamps2=list()
for(i in 1:8)
{
  j=(i*100)
  {
    if((894-j)==94)
    {
      c=j+94
      uniquedrilledavg[c]=mean(subsamples_chiquita$num_drilled.j.
[which(subsamples_chiquita$j==c)])
      uniquedrilledvar[c]=sd(subsamples_chiquita$num_drilled.j.
[which(subsamples_chiquita$j==c)])
```

```
      DFavg[c]=mean(subsamples_chiquita$DF.j.
[which(subsamples_chiquita$j==c)])
      DFvar[c]=sd(subsamples_chiquita$DF.j.
[which(subsamples_chiquita$j==c)])
      oute=data.frame(c,uniquedrilledavg[c],uniquedrilledvar[c],DFavg[c],DFvar[c],a)
      subsamps2[[i]] <-oute
    }
    else
    {
      uniquedrilledavg[j]=mean(subsamples_chiquita$num_drilled.j.
[which(subsamples_chiquita$j==j)])
      uniquedrilledvar[j]=sd(subsamples_chiquita$num_drilled.j.
[which(subsamples_chiquita$j==j)])
      DFavg[j]=mean(subsamples_chiquita$DF.j.
[which(subsamples_chiquita$j==j)])
      DFvar[j]=sd(subsamples_chiquita$DF.j.
[which(subsamples_chiquita$j==j)])
      out=data.frame(j,uniquedrilledavg[j],uniquedrilledvar[j],DFavg[j],DFvar[j],a)
      subsamps1[[i]] <-out
    }
  }
}
subsamples3<- do.call("rbind",subsamps1)
subsamples4=do.call("rbind",subsamps2)
colnames(subsamples4)=colnames(subsamples3)
subsamplesfinal14=rbind(subsamples3,subsamples4)

Drilledsp_avg=subsamplesfinal14$uniquedrilledavg.j.
Drilledsp_var=subsamplesfinal14$uniquedrilledvar.j.
DF_avg=subsamplesfinal14$DFavg.j.
DF_var=subsamplesfinal14$DFvar.j.


###########Repair frequency###################
Eventype=evenness
Location="Chiquita"
a=cbind(Eventype,Location)

randomnumber=vector(mode = "integer",length=894)
raredata=array(0,dim = c(894,2))
num_species1=vector(mode="integer",length=length(unique(neuseriver2$Species)))
num_repaired=vector(mode = "integer", length =
length(unique(neuseriver2$Species)))
RF=vector(mode = "integer", length = length(unique(neuseriver2$Species)))
neuseriver_subset_2=subset(neuseriver2,select = c(Species,Repaired))
subsamps <- list()
subsamps1=list()
subsamps2=list()
for (q in 1:1000)
{
  randomnumber=sample.int(894,894, replace = FALSE)
  raredata=neuseriver_subset_2[randomnumber[1:894],]
  for(i in 1:8)
  {
    j=(i*100)
    {
      if((894-j)==94)
      {
```

```
        c=j+94
        uniquesp=unique(raredata$Species[1:c])
        num_species1[c]=length(uniquesp)
        uniquedrilled=unique(raredata$Species[which(raredata$Repaired[1:c]==1)])
        RF[c]=(sum(raredata$Repaired[1:c]))/c
        num_repaired[c]=length(uniquerepaired)
        oute=data.frame(c,num_species1[c],num_repaired[c],RF[c])
        subsamps2[[c]] <-oute
      }
      else
      {
        uniquesp=unique(raredata$Species[1:j])
        num_species1[j]=length(uniquesp)
        uniquerepaired=unique(raredata$Species[which(raredata$Repaired[1:j]==1)])
        num_repaired[j]=length(uniquerepaired)
        RF[j]=(sum(raredata$Repaired[1:j]))/j
        out=data.frame(j,num_species1[j],num_repaired[j],RF[j])
        subsamps1[[i]] <-out
      }
    }
  }
  subsamples1 <- do.call("rbind",subsamps1)
  subsamples2<-do.call("rbind",subsamps2)
  colnames(subsamples2)=colnames(subsamples1)
  subsamplesfinal15=rbind(subsamples1,subsamples2)
  out2=data.frame(q,subsamplesfinal15)
  subsamps[[q]]=out2
  subsamples <- do.call("rbind",subsamps)
}
subsamples_chiquita_repair=as.data.frame(subsamples)
subsamples_chiquita_repair[is.na(subsamples_chiquita_repair)] = 0


#####################################################################################
############For calculating average and 
variance#########################################################
uniquerepairedavg=vector(mode = "integer", length = 
length(unique(neuseriver2$Species)))
uniquerepairedvar=vector(mode = "integer", length = 
length(unique(neuseriver2$Species)))
RFavg=vector(mode = "integer", length = 
length(unique(neuseriver2$Species)))
RFvar=vector(mode = "integer", length = 
length(unique(neuseriver2$Species)))
subsamps1=list()
subsamps2=list()
for(i in 1:8)
{
  j=(i*100)
  {
    if((894-j)==94)
    {
      c=j+94
      uniquerepairedavg[c]=mean(subsamples_chiquita_repair$num_repaired.j.
[which(subsamples_chiquita_repair$j==c)])
      uniquerepairedvar[c]=sd(subsamples_chiquita_repair$num_repaired.j.
[which(subsamples_chiquita_repair$j==c)])
```

```
      RFavg[c]=mean(subsamples_chiquita_repair$RF.j.
[which(subsamples_chiquita_repair$j==c)])
      RFvar[c]=sd(subsamples_chiquita_repair$RF.j.
[which(subsamples_chiquita_repair$j==c)])
      oute=data.frame(c,uniquerepairedavg[c],uniquerepairedvar[c],RFavg[c],RFvar[c],a)
      subsamps2[[i]] <-oute
    }
    else
    {
      uniquerepairedavg[j]=mean(subsamples_chiquita_repair$num_repaired.j.
[which(subsamples_chiquita_repair$j==j)])
      uniquerepairedvar[j]=sd(subsamples_chiquita_repair$num_repaired.j.
[which(subsamples_chiquita_repair$j==j)])
      RFavg[j]=mean(subsamples_chiquita_repair$RF.j.
[which(subsamples_chiquita_repair$j==j)])
      RFvar[j]=sd(subsamples_chiquita_repair$RF.j.
[which(subsamples_chiquita_repair$j==j)])
      out=data.frame(j,uniquerepairedavg[j],uniquerepairedvar[j],RFavg[j],RFvar[j],a)
      subsamps1[[i]] <-out
    }
  }
}

subsamples3<- do.call("rbind",subsamps1)
subsamples4=do.call("rbind",subsamps2)
colnames(subsamples4)=colnames(subsamples3)
subsamplesfinal16=rbind(subsamples3,subsamples4)

Repairedsp_avg=subsamplesfinal16$uniquerepairedavg.j.
Repairedsp_var=subsamplesfinal16$uniquerepairedvar.j.
RF_avg=subsamplesfinal16$RFavg.j.
RF_var=subsamplesfinal16$RFvar.j.

###subsamples11 for df###
####subsamples12 for rf#######

##############combining the plots#################################
combineddata_df=rbind(subsamplesfinal1,subsamplesfinal6,subsamplesfinal10,subsamplesfina
combineddata_rf=rbind(subsamplesfinal4,subsamplesfinal8,subsamplesfinal12,subsamplesfina


#write.table(combineddata_df,file="combined_df_ex_neuse n punta1.csv")
#write.table(combineddata_rf,file="combined_rf_ex_neuse n punta1.csv")

combinedata_df_1=read.csv(file="combined_df_ex_neuse n
punta1.csv",header=T)
combinedata_rf_1=read.csv(file="combined_rf_ex_neuse n
punta1.csv",header=T)

require(dplyr)
require(ggplot2)

#################drilling predation#####################
colorscale=c("dodgerblue4","skyblue2","orangered4","salmon2","green","pink")
Drilledsp_avg=combinedata_df_1$uniquedrilledavg.j.
Drilledsp_var=combinedata_df_1$uniquedrilledvar.j.
DF_avg=combinedata_df_1$DFavg.j.
DF_var=combinedata_df_1$DFvar.j.
```

```
j=combinedata_df_1$j
Eventype=combinedata_df_1$Eventype
Location=combinedata_df_1$Location
Evenness=factor(round(Eventype, 2))
Location=factor(Location)
df1=ggplot(combinedata_df_1,aes(j,DF_avg,gcoroup=Evenness,color=Evenness))+
xlab("Sample size") + ylab("DF")+scale_colour_manual(values =
c("dodgerblue4","skyblue2","salmon2","orangered4"))+geom_point(size = 1)
+geom_errorbar(aes(ymin = DF_avg-DF_var, ymax = DF_avg+DF_var))
+theme(legend.position="none")
#facet_grid(vars(Case), vars(DFtype))+ theme(strip.text.x =
element_text(size = 12, face = "bold"),strip.text.y = element_text(size =
12, face = "bold" ))+theme(axis.title =  element_text(size = 15))
+theme(axis.text =  element_text(size = 10))+theme(aspect.ratio=1)
df2=ggplot(combinedata_df_1,aes(j,Drilledsp_avg,color=Evenness)) +
xlab("Sample size") + ylab("Drilled species")+scale_colour_manual(values =
c("dodgerblue4","skyblue2","salmon2","orangered4"))+geom_point(size = 1)
+geom_errorbar(aes(ymin = Drilledsp_avg-Drilledsp_var, ymax =
Drilledsp_avg+Drilledsp_var))+theme(legend.position="none")
#facet_grid(vars(Case),vars(DFtype))+ theme(strip.text.x =
element_text(size = 12, face = "bold"),strip.text.y = element_text(size =
12, face = "bold" ))+theme(axis.title =  element_text(size = 15))
+theme(axis.text =  element_text(size = 10))+theme(aspect.ratio=1)
df3=ggplot(combinedata_df_1,aes(Drilledsp_avg,DF_avg,color=Evenness)) +
ylab("DF") + xlab("Drilled species")+scale_colour_manual(values =
c("dodgerblue4","skyblue2","salmon2","orangered4","green"))+geom_point(size
= 1)+ geom_errorbar(aes(ymin = DF_avg-DF_var, ymax = DF_avg+DF_var))
+geom_errorbarh(aes(xmin = Drilledsp_avg-Drilledsp_var, xmax =
Drilledsp_avg+Drilledsp_var))+theme(legend.position="none")
#facet_grid(vars(Case),vars(DFtype))+theme(strip.text.x = element_text(size
= 12, face = "bold"),strip.text.y = element_text(size = 12, face = "bold"
))+theme(axis.title =  element_text(size = 15))+theme(axis.text =
element_text(size = 10))+theme(aspect.ratio=1)

####repair frequency############
require(dplyr)
require(ggplot2)
colorscale=c("dodgerblue4","skyblue2","orangered4","salmon2","green","pink")
Repairedsp_avg=combinedata_rf_1$uniquerepairedavg.j.
Repairedsp_var=combinedata_rf_1$uniquerepairedvar.j.
RF_avg=combinedata_rf_1$RFavg.j.
RF_var=combinedata_rf_1$RFvar.j.
j=combinedata_rf_1$j
Eventype=combinedata_rf_1$Eventype
Location=combinedata_rf_1$Location
Evenness=factor(round(Eventype, 2))
Location=factor(Location)
rf1=ggplot(combinedata_rf_1,aes(j,RF_avg,gcoroup=Evenness,color=Evenness))+
xlab("Sample size") + ylab("RF")+scale_colour_manual(values =
c("dodgerblue4","skyblue2","salmon2","orangered4","green","pink"))
+geom_point(size = 1)+geom_errorbar(aes(ymin = RF_avg-RF_var, ymax =
RF_avg+RF_var))+theme(legend.position="none")
#facet_grid(vars(Case), vars(DFtype))+ theme(strip.text.x =
element_text(size = 12, face = "bold"),strip.text.y = element_text(size =
12, face = "bold" ))+theme(axis.title =  element_text(size = 15))
+theme(axis.text =  element_text(size = 10))+theme(aspect.ratio=1)
```

```
rf2=ggplot(combinedata_rf_1,aes(j,Repairedsp_avg,color=Evenness)) + 
xlab("Sample size") + ylab("Repaired species")+scale_colour_manual(values = 
c("dodgerblue4","skyblue2","salmon2","orangered4","green","pink"))
+geom_point(size = 1)+geom_errorbar(aes(ymin = Repairedsp_avg-
Repairedsp_var, ymax = Repairedsp_avg+Repairedsp_var))
+theme(legend.position="none")
#facet_grid(vars(Case),vars(DFtype))+ theme(strip.text.x = 
element_text(size = 12, face = "bold"),strip.text.y = element_text(size = 
12, face = "bold" ))+theme(axis.title =  element_text(size = 15))
+theme(axis.text =  element_text(size = 10))+theme(aspect.ratio=1)
rf3=ggplot(combinedata_rf_1,aes(Repairedsp_avg,RF_avg,color=Evenness)) + 
ylab("RF") + xlab("Repaired species")+scale_colour_manual(values = 
c("dodgerblue4","skyblue2","salmon2","orangered4","green","pink"))
+geom_point(size = 1)+ geom_errorbar(aes(ymin = RF_avg-RF_var, ymax = 
RF_avg+RF_var))+geom_errorbarh(aes(xmin = Repairedsp_avg-Repairedsp_var, 
xmax =Repairedsp_avg+Repairedsp_var))+theme(legend.position="bottom")
#facet_grid(vars(Case),vars(DFtype))+theme(strip.text.x = element_text(size 
= 12, face = "bold"),strip.text.y = element_text(size = 12, face = "bold" 
))+theme(axis.title =  element_text(size = 15))+theme(axis.text = 
element_text(size = 10))+theme(aspect.ratio=1)


require(gridExtra)

grid.arrange(arrangeGrob(df1, rf1,df2,rf2,df3,rf3,ncol = 2), nrow = 1)
# First row with one plot spaning over 2 columns


shared_legend <- function(...) {



  plots <- list(...)



  g <- ggplotGrob(plots[[1]] + theme(legend.position="bottom"))$grobs



  legend <- g[[which(sapply(g, function(x) x$name) == "guide-box")]]



  lheight <- sum(legend$height)



  grid.arrange(

    do.call(arrangeGrob, lapply(plots, function(x)

      x + theme(legend.position="none"))),

    legend,

    ncol = 1,
```

```
    heights = unit.c(unit(1, "npc") - lheight, lheight))

}
library(grid)
shared_legend(df1,rf1,df2,rf2,df3,rf3)

######Kolmogorov smironov test for PI and S values at j=500 between all
four locations############################
####################Drilling
frequency########################################
subsamples_chiquita_df=subset(subsamples_chiquita,subsamples_chiquita$j==500,select=c(D
subsamples_mcqueen_df=subset(subsamples_mcqueen,subsamples_mcqueen$j==500,select=c(DF.j
subsamples_punta_df=subset(subsamples_punta,subsamples_punta$j==500,select=c(DF.j.))
subsamples_miami_df=subset(subsamples_miami,subsamples_miami$j==500,select=c(DF.j.))
ks.test(subsamples_chiquita_df$DF.j.,subsamples_mcqueen_df$DF.j.)
ks.test(subsamples_chiquita_df$DF.j.,subsamples_miami_df$DF.j.)
ks.test(subsamples_chiquita_df$DF.j.,subsamples_punta_df$DF.j.)
ks.test(subsamples_mcqueen_df$DF.j.,subsamples_punta_df$DF.j.)
ks.test(subsamples_miami_df$DF.j.,subsamples_punta_df$DF.j.)
ks.test(subsamples_miami_df$DF.j.,subsamples_mcqueen_df$DF.j.)

####################Drilled species######################################
subsamples_chiquita_s=subset(subsamples_chiquita,subsamples_chiquita$j==500,select=c(nu
subsamples_mcqueen_s=subset(subsamples_mcqueen,subsamples_mcqueen$j==500,select=c(num_d
subsamples_punta_s=subset(subsamples_punta,subsamples_punta$j==500,select=c(num_drilled
subsamples_miami_s=subset(subsamples_miami,subsamples_miami$j==500,select=c(num_drilled
ks.test(subsamples_chiquita_s$num_drilled.j.,subsamples_mcqueen_s$num_drilled.j.)
ks.test(subsamples_chiquita_s$num_drilled.j.,subsamples_miami_s$num_drilled.j.)
ks.test(subsamples_chiquita_s$num_drilled.j.,subsamples_punta_s$num_drilled.j.)
ks.test(subsamples_mcqueen_s$num_drilled.j.,subsamples_punta_s$num_drilled.j.)
ks.test(subsamples_miami_s$num_drilled.j.,subsamples_punta_s$num_drilled.j.)
ks.test(subsamples_miami_s$num_drilled.j.,subsamples_mcqueen_s$num_drilled.j.)

#######################Repair species#####################################
subsamples_chiquita_repair_s=subset(subsamples_chiquita_repair,subsamples_chiquita_repa
subsamples_mcqueen_repair_s=subset(subsamples_mcqueen_repair,subsamples_mcqueen_repair$
subsamples_punta_repair_s=subset(subsamples_punta_repair,subsamples_punta_repair$j==500
subsamples_miami_repair_s=subset(subsamples_miami_repair,subsamples_miami_repair$j==500
ks.test(subsamples_chiquita_repair_s$num_repaired.j.,subsamples_mcqueen_repair_s$num_re
ks.test(subsamples_chiquita_repair_s$num_repaired.j.,subsamples_miami_repair_s$num_repa
ks.test(subsamples_chiquita_repair_s$num_repaired.j.,subsamples_punta_repair_s$num_repa
ks.test(subsamples_mcqueen_repair_s$num_repaired.j.,subsamples_punta_repair_s$num_repai
ks.test(subsamples_miami_repair_s$num_repaired.j.,subsamples_punta_repair_s$num_repaire
ks.test(subsamples_miami_repair_s$num_repaired.j.,subsamples_mcqueen_repair_s$num_repai

#######################Repair
frequency#####################################
subsamples_chiquita_repair_rf=subset(subsamples_chiquita_repair,subsamples_chiquita_rep
subsamples_mcqueen_repair_rf=subset(subsamples_mcqueen_repair,subsamples_mcqueen_repair
subsamples_punta_repair_rf=subset(subsamples_punta_repair,subsamples_punta_repair$j==50
subsamples_miami_repair_rf=subset(subsamples_miami_repair,subsamples_miami_repair$j==50
ks.test(subsamples_chiquita_repair_rf$RF.j.,subsamples_mcqueen_repair_rf$RF.j.)
ks.test(subsamples_chiquita_repair_rf$RF.j.,subsamples_miami_repair_rf$RF.j.)
ks.test(subsamples_chiquita_repair_rf$RF.j.,subsamples_punta_repair_rf$RF.j.)
ks.test(subsamples_mcqueen_repair_rf$RF.j.,subsamples_punta_repair_rf$RF.j.)
ks.test(subsamples_miami_repair_rf$RF.j.,subsamples_punta_repair_rf$RF.j.)
ks.test(subsamples_miami_repair_rf$RF.j.,subsamples_mcqueen_repair_rf$RF.j.)
```

```
########################Abundance vs Pi
PLOT######################################################################
casestudy=read.csv("abundance vs pi.csv", header = T)
locations=as.factor(casestudy[,1])
ggplot(casestudy,aes(casestudy$PI.prey.drilling,casestudy$Proportional.abundance))
+geom_line(aes(colour=locations))+geom_point(aes(colour=locations))
ggplot(casestudy,aes(casestudy$PI.prey.durophagy,casestudy$Proportional.abundance))
+geom_line(aes(colour=locations))+geom_point(aes(colour=locations))

chiquita=subset(casestudy,casestudy$X=="Chiquita")
cor.test(chiquita$Proportional.abundance,chiquita$PI.prey.drilling,method =
"spearman")
cor.test(chiquita$Proportional.abundance,chiquita$PI.prey.durophagy,method
= "spearman")
Punta=subset(casestudy,casestudy$X=="Punta Gorda")
cor.test(Punta$Proportional.abundance,Punta$PI.prey.drilling,method =
"spearman")
cor.test(Punta$Proportional.abundance,Punta$PI.prey.durophagy,method =
"spearman")
mcqueen=subset(casestudy,casestudy$X=="Mc Queens pit")
cor.test(mcqueen$Proportional.abundance,mcqueen$PI.prey.drilling,method =
"spearman")
cor.test(mcqueen$Proportional.abundance,mcqueen$PI.prey.durophagy,method =
"spearman")
miami=subset(casestudy,casestudy$X=="Miami Canal")
cor.test(miami$Proportional.abundance,miami$PI.prey.drilling,method =
"spearman")
cor.test(miami$Proportional.abundance,miami$PI.prey.durophagy,method =
"spearman")
```
